# MFSD1 in complex with its accessory subunit GLMP functions as a general dipeptide uniporter in lysosomes

**DOI:** 10.1101/2023.12.15.570541

**Authors:** Katharina Esther Julia Jungnickel, Océane Guelle, Miharu Iguchi, Wentao Dong, Vadim Kotov, Florian Gabriel, Cécile Debacker, Julien Dairou, Isabelle McCort-Tranchepain, Nouf N. Laqtom, Sze Ham Chan, Akika Ejima, Kenji Sato, David Massa López, Paul Saftig, Ahmad Reza Mehdipour, Monther Abu-Remaileh, Bruno Gasnier, Christian Löw, Markus Damme

**Affiliations:** Centre for Structural Systems Biology (CSSB), Notkestraße 85, 22607 Hamburg, Germany; 2European Molecular Biology Laboratory (EMBL) Hamburg, Notkestraße 85, 22607 Hamburg, Germany; Saints-Pères Paris Institute for the Neurosciences, Université de Paris, Centre National de la Recherche Scientifique, F-75006 Paris, France; Department of Chemical Engineering, Stanford University, Stanford, CA 94305; Department of Genetics, Stanford University, Stanford, CA 94305; The Institute for Chemistry, Engineering & Medicine for Human Health, Stanford University, Stanford, CA 94305; Laboratoire de Chimie et Biochimie Pharmacologiques et Toxicologiques, CNRS UMR 8601, Université Paris Cité, F-75006 Paris, France; Department of Pharmacology, University of Virginia School of Medicine, Charlottesville, VA, USA; Graduate School of Agriculture, Kyoto University, Kyoto 606 8502, Japan; Institute of Biochemistry, Christian-Albrechts-University Kiel, Kiel, Germany; UGent Center for Molecular Modeling, Ghent University, Ghent, Belgium

**Keywords:** Lysosomal transporter, Uniporter, Dipeptides, MFSD1, GLMP, Lysosomes, Cryo-EM structure

## Abstract

Lysosomal degradation of macromolecules in lysosomes produces diverse small metabolites exported by specific transporters for reuse in biosynthetic pathways. Here, we deorphanized the Major Facilitator Superfamily Domain Containing 1 (MFSD1) protein, which forms a tight complex with the Glycosylated Lysosomal Membrane Protein (GLMP) in the lysosomal membrane. Untargeted metabolomics analysis of MFSD1-deficient mouse lysosomes revealed an increase in cationic dipeptides. Purified MFSD1 selectively bound diverse dipeptides, while electrophysiological, isotope tracer, and fluorescence-based studies in *Xenopus* oocytes and proteoliposomes showed that MFSD1/GLMP acts as a uniporter for cationic and neutral dipeptides. Cryo-EM structure of the dipeptide-bound MFSD1/GLMP complex in outward-open conformation characterized the heterodimer interface and, in combination with molecular dynamics simulations, provided a structural basis for its selectivity towards diverse dipeptides. Together, our data identify MFSD1 as a general lysosomal dipeptide uniporter, providing an alternative route to recycle lysosomal proteolysis products when lysosomal amino acid exporters are overloaded.

## INTRODUCTION

One of the major functions of lysosomes is the hydrolytic degradation of various macromolecules, including complex lipids, oligosaccharides, and nucleic acids. Another central catabolic function, however, is their critical role in maintaining cellular proteostasis via the hydrolytic degradation of extracellular and intracellular proteins reaching lysosomes by endocytosis, phagocytosis (from the extracellular compartment), or autophagy (intracellular proteins and organelles) facilitated by various lysosomal proteases (Ballabio and Bonifacino, 2020; Settembre and Perera, 2023). Bulk proteolysis is important during homeostatic conditions to prevent the buildup of undegraded proteins in lysosomes but is especially critical under conditions of starvation for amino acid recycling or for the removal of aggregated proteins. Indeed, the lysosomal pool of free amino acids critically governs amino acid sensing and synthesis of proteins (Wolfson and Sabatini, 2017). A set of ∼15 relatively promiscuous lysosomal proteases mediates the hydrolysis of various peptide bonds between most amino acids, yielding short peptides and free amino acids (Winchester, 2005). Specific transport systems for individual amino acids or different classes, such as hydrophobic, negatively- or positively charged, bulky amino acids, and short peptides, can facilitate their export from the lysosomal lumen to the cytoplasm (Jezegou et al., 2012; Kalatzis et al., 2001; Lloyd, 1996; Sakata et al., 2001; Verdon et al., 2017; Wyant et al., 2017). While many lysosomal proteases possess both exo- and endopeptidase activity with a relatively low substrate specificity, some lysosomal proteases are specific dipeptidyl and tripeptidyl peptidases cleaving two or three amino acids from the C- and N-termini. Notably, the bulk of intracellular dipeptidase activity is localized in the cytosol (Thamotharan et al., 1997), and in fact, it is unclear if any nonspecific lysosomal dipeptidase exists (Lloyd, 1996; Winchester, 2005). Moreover, it was already noted in the 1970s that several peptides are resistant to lysosomal proteolysis, including peptides containing proline and hydroxyproline (Coffey et al., 1976). These data indicate that dipeptides are among the end products of the lysosomal digestion of proteins that require exportation to the cytoplasm (Coffey and De Duve, 1968; Lloyd, 1996).

ParaParaTo characterize orphan lysosomal transporters, we started to investigate Major facilitator superfamily domain containing 1 (MFSD1) transporter, which we and others found by mass spectrometry in isolated lysosomes (Chapel et al., 2013; Markmann et al., 2017). MFSD1 belongs to the Major Facilitator Superfamily (MFS) of transporters, but its substrate(s) are unknown (Ferrada and Superti-Furga, 2022). Members of this superfamily are comprised of

12 transmembrane (TMs) helices organized into two pseudosymmetric six-helix bundles called the N-terminal domain (N-domain, TMs 1-6) and the C-terminal domain (C-domain, TMs 7-12) (Drew et al., 2021; Reddy et al., 2012). They typically mediate the import/export of water-soluble molecules such as sugars, nucleosides, monocarboxylates, or peptides following a rocker-switch mechanism (Bartels et al., 2021; Drew et al., 2021; Law et al., 2008; Quistgaard et al., 2016).

MFSD1 is ubiquitously expressed in mouse tissues (Massa Lopez et al., 2019). We previously validated its lysosomal localization upon ectopic expression in cultured cells and at the endogenous level (Massa Lopez et al., 2019). In contrast to the majority of integral lysosomal transmembrane proteins, MFSD1 is not N-glycosylated (Massa Lopez et al., 2019). We found that MFSD1 forms a tight heterodimeric complex with another poorly characterized lysosomal membrane protein: Glycosylated lysosomal membrane protein (GLMP) (Massa Lopez et al., 2019), a single pass type I transmembrane protein that is extensively N-glycosylated. MFSD1 and GLMP form a tight complex, and in the absence of one of the proteins, the other partner is rapidly degraded in lysates of cells or tissues, proposing a chaperone function and protective effect of the two proteins towards lysosomal proteases (Massa Lopez et al., 2019). The remaining MFSD1 is quantitatively retained in the Golgi-apparatus in GLMP-deficient cells, indicating an additional role of the two interacting partners in the transport of the complex from the Golgi-apparatus to lysosomes (Lopez et al., 2020).

To identify the substrate(s) of MFSD1, we performed untargeted metabolomic analyses of liver lysosomes from *Mfsd1* knockout mice and identified different dipeptides to be drastically enriched in these lysosomes, making them prime candidates as putative substrates. We validated the transport of dipeptides *in cellula* in oocytes upon expression of MFSD1 together with GLMP by means of electrophysiology and mass spectrometry, observed direct binding of dipeptides to recombinantly expressed and purified MFSD1, and confirmed the transport in a reconstituted *in vitro* system. The structure of the MFSD1-GLMP complex was determined by cryo-electron microscopy (cryo-EM) in its apo-form and in the presence of a dipeptide. Together with molecular dynamics (MD) simulations, we obtained a detailed molecular picture of how lysosomal dipeptides are recognized and transported. By combining cell biology, advanced biochemical transport assays, structural biology, and MD simulations, we have deorphanized MFSD1 as a novel lysosomal dipeptide uniporter.

## MATERIAL & METHODS

Peptides were purchased from Bachem or Sigma Aldrich. All amino acids used belong to the L series. Most charged peptides were obtained as salts with the following counterions: hydrochloride (Ala-Lys; Lys-Pro; Lys-Val); hydrobromide (Lys-Ala); acetate (Arg-Ala; Lys-Ala-Ala; Pro-Arg); nitrate (anserine); Chemicals and reagents were purchased, if not otherwise indicated, from Sigma-Aldrich. A complete list of peptides is depicted in **Supplemental Table 1**. Hydroxyproline-bound 2-chlorotrityl chloride (Barlos) resin and *N*-α-(9- Fluorenylmethoxycarbonyl (Fmoc)-*N*-ω-(4-methoxy-2,3,6-trimethylbenzenesulfonyl)-L-arginine were obtained from Watanabe Chemicals (Hiroshima, Japan). 6-aminoquinolyl-*N*- hydroxysuccinimidyl carbamate (AccQ) was purchased from Toronto Research Chemicals, Toronto, Canada). Arginyl-hydroxyproline (Arg-Hyp) was synthesized according to the Fmoc strategy using a PSSM-8 peptide synthesizer (Shimadzu, Kyoto, Japan). Synthesized Arg- Hyp was purified by reversed-phase HPLC using a Cosmosil 5C18-MS-II column (10 mm × 250 mm, Nacalai Tesque, Kyoto, Japan). A binary gradient was used with 0.1% formic acid (solvent A) and 0.1% formic acid containing 80% acetonitrile (solvent B) at a flow rate of 2.0 mL/min. The chemicals for leucine-5,5,5-*d_3_*-alanine synthesis, *tert*-butoxycarbonyl-Leucine- 5,5,5-*d*_3_ (98%), HCl.Alanine-O*t*Bu (99%), (Benzotriazol-1-yloxy)tripyrrolidinophosphonium hexafluorophosphate (PyBOP) were purchased from Cambridge Isotope Laboratories, Sigma, and Novabiochem, respectively. The leucine-5,5,5-*d*_3_ standard was from Cambridge Isotope Laboratories. BAY-069 was from MedChemExpress.

### Synthesis of Leucine-5,5,5-d_3_-alanine

Dipeptide leucine-5,5,5-*d_3_*-alanine (Leu(*d_3_*)-Ala) hydrochloride, as a mixture of 2 diastereoisomers, was synthesized in 2 steps by coupling Boc-Leu-5,5,5-*d_3_*-OH with HCl.Ala- O*t*Bu using PyBOP as the coupling reagent (Nasief and Hangauer, 2014), followed by the deprotection of the protecting groups in acidic conditions, as shown in the following scheme:

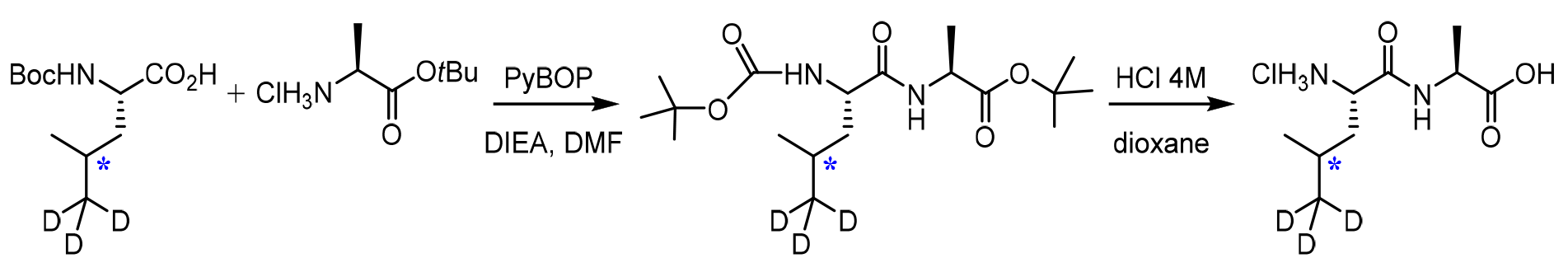

### General synthesis protocol for Leu-5,5,5-*d*_3_-Ala

All reactions were carried out under an argon atmosphere with anhydrous solvent and were monitored by thin layer chromatography (TLC) with silica gel Merck 60 F254 on aluminum sheets. Automated flash chromatography was performed with a Biotage apparatus with evaporative light scattering detection and UV detectors using a Büchi FlashPure silica column. Solvent systems were given according to (s/s: v/v). ^1^H (500.16 MHz), ^13^C (125.78 MHz), and nuclear magnetic resonance (NMR) spectra were recorded on a 500 Bruker spectrometer equipped with a sensitivity-optimized measurement head (cryoprobe). Chemical shifts (δ, ppm) are given with reference to deuterated solvents for ^1^H and ^13^C NMR, respectively, CDCl_3_: 7.24, 77.23; D_2_O: 4.78. Signal multiplicity is described as follows: s (singlet), d (doublet), t (triplet), q (quartet), quin (quintuplet), m (multiplet). Broad signals are described as br. Coupling constants (*J*) are given in Hz. Greek letters are used as locants for NMR attributions, which were established on the basis of ^13^C using ^1^H decoupled spectra as well as COSY, HSQC, and HMBC.

### Synthesis of *tert*-Butyl (*tert*-butoxycarbonyl)-leucyl-5,5,5-*d_3_*-alaninate

To a cooled solution of Boc-leucine-5,5,5-*d*_3_ (469.0 mg, 2.0 mmol, 1.0 eq), HCl.alanine-O*t*Bu (550.50 mg, 3.0 mmol, 1.5 eq), (benzotriazol-1-yloxy)tripyrrolidinophosphonium hexafluorophosphate (PyBOP) (1.25 g, 2.4 mmol, 1.2 eq) in dimethylformamide (9.6 mL), *N*,*N*-diisopropylethylamine (DIEA) (1.4 mL, 8.0 mmol, 4 eq) was added slowly. The reaction mixture was stirred at room temperature overnight, diluted with EtOAc (10 mL for 1 mL of DMF), and then extracted with a cooled solution of 5% aqueous KHSO_4_ (2x), saturated NaHCO_3_ (2x) and brine (2x). The organic layer was then dried with Na_2_SO_4_, filtered, and evaporated under vacuum to give the product after purification by flash chromatography (Cyclohexane/EtOAc: 90/10) as a white solid in 51% yield (370 mg, 1.02 mmol). ^1^H NMR in CDCl_3_ showed the presence of two rotamers due to the Boc group (80/20).

^1^H NMR (500 MHz, CDCl_3_) *δ*: 7.06 (d, *J* = 6.0 Hz, 0.8H, NH-Ala), 6.74 (brs, 0.2H, NH-Ala), 5.69 (brs, 0.2H, NH-Boc), 5.32 (d, *J* = 8.5 Hz, 0.8H, NH-Boc), 4.28 (quin, *J* = 7.0 Hz, 1H, Hα-Ala), 4.12 (m, 0.8 H, Hα-Leu), 3.89 (brs, 0.2H, Hα-Leu), 1.59 (m, 1H, Hγ-Leu), 1.55─1.38 (m, 2H, Hβ-Leu), 1.34 (s, 9H, CO_2_*t*Bu), 1.31, 1.30 (2s, 9H, Boc), 1.22 (d, *J* = 7.5 Hz, 3H, Hβ-Ala), 0.82, 0.80 (2d, *J* = 6.0 Hz, 3H, Hδ-Leu); ^13^C NMR (500 MHz, CDCl_3_) *δ*: 172.6, 172.5 (CONH), 171.9 (CO_2_*t*Bu), 155.9 (CO-NHBoc), 81.5, 81.4 (Cq-NHBoc), 79.5 (Cq-CO_2_*t*Bu), 52.9 (Cα-Leu), 48.6 (Cα-Ala), 41.5 (Cβ-Leu), 28.3, 27.9 (CH_3_-*t*Bu), 24.4 (Cγ-Leu), 23.0, 21.8 (Cδ-Leu), 18.0, 17.9 (Cβ-Ala).

### Synthesis of leucyl-5,5,5-*d_3_*-alanine hydrochloride: LSP11-280723

To a solution of Boc-Leu-5,5,5-*d*_3_-Ala-O*t*Bu (120.0 mg, 0.33 mmol) in dioxane (0.25 mL) at 0 °C was added slowly a solution of HCl 4 M in dioxane (2.5 mL). After 30 min at this temperature, the reaction mixture was stirred at room temperature overnight. Evaporation of the solvent under vacuum and recrystallization with MeOH─Et_2_O afforded HCl.leucyl-5,5,5- *d_3_*-alanine as a white solid (66.5 mg, 0.275 mmol) in 83% yield.

^1^H NM (500 MHz, D_2_O) *δ*: 4.36 (q, *J* = 7.0 Hz, 1H, Hα-Ala), 3.93 (t, *J* = 7.0 Hz, 1H, Hα-Leu), 1.77-1.61 (m, 3H, Hγ-Leu, Hβ-Leu), 1.40 (d, *J* = 7.0 Hz, 3H, Hβ-Ala), 0.92, 0.90 (2d, *J* = 6.0 Hz, 3H, Hδ-Leu), ; ^13^C NMR (500 MHz, D_2_O) *δ*: 176.0 (CO_2_H), 170.1 (CONH), 51.6 (Cα-Leu), 48.8 (Cα-Ala), 39.7 (Cβ-Leu), 23.4 (Cγ-Leu), 21.5, 20.9 (Cδ-Leu), 15.9 (Cβ-Ala).

### Cell lines, mouse strains, antibodies

*Mfsd1* knockout mice were described previously (Massa Lopez et al., 2019). HeLa cells were purchased from CLS Cell Lines Service and used at low passage numbers. Expi293F cells were purchased from Thermo Fisher (Cat. number: A14527). Mouse embryonic fibroblasts (MEF) from *Glmp* knockout mice were described previously (Massa Lopez et al., 2019).

Antibodies used throughout the study: LAMP1 clone 1D4B (rat monoclonal, Developmental Studies Hybridoma Bank); LAMP1 clone 1D4B (rat monoclonal, conjugated to AlexaFluor 647, BioLegend); HA clone 3F10 (rat monoclonal, Sigma-Aldrich / Merck),); HA clone 3F10 (rat monoclonal, conjugated to FITC, Sigma-Aldrich / Merck), GFP (mouse monoclonal, Roche Molecular Biochemicals), mKate2 (rabbit polyclonal, Origene), KDEL (mouse monoclonal, Enzo Life Sciences), Cox IV (rabbit polyclonal, ab16056, Abcam), Golgin 97 (clone CDF4, mouse monoclonal, Thermo Scientific Fisher). The antibody against cathepsin D (CTSD) was custom-made against a synthetic peptide (CKSDQSKARGIKVEKQIFGEATKQP) and immunization of rabbits, followed by affinity purification against the immunization peptide. The custom-made MFSD1- and GLMP-specific antibodies were described before (Massa Lopez et al., 2019).

### Cell culture and transfection of eukaryotic cells

For transfection of Hela cells, 1–5 μg of DNA were incubated with polyethylenimine (PEI) in DMEM (without antibiotics nor FBS) for 15 min at room temperature. The mix was applied to the culture of cells, and after ∼6 hours, the media was exchanged. The transfected cells were analyzed 48 hours post-transfection.

### Cloning of cDNA constructs for oocyte expression

Lysosomal sorting motif mutations, Y400A and L11A/L12A were introduced into mouse GLMP and MFSD1 plasmids, respectively, using Q5® Site-Directed Mutagenesis Kit (New England Biolabs). The whole coding sequence was verified by automated sequencing. mGLMP_Y400A_-mKate2 and mMFSD1_L11A/L12AA_-EmGFP cDNAs were then subcloned into the pOX(+) vector for *Xenopus* oocyte expression. In this vector, the cDNA of interest is flanked by the 5′- and 3′-noncoding sequences from *Xenopus laevis* β-globulin mRNA to increase expression.

### Cloning, expression, and purification of MFSD1, GLMP and GLMP-MFSD1-fusion protein for recombinant expression

The gene encoding mouse MFSD1 (Uniprot: Q9DC37) was cloned into a pXLG vector (Backliwal et al., 2008) containing an N-terminal Twin-Streptavidin-tag followed by a human rhinovirus 3C cleavage site, referred to as MFSD1-strep. The encoding sequence of mouse GLMP (Uniprot: Q9JHJ3) was cloned into the pXLG vector containing a C-terminal tobacco etch virus cleavage site, green fluorescent protein (GFP)-tag followed by an 8×Histidine (8×His)-tag, termed GLMP-Ct-His-GFP. A fusion construct of mouse GLMP and mouse MFSD1 connected by a linker region (GSAGSAAGSGEF) termed GLMP-MFSD1-strep was inserted into a pXLG vector with a C-terminal 3C-protease cleavage site followed by a Twin-Streptavidin-tag. Expi293F cells were transiently transfected as described elsewhere (Pieprzyk et al., 2018), and cells were harvested 48 h post-transfection. MFSD1-strep, coexpressed MFSD1-strep, and GLMP-Ct-His-GFP, referred to as GLMP+MFSD1, and GLMP-MFSD1-strep, referred to as GLMP-MFSD1, proteins were directly purified from the cell pellet by standard affinity purification. Briefly, the cell pellets were solubilized for 1 h at 4 °C in buffer containing 1x PBS pH 7.4, 150 mM NaCl, 5 mM MgCl_2_, 5% glycerol, 1% (w/V) n-dodecyl-β-D-maltopyranoside (DDM) detergent, 0.1 % (w/V) cholesterol hemisuccinate (CHS), 20 U/ml DNase I and EDTA-free protease inhibitors (Roche). The sample was centrifuged for 30 min at 35,000 ×g, and the supernatant was directly applied to Strep-TactinXT beads (IBA), incubated for 1 h at 4 °C, and loaded onto a gravity column. The beads were washed with 20 column volumes (CV) of washing buffer (1x PBS pH 7.4, 150 mM NaCl, 0.03% DDM, 0.003% CHS) prior to elution with 3 CV of size exclusion (SEC) buffer (20 mM HEPES pH 7.5, 150 mM NaCl, 0.03% DDM, 0.003% CHS) containing 10 mM desthiobiotin.

For GLMP+MFSD1, the elution fraction from the strep-tactin purification was incubated with Ni-NTA beads for 1 h at 4 °C and loaded onto a gravity column. The beads were washed with 10 CV of SEC buffer before elution with 3 CV of SEC buffer containing 250 mM Imidazole. TEV was added to the elution fraction, and the mixture was dialyzed against SEC buffer. The dialyzed sample was again incubated with Ni-NTA beads for 30 min at 4 °C, loaded onto a gravity column, and the flow-through collected and combined with that of one washing step of 2 CV of SEC buffer. The sample was then concentrated, as were the elution fractions of strep-tactin affinity purification of MFSD1-strep and GLMP-MFSD1-strep. Concentrated samples were applied onto either a Superose 6 increase 3.2/300 (Cytiva), in the case of GLMP-MFSD1 and GLMP+MFSD1, or a Superdex200 5/150(Cytiva) column for MFSD1 sample. For all samples, the columns were equilibrated in SEC buffer (20 mM HEPES pH 7.5, 150 mM NaCl, 0.03% DDM, and 0.003 % CHS). For Cryo-EM sample preparation, the size exclusion buffer contained 20 mM HEPES pH 7.5, 150 mM NaCl, 0.015% DDM, and 0.0015% CHS.

### Cloning and characterization of MFSD1 mutants for recombinant expression

Binding site mutations within the MFSD1 gene were generated via amplification of the mMFSD1 gene in combination with primers carrying the respective mutations, followed by SLiCE cloning (Zhang et al., 2014) of the amplified gene into a pXLG vector. For initial expression tests, the mutants and wildtype MFSD1 were cloned with an additional N-terminal 8×His and GFP-tag. Expression levels of each mutant were assessed by fluorescent size exclusion chromatography in comparison to the expression level of wildtype MFSD1. For this, the cell pellet of a 10 ml Expi293F culture overexpressing MFSD1 wildtype or mutant was solubilized in 1x PBS pH 7.4, 150 mM NaCl, 5 mM MgCl_2_, 5% glycerol, 1% (w/V) DDM detergent, 0.1% (w/V) CHS, 20 U/ml Dnase I and EDTA-free protease inhibitors (Roche) for 1 h at 4 °C. This was followed by ultra-centrifugation at 100,000 ×g for 1 h at 4 °C using a MLA130 rotor. The supernatant was then loaded onto a Superose 6 5/150 home-packed column, equilibrated in size exclusion buffer, monitoring the GFP-fluorescence at λ_ex_=488 nm/λ_em_=510 nm. Based on the expression and solubilization screening results, selected mutants were cloned into the pXLG vector carrying only an N-terminal Twin-Streptavidin- tag. The mutants were expressed and purified as wildtype MFSD1.

### LC-MS/MS-based analysis of dipeptides from tissues

#### Sample preparation

Aliquot (30 μL) of mouse embryonic fibroblasts (MEF) suspensions were mixed with 90 μL of ethanol. Aliquot (50 μL) of liver lysosome suspension was also mixed with 150 μL of ethanol. Aliquot of the liver (approximately 150 mg) was homogenized with PBS (150 μL) in a Biomasher II (Nippi, Tokyo, Japan). The homogenate was mixed with 900 μL of ethanol. The ethanol (75%) suspension was centrifuged at 10,000 g for 5 min after strong agitation. The supernatants were used for further analysis.

#### Derivatization with AccQ

Aliquots (100 μL) of 75% ethanol soluble fractions and peptide standards (1 mM, 20 μL) were dried under vacuum and dissolved into 80 μL of 50 mM sodium borate buffer, pH 8.8. Then, 20 μL of AccQ acetonitrile solution (0.3%) was added and kept at 50°C for 10 min. The reaction mixture was mixed with 100 μL of 5 mM sodium phosphate buffer, pH 7.5, and used as a sample for LC-MS/MS. For standard, the reaction mixture was further diluted to 1/10.

#### LC-MS/MS analyses

Aliquots (10 μL) of AccQ derivatives of standard peptide were injected into an electron spray ionization tandem mass spectrometer (LCMS-8040, Shimadzu, Kyoto, Japan) without using a column. Multiple-reaction monitoring (MRM) conditions for each AccQ-peptide were optimized using LaboSolution LCMS Ver 5.5 (Shimadzu) after the detection of monovalent and divalent ions.

Each peptide was determined by reversed-phase high-performance liquid chromatography-electron spray ionization tandem mass spectrometer (LC-MS/MS) equipped with an Inertsil ODS 3 column (2.1 mm ✕250 mm, GL Science, Tokyo, Japan). A binary gradient was carried out at a flow rate of 0.2 mL/min. The gradient program was as follows: 0-15 min, 0- 50% B; 15-20 min, 50-100% B; 20-25 min, 100% B; 25.01-35, 0% B. Detection was carried out in MRM mode. For sample and standard, 20 and 1 μL were injected, respectively.

### Thermal stability measurements

The unfolding of individual target proteins was followed by the nanoDSF method (Alexander et al., 2014). Purified wildtype and mutant MFSD1, or GLMP-MFSD1 and GLMP+MFSD1, was diluted to 0.2 mg/ml into nanoDSF buffer containing 100 mM HEPES pH 7.5, 150 mM

NaCl, 0.03 % DDM, 0.003 % CHS. 50 mM ligand stock solutions were prepared in 100 mM HEPES pH 7.5 buffer. The transporter was incubated at a ligand concentration of 5 mM at room temperature for 30 min before starting the nanoDSF measurement using a Prometheus NT.48 device. Measurements were performed in a temperature range from 20 °C to 95 °C in 1 °C/min increments. Melting temperatures were determined by the Nanotemper software and plotted usingGraphPad Prism. Estimation of K_D_ was performed as described in (Kotov et al., 2023).

### Reconstitution of MFSD1 into liposomes

For the liposome-based uptake assays, GLMP-MFSD1, GLMP+MFSD1 wildtype MFSD1, and MFSD1 mutants were reconstituted into liposomes containing 1-palmitoyl-2-oleoyl-sn-glycero-3-phosphoethanolamine (POPE), P1-palmitoyl-2-oleoyl-sn-glycero-3-phospho-1’-rac-glycerol (POPG) and CHS in a 3:1:1 (w/w) ratio. Lipids were mixed in chloroform, and the solvent was removed using a rotary evaporator. Dried lipids were washed twice with pentane, followed by solvent removal. The lipid film was resuspended in reconstitution buffer (50 mM potassium phosphate, pH 7.0) to a final lipid concentration of 20 mg/ml. On the day of the reconstitution, lipids were diluted to 5 mg/ml in reconstitution buffer and extruded through a 400 nm filter unit (Avanti). Preformed liposomes were disrupted with a final concentration of 0.075 % (w/w) Triton X-100 and incubated for 10 min at room temperature. Protein at a concentration of 0.5 mg/ml, or similar amounts of size exclusion buffer (empty control), was added to the lipids to reach a protein: lipid ratio of 1:60 (w/w), and the mixture was incubated at 4 °C for 1h. The detergent was removed by sequentially adding Bio-Beads SM-2 (Bio-Rad) and incubating overnight at 4 °C. The mixture was harvested, and the liposomes resuspended in reconstitution buffer, flash-frozen three times in liquid nitrogen, and then stored at -80°C until further use.

### Liposome-based pyranine assays

For liposome-based uptake assays (Parker et al., 2014), liposomes were thawed and harvested using a total amount of 5 µg of protein per experiment. The pelleted liposomes were resuspended in uptake buffer 1 (5 mM HEPES pH 6.8, 150 mM KCl, 2 mM MgSO_4_) containing 1 mM pyranine. The resuspended liposomes were subjected to seven freeze-thaw cycles in liquid nitrogen before being extruded through a 400 nm filter unit and then harvested. Excess pyranine was removed using a G-25 spin column (Cytiva) equilibrated in uptake buffer 1. Liposomes were again harvested and resuspended in uptake buffer 1 to a final volume of 4 µl per experiment.

Pyranine-loaded liposomes were diluted 1:50 into uptake buffer 2 (5 mM HEPES pH 6.8, 120 mM NaCl, 2 mM MgSO_4_) in a 96-black chimney deep well plate. The fluorescence of pyranine was measured at excitation wavelengths of 415 nm and 460 nm, with an emission wavelength of 510 nm for both excitations using a TECAN Spark2000 operating at 22 °C. Peptide or buffer was added after a short equilibration period to a final concentration of 2.5 mM. The uptake reaction was initiated after the addition of valinomycin at a final concentration of 1 µM. For analysis, the fluorescent counts at λ_ex_=415 nm/λ_em_=510 nm were divided by the fluorescent counts at λ_ex_=460 nm/ λ_em_=510 nm. The average value of the first 25 s after the addition of peptide was used for normalization, and the normalized counts were plotted against the assay time using Prism GraphPad. For bar graphs and K_M_ measurements, the initial uptake velocity in the linear range of the uptake curve after the addition of valinomycin was determined by linear regression using Prism GraphPad.

### Expression of MFSD1 and GLMP in Xenopus Oocytes

Xenopus oocytes were either purchased from Ecocyte Bioscience (Dortmund, Germany) or prepared from frogs housed in the local animal facility in compliance with the European Animal Welfare regulations (ethical agreement APAFiS #14316-2017112311304463 v4). Ovarian lobes were extracted from Xenopus laevis females under anesthesia, and oocyte clusters were incubated on a shaker in OR2 medium (85 mM NaCl, 1 mM MgCl_2_, 5 mM Hepes-K^+^ pH 7.6) containing 2 mg/mL of collagenase type II (GIBCO) for 1 h at room temperature. Defolliculated oocytes were sorted and kept at 19 °C in Barth’s solution (88 mM NaCl; 1 mM KCl; 2.4 mM NaHCO_3_; 0.82 mM MgSO_4_; 0.33 mM Ca(NO_3_)_2_; 0.41 mM CaCl_2_; 10 mM Hepes-Na^+^ pH 7.4), supplemented with 50 μg/mL of gentamycin.

Capped mRNAs were synthesized *in vitro* from the linearized pOX(+) plasmids using mMessage-mMachine SP6 kit (Invitrogen). Unless stated otherwise, defolliculated oocytes were injected with both mGLMP_Y400A_-mKate2 mRNA and mMFSD1_L11A/L12A_-EmGFP mRNA (25 ng each at 1 μg/μL). For co-expression with PQLC2, oocytes were injected with these two mRNAs and an mRNA-encoding rat PQLC2^L11A/L12A^-EGFP (Leray et al., 2021) at 16 ng each. Non-injected oocytes were used as negative controls.

### Cell surface biotinylation

Two days after injection, 5 oocytes were washed twice with ice-cold phosphate buffer saline (PBS) and biotinylated for 20 min at 4 °C using 2.5 mg/mL of the membrane-impermeable, cleavable reagent sulfo-NHS-SS-biotin EZ-LinkTM (Thermo Scientific). After four washes, oocytes were lysed for 30 min in 500 μl lysis buffer (150 mM NaCl, 5 mM EDTA, 50 mM Tris-HCl pH 7.5, 0.1% SDS, 1% Triton X-100, Halt Protease Inhibitor Cocktail). Cell lysates were clarified by sedimentation at 14.000 × g for 10 min, and the supernatant was incubated for 2 h at 4 °C with 150 μL streptavidin-agarose beads (Sigma-Aldrich) under gentle agitation. Beads were sedimented at 100 ×g for 30 s. Supernatants (unbound material) were recovered, and beads were washed three times with 1 ml lysis buffer. Streptavidin-bound material was then eluted in 100 μL Laemmli’s sample buffer. Half of the bound proteins were resolved by 10% SDS/PAGE. Following electrophoresis, transfer, and blocking, the nitrocellulose membrane was incubated with mouse anti-GFP antibodies (1∶1000, Roche Molecular Biochemicals) and rabbit anti-mKate2 antibodies (1:2000, Origene). The protein bands were obtained using horseradish peroxidase-conjugated antibodies against mouse whole immunoglobulins and horseradish peroxidase-conjugated antibodies against rabbit whole immunoglobulins (1∶10,000, Sigma-Aldrich) as secondary antibodies and detection with the Lumi-Light Plus Western Blotting Substrate (Roche). Images were acquired and quantitated with an ImageQuant LAS 4000 chemiluminescence imager (GE Healthcare).

### Two-electrode voltage clamp (TEVC) recording in Xenopus oocytes

Electrophysiological recordings were done at room temperature (20 °C), usually two days after cRNA injection. For each experiment, mMFSD1^L11A/L12A^-EmGFP expression at the plasma membrane was verified under an Eclipse TE-2000 epifluorescence microscope (Nikon) with a 4× objective focused at the equatorial plane. Voltage clamp was applied with two borosilicate-glass Ag/AgCl microelectrodes filled with 3 M KCl (0.5 to 3 MΩ tip resistance) connected to an O725C amplifier (Warner Instrument) and a Digidata 1440A interface controlled via pCLAMP 10.7 software (Molecular Devices). Currents were filtered using a 10-Hz low-pass filter and sampled at 1 kHz. Solutions were applied with a gravity-fed perfusion system in a Xenoplace™ recording chamber (ALA Scientific Instruments) with built-in Ag/AgCl reference electrodes. Oocytes were perfused in ND100 medium (100 mM NaCl, 2 mM KCl, 1 mM MgCl_2_, and 1.8 mM CaCl_2_) buffered with 10 mM 2-(N-morpholino)ethanesulfonic acid (MES)-NaOH to pH 5.0 unless stated otherwise. After recording a stable baseline current, peptides (10 mM unless stated otherwise) were applied in this medium and eventually washed to measure the evoked current. For peptides purchased as hydrochloride salts, the substrate-free solution was supplemented with *N*- methyl-*D*-glucamine (NMDG) hydrochloride (Sigma-Aldrich) at the same concentration to avoid interference with the Ag/AgCl reference electrode. For Lys-Ala application, the substrate-free solution was supplemented with the same concentration of sodium bromide (Merck) to avoid an endogenous current artifact induced upon bromide washing.

### Combined TEVC and pH_in_ recording in Xenopus oocytes

In these experiments, a third ion-selective electrode connected to an FD223a dual channel differential electrometer (World Precision Instruments) was impaled into the oocyte. To prepare this intracellular pH electrode, a silanized micropipette with dichlorodimethylsilane (Sigma) was tip-filled with a proton ionophore (Hydrogen ionophore I, cocktail B, Sigma-Aldrich) and backfilled with 150 mM NaCl, 40 mM KH_2_PO_4_, 23 mM NaOH pH 6.8. The two channels of the FD223a electrometer were connected to the pH electrode and the voltage ground electrode of the TEVC setup, respectively. The potential difference between the two inputs tested in diverse buffers (pH range: 5.0-7.5) was proportional to pH with a mean slope of -82 mV (n = 3). The relative level of substrate-evoked intracellular acidification was quantified by the slope, in mV/s, of the ion-selective electrode voltage trace.

### Leu(d_3_)-Ala uptake into Xenopus oocytes

Two days after cRNA injection, oocytes were washed and individually incubated in 200 µL ND100 pH 5.0 medium supplemented, or not, with 10 mM Leu(*d_3_*)-Ala for 20 min at room temperature. After 3 washes in 0.55 mL ice-cold ND100 pH 5.0 medium, oocytes were transferred into 100 µL ice-cold methanol/water (50:50) and homogenized by pipetting up and down with a P1000 tip. After centrifugation for 5 min at 4°C and 16,000×*g*, supernatants were collected and stored at -20°C before analysis. In experiments for absolute quantification of the Leu(*d_3_*)-Ala flux, a subset of MFSD1/GLMP oocytes was treated before (5 min) and during the transport reaction with 10 µM of the branched-chained amino acid transaminase inhibitor BAY-069 (Gunther et al., 2022) to prevent metabolization of the accumulated leucine. Quantification of dipeptides and amino acids in oocyte extracts was done by LC-MS/MS. Lysis supernatants were diluted 20-fold in water, and 20 microliters of the dilution were injected into a reverse phase column (Phenomenex-C18, 2.1 × 150 mm; 3 µm). The mobile phases were water with 0.1% formic acid for phase A and acetonitrile with 0.1% formic acid for phase B. Elution was programmed to start at 100% phase A for 3 min, then fall to 20% phase A at 10 min, return to 100% phase A at 11 min and equilibrate for 6 min prior to the next sample injection. The flow rate was 0.3 mL/min, and the detection was done using an 8060NX triple quadrupole mass spectrometer (Shimadzu) with an electrospray ion probe (ESI) operated at 250° C. The selected ions monitored (SIM) and multiple reaction monitoring (MRM) are listed with the retention times in **Suppl. Table 2**. Quantification was done by integrating the chromatographic peak area using Labsolution software (Shimadzu, France). For absolute quantification, a calibration curve was established with various known concentrations (0.2 to 100 µM) of Leu(*d_3_*), Ala and Leu(*d_3_*)-Ala standards.

### CryoEM sample preparation and data collection

Gel filtration peak fractions containing GLMP-MFSD1 were used for CryoEM sample preparation. For the apo state structure, grids at a concentration of 3.33 mg/ml purified GLMP-MFSD1 were prepared. For GLMP-MFSD1 in the presence of the dipeptide His-Ala, termed GLMP-MFSFD1_His-Ala_, purified GLMP-MFSD1 at 3 mg/ml was dialyzed over night against buffer containing 150 mM NaCl, 0.015 % (w/V) DDM, 0.0015% (w/V) CHS supplemented with 20 mM His-Ala. 3.6 µl of purified protein were applied onto glow-discharged holey carbon-coated grids (Quantifoil R1.2/1.3 Au 300 mesh) and blotted for 3.5 s with a blot force of 0 at 100% humidity and 4 °C before being frozen in liquid ethane using a Vitrobot Mark IV (Thermo Fisher Scientific). Data were collected in counted super-resolution mode, with a binning of 2, on a Titan Krios (Thermo Fisher Scientific) equipped with a K3 camera and a BioQuantum energy filter (Gatan) set to 15 eV. For the two data sets, movies were collected at a nominal magnification of × 81,000 with a pixel size of 1.1 Å using EPU (Thermo Fisher Scientific). For the GLMP-MFSD1_apo_ structure, two separate data sets were collected, consisting of 3179 and 2551 movies. For the GLMP-MFSD1_His-Ala_ structure, one dataset consisting of 3193 movies was collected. For both GLMP-MFSD1_apo_ and GLMP- MFSD1_His-Ala_, data were collected at a dose rate of 15 e^-^/pixel/second, with an exposure time between 4 and 4.2 seconds to reach a total dose of 55 e^-^/Å^2^.

### CryoEM data processing and modeling

Data processing was performed in cryoSPARC (Punjani et al., 2017). Collected movies were subjected to patch motion correction with a maximum alignment resolution of 4 Å. After CTF estimation using CTFFIND4 (Rohou and Grigorieff, 2015), micrographs were curated based on CTF Fit resolution and total full-frame motion. Particles were selected using Blob picker with a minimum particle diameter of 100 Å and a maximum particle diameter of 200 Å, followed by manual inspection and adjustment of the NCC score (>0.49) and local power to reduce duplicate particle picks and picking of ethane contaminations on the sample. Particles were extracted with a 256-pixel (GLMP-MFSD1_apo_) and 300-pixel (GLMP-MFSD1_His-Ala_) box size, followed by several rounds of 2D classification. Particles of the final 2D classification were subjected to *ab initio* reconstruction of 4 classes. Upon visual inspection, two reconstructions, one representing the ’model class’ and the other one a ’decoy class’, depicting a corrupted model, were selected for heterogeneous refinement of the whole particle stack used in the previous *ab initio* reconstruction. The resulting ’model class’ after heterogeneous refinement was then subjected to non-uniform refinement (Punjani et al., 2020), and the resulting reconstruction was used for another round of heterogeneous refinement while the ’decoy class’ stayed the same. After several rounds of these two steps, per particle local motion correction (Rubinstein and Brubaker, 2015) was performed, followed by one more non-uniform refinement step that resulted in maps of a final global resolution of 4.2 Å and 4.1 Å for GLMP-MFSD1_apo_ and GLMP-MFSD1_His-Ala_, respectively.

Initial model fitting was performed in UCSF ChimeraX (Goddard et al., 2018). A first model of MFSD1, representing an inward open conformation in complex with GLMP, was obtained by AlphaFold2 Multimer (Jumper et al., 2021) using both protein sequences as input. First, the model of MFSD1 was manually placed into the experimental density, and the fit was refined in UCSF ChimeraX (Goddard et al., 2018). Then, the model was refined in Cartesian space using the Rosetta/StarMap workflow (Lugmayr et al., 2023) with the map resolution set to 7.5 Å. Next, the GLMP model was manually placed into the density, followed by fit refinement in UCSF ChimeraX, and the complex model was refined again with the same settings in Rosetta/StarMap. The model with the highest iFSC metric of 0.64 (Wang et al., 2016) was selected for downstream analyses. The model was further fit into the map with ISOLDE (Croll, 2018). Subsequent model building and refinement were iteratively performed in Coot (Emsley et al., 2010) and PHENIX (Liebschner et al., 2019). Figures were generated using PyMOL and UCSF ChimeraX. Electrostatic surfaces were generated using the APBS plugin provided in PyMOL (Jurrus et al., 2018).

### MD simulation of ligand-bound MFSD1

The MFSD1 structures were placed in a heterogenous bilayer composed of POPE (20%), 1- palmitoyl-2-oleoyl-glycero-3-phosphocholine (POPC, 30%), Cholesterol (30%), and N-Palmitoyl-sphingomyelin (PSM, 20%) using CHARMM-GUI scripts (Jo et al., 2008). The protonation states of titratable residues were determined using the MCCE program (Song et al., 2009). For the substrates, both termini are assigned charged. In the case of the dipeptide His-Ala, both neutral and charged side chains were simulated. All systems were hydrated with 150 mM NaCl electrolyte. The all-atom CHARMM36m force field was used for lipids, ions, cofactors, and protein with TIP3P water. Molecular dynamics (MD) trajectories were analyzed using Visual Molecular Dynamics (VMD) (Humphrey et al., 1996) and MDAnalysis (Michaud-Agrawal et al., 2011).

All simulations were performed using GROMACS 2021.3. A description of the dipeptide simulations performed for this study is provided in **Supplemental Figure 10**. The conditions

and substrates for MD analyses are summarized in **Supplemental Table 3**. Starting systems were energy-minimized for 5,000 steepest descent steps and equilibrated initially for 500 ps of MD in a canonical (NVT) ensemble and later for 7.5 ns in an isothermal-isobaric (NPT) ensemble under periodic boundary conditions. During equilibration, the restraints on the positions of non-hydrogen protein atoms of initially 4,000 kJꞏmol^-1^ꞏnm^2^ were gradually released. Particle-mesh Ewald summation with cubic interpolation and a 0.12-nm grid spacing was used to treat long-range electrostatic interactions. The time step was initially 1 fs and was increased to 2 fs during the NPT equilibration. The LINCS algorithm was used to fix all bond lengths. The constant temperature was established with a Berendsen thermostat, combined with a coupling constant of 1.0 ps. A semi-isotropic Berendsen barostat was used to maintain a pressure of 1 bar. During production runs, a Nosé-Hoover thermostat and a Parrinello–Rahman barostat replaced the Berendsen thermostat and barostat. Analysis was carried out on unconstrained simulations.

### Indirect immunofluorescence and microscopy

Semi-confluent cells were grown on glass coverslips and fixed 48 hours after transfection for 20 min with 4% (w/v) paraformaldehyde at room temperature. Cells were permeabilized, quenched, and blocked with normal goat serum before incubation with primary antibodies overnight at 4°C. The coverslips were washed and incubated for 90 min with AlexaFluor dye-conjugated secondary antibodies (Thermo Fisher Scientific). Afterward, the coverslips were washed four times and mounted on microscope slides with mounting medium including DAPI (4-,6-diamidino-2-phenylindole). An Airyscan2 980 laser scanning microscope (Zeiss) equipped with a 63x oil immersion objective (NA = 1.40) was used for microscopy. The images were acquired and processed with the Zen 3.2 (Blue edition) software. The Pearson correlation coefficient was calculated with the Zen 3.2 (Blue edition) software.

### SDS-PAGE and immunoblotting

SDS-PAGE and immunoblotting were performed according to standard protocols. Protein lysates were transferred to the nitrocellulose membrane by semi-dry blotting. For MFSD1- immunoblotting, lysates were denatured for 10 min at 55 °C prior to SDS-PAGE. The protein bands were detected using horseradish peroxidase-conjugated and detection with the Lumi-Light Plus Western Blotting Substrate (Roche). Luminescence was detected with an ImageQuant LAS 4000 chemiluminescence imager (GE Healthcare).

### Enrichment of lysosomal fractions from the mouse liver

Liver lysosome enrichment was performed according to a previously published method (Markmann et al., 2017; Massa Lopez et al., 2019). Four days prior to the experiment, mice were injected intraperitoneally with 4 µl/g body weight with 17% (v/v) Tyloxapol diluted in 0.9% NaCl. The mice were sacrificed in a CO_2_-flooded chamber. The liver was removed immediately and homogenized in three volumes of isotonic 250 mM sucrose solution in a Potter-Elvejhem and a glass homogenizer (B. Braun type 853202) with five strokes. The homogenate was centrifuged for 10 min at 1,000 ×g to remove unbroken cells and nuclei. The pellet was re-extracted in the same volume of 250 mM sucrose solution in the Potter- Elvejhem and centrifuged again. The supernatants were pooled (postnuclear supernatant, PNS) and transferred to ultracentrifugation tubes. In the first differential centrifugation step, lysosomes and mitochondria were enriched by centrifugation of the pooled PNS at 56,500 ×g for 7 min at 4 °C (Beckman-Coulter, 70 Ti fixed-angle rotor). The supernatant was removed, and the pellet was resuspended in 250 mM sucrose solution. The resuspended solution was centrifuged again for 7 min at 56,500 ×g, and the supernatant was carefully discarded. The differential centrifugation was followed by a discontinuous sucrose gradient: The final pellet was resuspended in a volume of 3.5 ml sucrose solution with a density of ρ 1.21 and transferred into a new ultracentrifugation tube. This fraction was carefully layered with a sucrose solution of a density of ρ 1.15 (3 ml), ρ 1.14 (3 ml), and ρ 1.06 (0.5 ml). The gradient was centrifuged for 2.30 hours at 4 °C and 111,000 ×g in a swing-out rotor (Beckman-Coulter, SW41). The brownish lysosome-enriched fraction (∼ 1ml) was collected from the interface between the ρ 1.14 and ρ 1.06 sucrose layers. All animal experiments were approved by the local authorities.

### Untargeted metabolomics and targeted metabolite quantitation

The polar metabolites were profiled using a Thermo Fisher Scientific ID-X Tribrid mass spectrometer with an ESI probe. Metabolite separation before mass spectrometry was achieved through HILIC, conducted using a MilliporeSigma SeQuant ZIC-pHILIC 150-mm by 2.1-mm column (cat# 1504600001) along with a 20-mm by 2.1-mm guard (cat# 1504380001). Mobile phases consisted of 20 mM ammonium carbonate and 0.1% ammonium hydroxide dissolved in 100% LC-MS grade water (Phase A) and 100% LC-MS grade acetonitrile (Phase B). The chromatographic gradient involved a linear decrease from 80 to 20% of Phase B from 0 to 20 minutes, followed by a linear increase from 20 to 80% from 20 to 20.5 minutes, and maintaining at 80% from 20.5 to 29.5 minutes. The LC flow rate and injection volume were set to 0.15 ml/min and 1.5-3 μL, respectively. The mass spectrometer settings included Orbitrap resolution of 120,000, positive and negative ion voltages of 3,000 V and 2,500 V, respectively, an ion transfer tube temperature of 275 °C, a vaporizer temperature of 350 °C, an RF lens at 40%, an AGC target of 1 × 10^6^, and a maximum injection time of 80 ms. A full scan mode with polarity switching at m/z 70 to 1,000 was executed. Gas flowrates: sheath, 40U; aux, 15U; sweep 1 U. Internal calibration was achieved by EasyIC.

Metabolite samples were pooled by combining replicates for data-dependent MS/MS collection. The orbitrap resolution was set at 240,000, HCD stepped energies at 15, 30, and 45%, an isolation window at 1 m/z, intensity threshold at 2 × 10^4^, and exclusion duration at 5 seconds, AGC target at 2 × 10^6^, maximum injection time at 100 ms. Both isotope and background exclusions were enabled, with background exclusion being performed via AcquireX (ThermoFisher Scientific). TraceFinder (Thermo Fisher Scientific) was used in combination with a library of known metabolite standards (MSMLS, Sigma-Aldrich) for targeted metabolite quantification. The mass tolerance for extracting ion chromatograms was set at 5 ppm.

### Statistics

GraphPad Prism 9.3.1 (GraphPad Software) was used for data representation and calculation of statistic testing. The statistic test applied for each graph is indicated in the figure legends. For most panels, a two-tailed paired t-test, a two-tailed unpaired t-test, or a nonparametric Mann-Whitney U test was used. Statistical differences in the graphs were generally depicted as ns = not significant; * p ≤ 0.05; ** ≤ 0.01; *** p ≤ 0.001. Error bars in the graphs represent the standard error of the mean (SEM) or standard deviation (SD), as indicated in the figure legends.

## RESULTS

### Dipeptides accumulate in the lysosomes of MFSD1-deficient mice

To identify the substrate(s) potentially exported by MFSD1 from lysosomes to the cytoplasm, we enriched lysosomes from wildtype and *Mfsd1* knockout mice (*Mfsd1^tm1d/tm1d^)* (Massa Lopez et al., 2019) by differential centrifugation followed by a sucrose density gradient (**Figure 1A**). This procedure yields fractions highly enriched for the lysosomal markers LAMP1 and cathepsin D (CTSD), containing little contamination from other organelles, including the Golgi apparatus, the endoplasmic reticulum, or mitochondria (**Figure 1B**) (Markmann et al., 2017). These lysosome-enriched fractions were subsequently used for a differential untargeted mass spectrometry-based metabolomics analysis. Only two metabolites showed significant differences above the defined thresholds (p ≤ 0.05, fold change ≥2) and were tentatively identified as dipeptides containing one uncharged and one cationic amino acid: Arg-Pro (or Pro-Arg) and Pro-Lys (**Figure 1C**).). In fact, The extracted ion chromatograms (EICs) of the metabolite with m/z [M+H] 272.1717, which is putatively identified as Arg-Pro or Pro-Arg, show that the retention time of this dipeptide matches perfectly with that of the authentic chemical standard, and its level is significantly higher in *Mfsd1 ^tm1d/tm1d^* lysosomes (**Figure 1D**). Tandem mass spectrometry (MS/MS) analysis against spectral libraries additionally confirmed the identity of both dipeptides (**Suppl. Figure 1A**). Quantification of Pro-Lys and Arg-Pro and targeted analysis of additional dipeptides (Arg- HydroxyPro, anserine) revealed a pronounced increase of these dipeptides in liver lysosomes from *Mfsd1* knockout mice compared to the wildtype controls (**Figure 1E**). Analysis and quantification of different dipeptides in different organs (liver, spleen, lung) (**Suppl. Figure 1B**) demonstrate an increase of anserine, Arg-Pro, Pro-Arg, and Arg- HydroxyPro in spleen lysates but not in the other analyzed organs of *Mfsd1* knockout mice.

**Figure 1.**
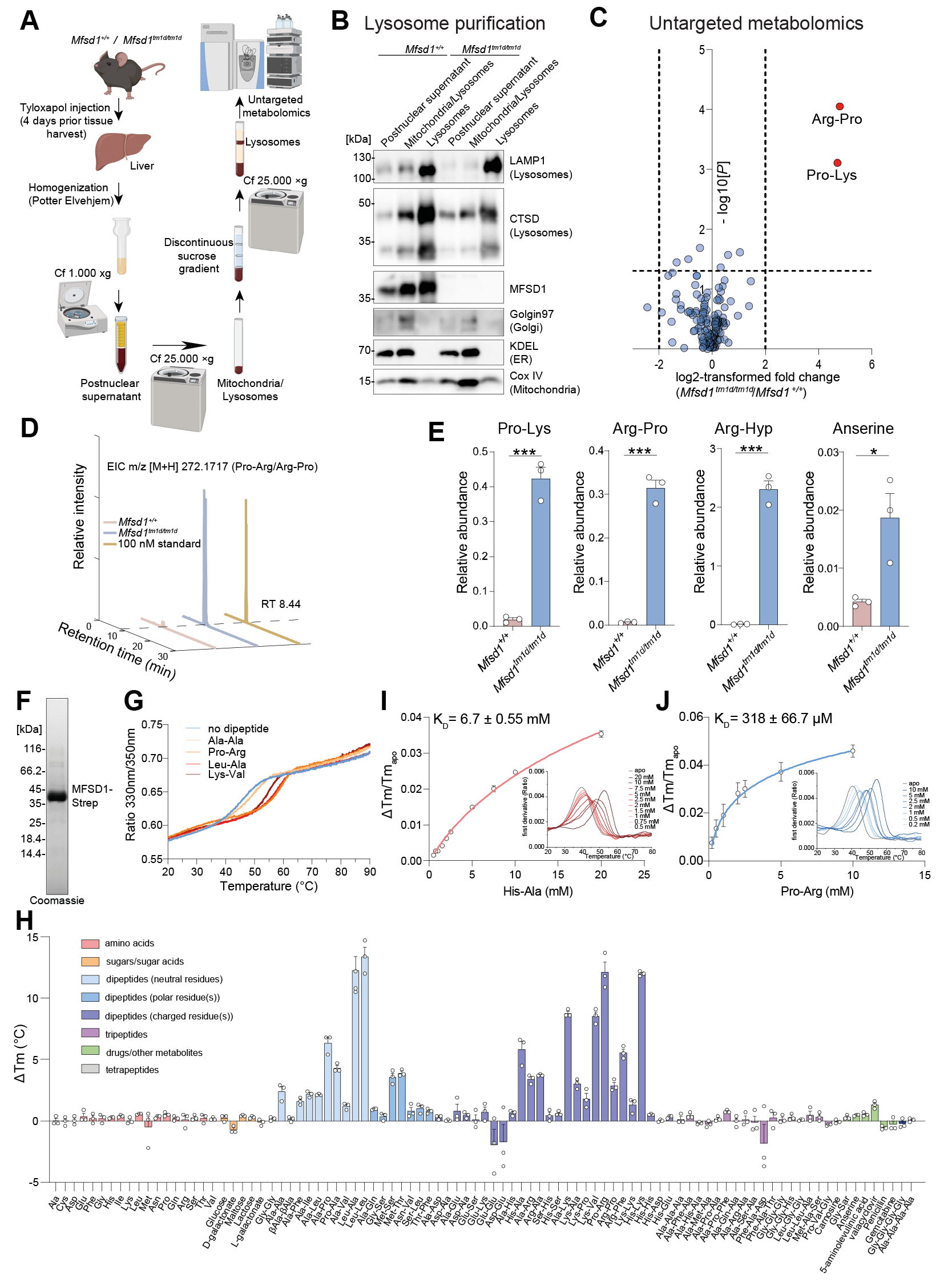
*Mfsd1* knockout mice accumulate cationic dipeptides in liver lysosomes, and recombinant MFSD1 binds various dipeptides. **(A)** Schematic representation of the lysosome enrichment by ultracentrifugation and untargeted metabolomics. **(B)** Immunoblot analysis of postnuclear supernatant (PNS), mitochondria, and lysosome-enriched fractions and the final lysosome-enriched fraction from wildtype and *Mfsd1* knockout mice for markers of various cellular compartments. **(C)** Volcano plot of differential metabolites between liver lysosomes of wildtype and *Mfsd1* knockout mice. **(D)** Extracted ion chromatogram (EIC) for the chemical standard Pro-Arg (yellow, 100 nM) and representative samples from wildtype (red) and *Mfsd1* knockout mice (blue). Pro- Arg is detected as a peak eluting at a retention time of 8.44 min. **(E)** Relative abundance of Pro-Lys, Arg-Pro, and anserine between wildtype and *Mfsd1* knockout mice. The abundance was normalized to the isotopically-labeled arginine levels, which did not show any differences between the two genotypes in the untargeted metabolomic analysis. Two-tailed unpaired t-tests. * p ≤ 0.05; *** p ≤ 0.001 **(F)** Coomassie-stained SDS-PAGE gel of purified MFSD1 with a Twin-Streptavidin-tag. **(G)** Unfolding traces of MFSD1 in the absence and presence of Ala-Ala, Pro-Arg, Leu-Ala, and Lys-Val at a concentration of 5 mM. **(H)** Thermal stability of MFSD1 in the presence of a compound library at 5 mM final ligand concentration. Changes in the melting temperature of MFSD1 (ΔTm) are given as a difference to the melting temperature of apo MFSD1 (Tm_apo_). Experiments were done in triplicates. **(I, J)** Examples of K_D_ measurements are based on changes in the thermal stability of MFSD1 in the presence of varying concentrations of the dipeptides His-Ala (red) **(I)** or Pro-Arg (blue) **(J)**. K_D_ values were determined using Moltenprot (Kotov et al., 2021).

### Recombinant MFSD1 binds dipeptides

The metabolomics data prompted us to test whether MFSD1 is involved in lysosomal peptide transport. MFSD1 was transiently expressed in Expi293F cells and purified to homogeneity in detergent solution (**Figure 1F, Suppl. Figure 1C**). To screen for potential interactions with peptides, MFSD1 was subjected to thermal shift experiments using differential scanning fluorimetry (nanoDSF) (**Figure 1G-J**). Upon interaction of MFSD1 with a substrate, the protein is stabilized, resulting in an increased melting temperature (T_m_) compared to its apo form. Initial nanoDSF experiments at 5 mM ligand concentration showed a stabilization effect on MFSD1 by Leu-Ala, Lys-Val, and Pro-Arg but not with Ala-Ala (**Figure 1G**). To further investigate the variety of molecules MFSD1 might bind, we performed a larger screen using nanoDSF covering 18 amino acids, 68 di- and tripeptides, two tetrapeptides, five sugars, and seven drug molecules (**Figure 1H**). The strongest stabilization effects were observed for neutral dipeptides (e.g., Leu-Leu, ΔT_m_= 14 ⁰C) and dipeptides that possess at least one positively-charged residue (e.g., Pro-Arg, ΔT_m_= 12.1 °C and His-Lys, ΔT_m_= 12 °C) (**Figure 1H**). No, or only small, thermal shift changes of MFSD1 were detected for any of the other compound classes screened (**Figure 1H**), indicating that MFSD1 primarily binds dipeptides. Titration experiments of varying concentrations of His-Ala, His-Lys, Leu-Ala, Lys-Val or Pro-Arg in nanoDSF experiments yielded dissociation constants (K_D_) of 6.7 ± 0.55 mM, 765 ± 136 µM, 2.2 ± 0.42 mM, 4.3 ± 0.6 mM and 318 ± 66.7 µM (**Figure 1I, J; Suppl. Figure 1D**), respectively (Kotov et al., 2023). The determined K_D_ values are within the range of reported binding affinities of other peptide transporters of the MFS family (Flayhan et al., 2018; Kotov et al., 2023; Martinez Molledo et al., 2018b; Scow et al., 2011; Ural-Blimke et al., 2019).

### Uptake of dipeptides by MFSD1 and GLMP

We next tested whether MFSD1 not only binds but also transports dipeptides using a whole-cell transport assay in *Xenopus* oocytes. In this approach, the lysosomal transporter of interest is misrouted to the plasma membrane by mutagenesis of its lysosomal sorting motif(s), replacing the poorly tractable lysosomal export with whole-cell import. The transport reaction is initiated by adding the substrate in an acidic extracellular medium to mimic the pH of the lysosomal lumen (Kalatzis et al., 2001; Ruivo et al., 2012). Expression of an MFSD1 sorting mutant (Massa Lopez et al., 2019) fused to Emerald Green Fluorescent Protein (EmGFP), MFSD1_L11A/L12A_-EmGFP, in *Xenopus* oocytes resulted in limited localization to the plasma membrane, as determined by cell surface biotinylation. However, co-expression of a sorting mutant of GLMP (the accessory subunit of MFSD1, (Massa Lopez et al., 2019)), GLMP_Y400A_-mKate2, increased the surface level of MFSD1 by ∼10-fold (**Figure 2A**). Dual-color fluorescence microscopy confirmed this effect and showed colocalization of the EmGFP and Kate2 signals in co-injected oocytes (**Figure 2B**), indicating localization of the MFSD1/GLMP complex at the plasma membrane. We thus used oocytes co-expressing MFSD1_L11A/L12A_-EmGFP and GLMP_Y400A_-mKate2 (‘MFSD1/GLMP oocytes’) for the transport assays.

**Figure 2.**
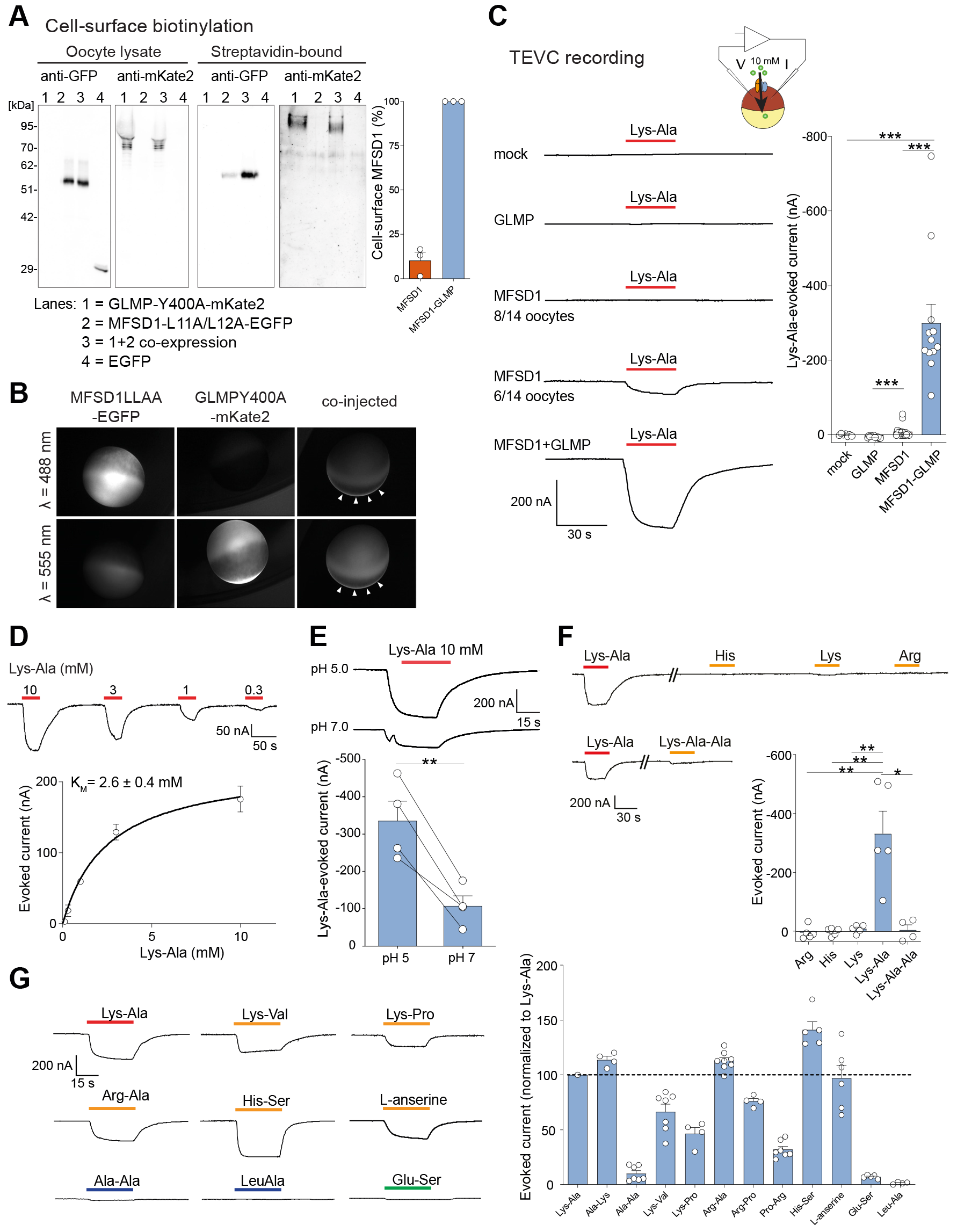
Cationic dipeptides evoke an inward current in MFSD1/GLMP-expressing oocytes. **(A)** Surface biotinylation analysis of Xenopus oocytes expressing MFSD1L11A/L12A-EmGFP and/or GLMPY400A-mKate2. Oocytes expressing EGFP in the cytosol validated the selectivity of surface labeling in streptavidin-bound fractions. Western blots are representative of three independent experiments. **(B)** Fluorescence micrographs of representative oocytes. Arrowheads show MFSD1/GLMP colocalization at the plasma membrane. **(C)** TEVC recording of oocytes clamped at -40 mV and perfused with 10 mM Lys-Ala at pH 5.0. Traces show representative Lys-Ala-evoked currents of 7 to 14 oocytes per expression condition. About 60% of the oocytes expressing only MFSD1L11A/L12A-EGFP did not respond to Lys-Ala. P values were calculated using Mann-Whitney U tests (*** p ≤ 0.001). **(D)** Dose-response relationship of the Lys-Ala current in MFSD1/GLMP oocytes. The current follows Michaelis–Menten kinetics with a KM of 2.6 ± 0.4 mM (mean ± SEM of n = 3 oocytes). **(E)** Lys-Ala was applied to each MFSD1/GLMP oocyte at pH 5.0 and pH 7.0. Two-tailed paired t-test: ** p ≤ 0.01. **(F)** Response of MFSD1/GLMP oocytes to cationic amino acids and to the tripeptide Lys-Ala-Ala (10 mM each) at pH 5.0. P values were calculated using Mann-Whitney U tests:* p ≤ 0.05, ** p ≤ 0.01. **(G)** Response of MFSD1/GLMP oocytes to diverse dipeptides compared to Lys- Ala (4 to 11 oocytes per substrate).

Oocytes were recorded under a two-electrode voltage clamp (TEVC) at -40 mV, and dipeptides (10 mM) were applied at extracellular pH (pH_out_) 5.0 to test them for electrogenic transport (**Figure 2C**). Interestingly, Lys-Ala evoked a robust inward current (-300 ± 50 nA, mean ± SEM, n=12 oocytes) in MFSD1/GLMP oocytes but not in mock (non-injected) oocytes nor oocytes expressing only GLMP_Y400A_-mKate2, while it evoked at best, a very low current (-9.2 ± 4.8 nA, n=14) in oocytes expressing only MFSD1_L11A/L12A_-EmGFP (**Figure 2C**).

Varying the Lys-Ala concentration showed that the MFSD1/GLMP current follows Michaelis-Menten kinetics, with a K_M_ of 2.6 ± 0.4 mM (n = 3) (**Figure 2D**). This current was ∼3-fold stronger at pH_out_ 5.0 than pH_out_ 7.0 (**Figure 2E**), in agreement with the role of MFSD1/GLMP at the lysosomal membrane. In contrast, the MFSD1/GLMP current did not depend on the Na^+^ ion (**Suppl. Figure 2A**).

We then characterized the substrate specificity of the MFSD1/GLMP current. Single cationic amino acids (His, Lys, or Arg) and the tripeptide Lys-Ala-Ala (10 mM) did not evoke any current in MFSD1/GLMP oocytes (**Figure 2F**), in agreement with the dipeptide selectivity of the nanoDSF data (**Figure 1H**). In contrast, several cationic dipeptides such as Ala-Lys, Arg-Ala, His-Ser, Arg-Pro and, to a lesser extent, Lys-Pro and Pro-Arg, evoked a robust current, whereas neutral dipeptides (Leu-Ala; Ala-Ala) and an anionic dipeptide (Glu-Ser) had no effect (**Figure 2G, Suppl. Figure 2B**). We performed competition experiments to test whether neutral or anionic dipeptides interact with MFSD1/GLMP in oocytes. Leu-Ala (20 mM) applied simultaneously with Lys-Ala (3 mM) abolished the Lys-Ala current (**Suppl. Figure 2C**), in agreement with its strong interaction with purified MFSD1 (**Figure 1H**). Ala-Ala (20 mM) and Glu-Ser (10 mM) also inhibited the Lys-Ala current by 66 ± 3% (n=6) and 26 ± 3% (n=3), respectively (**Suppl. Figure 2D, E**). We concluded that MFSD1/GLMP interacts with diverse dipeptides in the oocyte membrane and that it transports cationic dipeptides in an electrogenic manner.

As an alternative *in vitro* approach, the transport activity was characterized using purified MFSD1_WT_ reconstituted into POPE:POPG:CHS (3:1:1) lipid vesicles (**Figure 3A**). To monitor possible proton-coupling by MFSD1_WT_ as observed for other lysosomal transporters (Kalatzis et al., 2001; Morin et al., 2004; Ruivo et al., 2012), the liposomes were loaded with the pH-sensitive dye pyranine (Kano and Fendler, 1978). A membrane potential of around -100 mV was applied using valinomycin to drive the uptake of dipeptides (**Figure 3B**). As a control experiment, liposomes devoid of MFSD1, termed empty liposomes, were analyzed under identical conditions. Time-dependent uptake assays of empty and MFSD1-containing liposomes highlight that only proteoliposomes in the presence of the dipeptide, His-Ser, exhibit a decrease in fluorescence (F_norm_.) after the addition of valinomycin (**Figure 3C**). All other traces did not show a signal change over a time period of 10 min (**Figure 3C**). Since this method allowed us to follow the uptake of protons, we screened a similar set of dipeptides used in the oocyte-based assays (**Figure 3D**) and determined the Michaelis-

**Figure 3.**
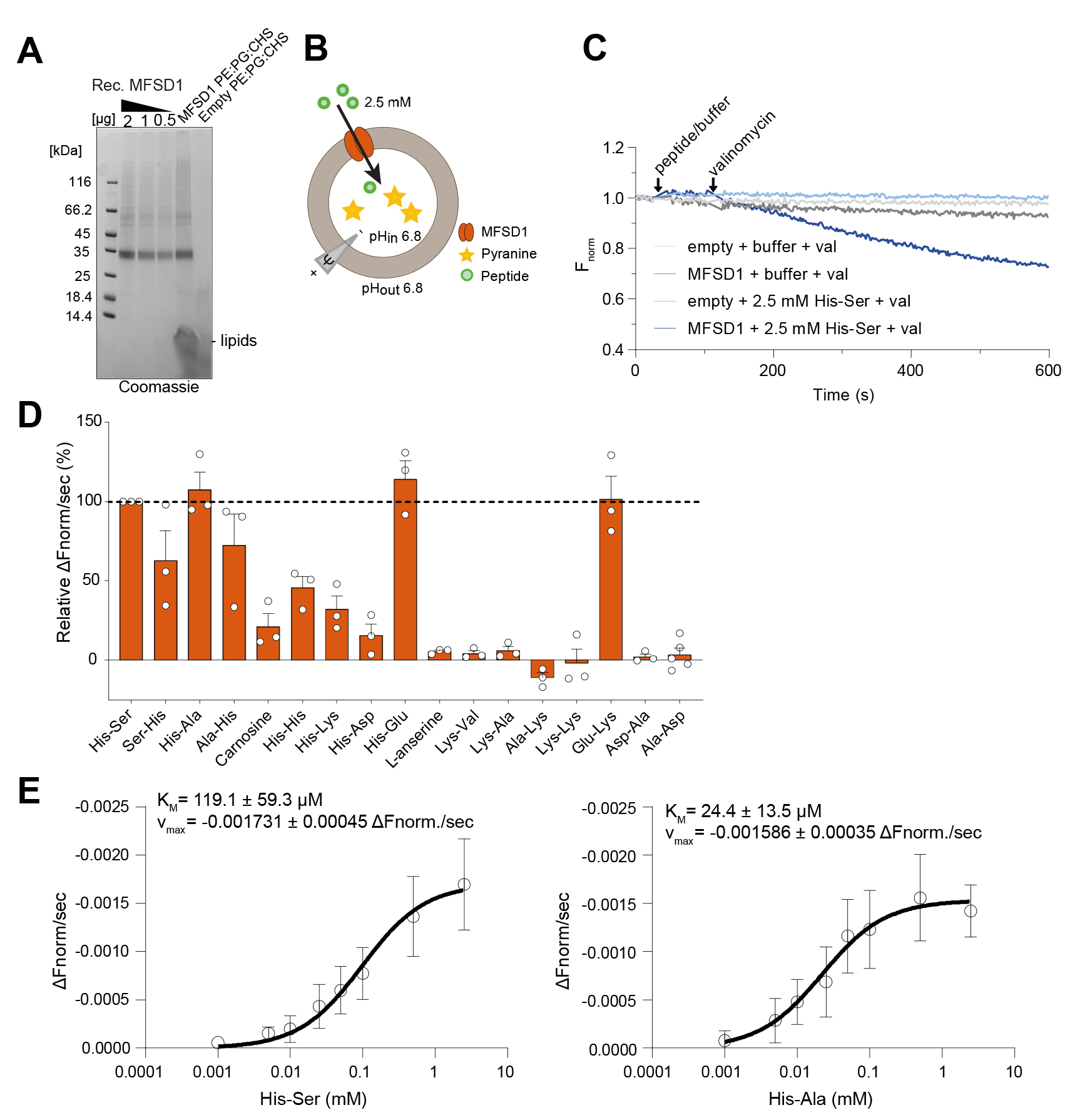
MFSD1 is active as a dipeptide transporter in a liposome-based assay. **(A)** Coomassie-stained SDS-PAGE gel of MFSD1 after reconstitution into POPE:POPG:CHS liposomes (PE:PG:CHS). **(B)** Schematic of the experimental setup of liposome-based transporter assay. **(C)** Representative traces of time-course measurements of uptake in the presence of 2.5 mM His-Ser (HS) and 1 µM valinomycin (val) using MFSD1-containing liposomes (mmMFSD1) and those devoid of protein (empty). The addition of peptide or buffer and valinomycin during the measurements is indicated by arrows. **(D)** Substrate specificity of MFSD1 measured for liposome-based uptake assays. Initial uptake rates for each peptide are given as a percentage of the determined initial uptake rate of His-Ser. Data are shown as mean ± SD of three independent experiments. **(E)** Michaelis-Menten kinetics of uptake of His-Ser and His-Ala by MFSD1. K_M_ and v_max_ values were calculated from three independent experiments using Prism GraphPad. Individual data points are plotted as mean ± SD.

Menten kinetics for the dipeptides His-Ala and His-Ser (**Figure 3E**). The resulting K_M_ values were 119.1 ± 59.3 µM for His-Ser and 24.4 ±13.5 µM for His-Ala with v_max_ of -0.001731 ± 0.00045 ΔF_norm_/sec and -0.001586 ± 0.00035 ΔF_norm_/sec, respectively (**Figure 3E**).

Intriguingly, uptake was exclusively observed for peptides containing at least one Histidine residue, with Glu-Lys being the only exception (**Figure 3D**). Therefore, although the liposome

and oocyte activities shared common features (strong His-Ser signal; lack of response to neutral and anionic dipeptides), they diverged for a subset of cationic dipeptides, such as Lys-Ala, Ala-Lys, Lys-Val, and L-anserine, which evoked a robust inward current in the oocyte assay, yet had no effect in the liposome assay.

### MFSD1 operates as a dipeptide uniporter

To investigate the origin of this apparent discrepancy, we examined whether MFSD1 co- transports protons, as initially postulated, using combined TEVC and intracellular pH (pH_in_) recording of MFSD1/GLMP oocytes (**Figure 4A**). We used two approaches to analyze the sensitivity of the pH_in_ microelectrode impaled in the oocyte. First, we co-expressed MFSD1/GLMP with the lysosomal uniporter for cationic amino acids PQLC2 (sorting mutant PQLC2_L290A/L291A_-EGFP) to serve as a positive control (Jezegou et al., 2012; Leray et al., 2021). The uptake of cationic histidine by PQLC2 induces an intracellular acidification, reflecting the release of its sidechain proton (pKa = 6.0) when the substrate faces the cytosol (pH = 7.2). In contrast, the uptake of lysine and arginine induces an inward PQLC2 current but no intracellular acidification, in agreement with their higher side chain pKa (10.5 and 12.5, respectively) (Leray et al., 2021). We verified that PQLC2 does not respond to Lys-Ala (**Suppl. Figure 3A**), allowing us to monitor the MFSD1/GLMP and PQLC2 activities independently. Sequential application of Lys-Ala and His to MFSD1/GLMP + PQLC2 oocytes showed that the uptake of Lys-Ala by MFSD1/GLMP does not evoke any intracellular acidification under conditions where the pH_in_ microelectrode detects a slower flux of cationic histidine through PQLC2 (**Figure 4A**, **Suppl. Figure 3B**), ruling out an H^+^ symport mechanism for MFSD1/GLMP (**Figure 4B**). Second, we compared the responses evoked by Lys-Ala and His-containing dipeptides in MFSD1/GLMP oocytes. Similar to the uptake of His by PQLC2, His-containing dipeptides should release their sidechain proton within the oocyte if MFSD1/GLMP transports them in cationic form. Indeed, His-Ala and His-Ser, but not Lys- Ala, evoked intracellular acidification in MFSD1/GLMP oocytes (**Figure 4C, D**). To quantify the rate of acidification, we normalized the current and pH_in_ signal (initial slope) evoked by each substrate to those evoked by His-Ala in the same oocyte and used the ratio between the normalized acidification signal and the normalized current as a proxy for the number of protons released per elementary charge during substrate translocation (**Figure 4E, F**). This analysis yielded ratios of 1.2 ± 0.1 (n=4) and -0.08 ± 0.03 (n=4) proton per elementary charge for His-Ser and Lys-Ala, respectively, excluding further the H^+^ symport model and corroborating the concept of cytosolic acidification caused by the release of proton(s) bound to the translocated substrate (**Figure 4E, F**).

**Figure 4.**
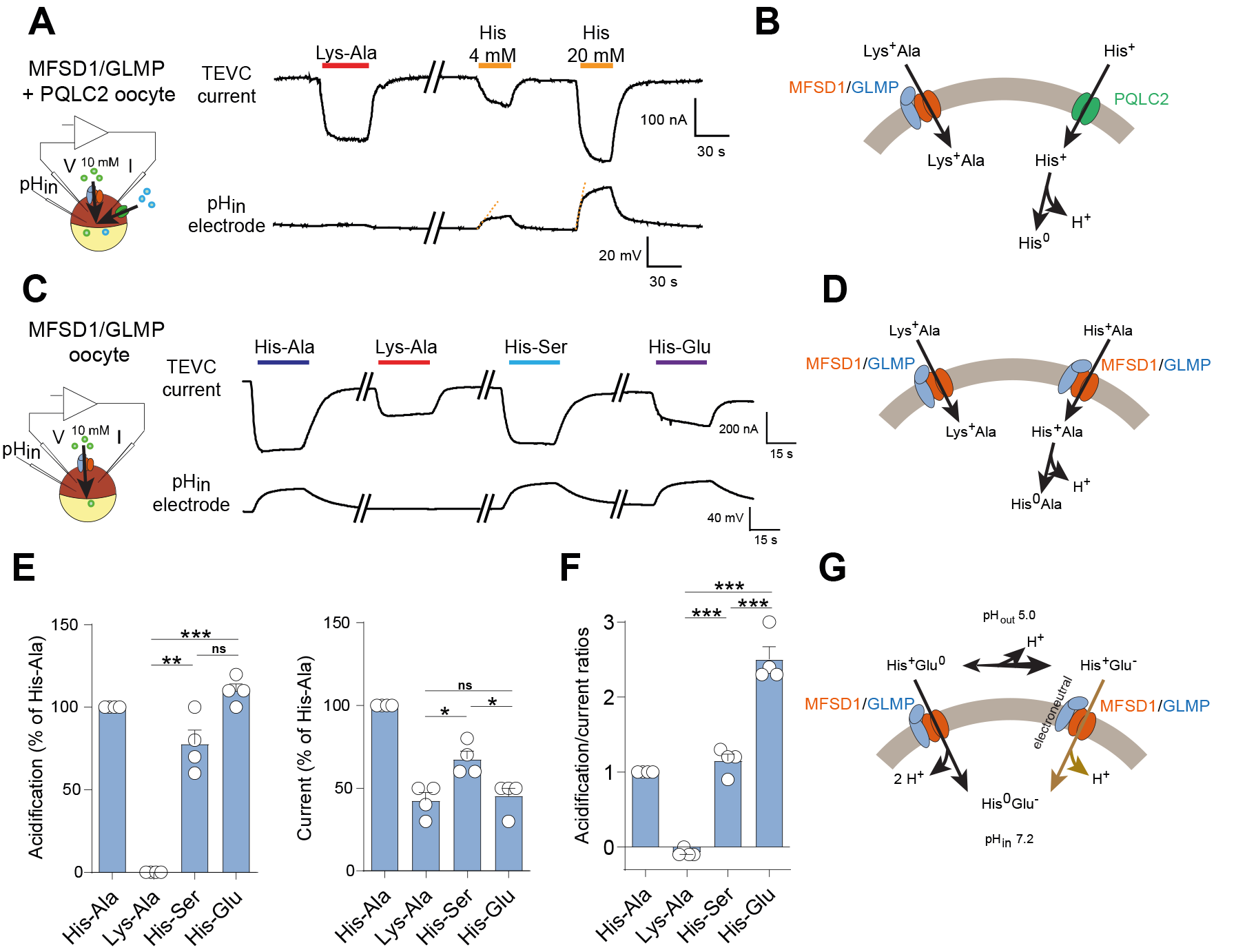
MFSD1 is a dipeptide uniporter. **(A)** Combined TEVC and intracellular pH (pH_in_) recording of oocytes expressing both MFSD1/GLMP and PQLC2 (sorting mutant L290A/L291A) at their surface. His, but not Lys-Ala, applied at pH 5.0, induces intracellular acidification (orange dotted lines). The traces are representative of five oocytes shown in **Suppl. Figure 4B**. **(B)** Model for the acidification induced by His following its release from PQLC2. **(C)** Combined TEVC and pH_in_ recording of an MFSD1/GLMP oocyte perfused with the indicated dipeptides (10 mM) at pH 5.0. **(D)** A model accounting for the selective acidification by His-containing dipeptides. **(E)** The experiment in (C) was repeated on four MFSD1/GLMP oocytes. Data are means ± SEM of the acidification and current responses normalized to His-Ala. ns = not significant; * p ≤ 0.05; ** ≤ 0.01; *** p ≤ 0.001. **(F)** Normalized acidification/current ratios provide the number of protons released per translocated elementary charge for each substrate. **(G)** A model accounting for the high number of protons released by His-Glu. At the tested potential (-40 mV), ∼67% of His-Glu molecules would be taken up by MFSD1/GLMP in the minor cationic form, His^+^-Glu^0^, releasing 2 protons per elementary charge, while ∼33% would be taken up in the predominant zwitterionic form, His^+^-Glu^-^, releasing another proton in an electroneutral manner.

To test this model further, we measured the responses induced by the application of His-Glu. This dipeptide exists in four protonation states: a zwitterionic form (His^+^-Glu^-^), which predominates in the perfusion medium (pH_out_ = 5.0, one unit above the sidechain pKa of Glu: 4.1); a cationic form, His^+^-Glu^0^, with a protonated Glu residue; an anionic form, His^0^-Glu^-^, with a deprotonated His residue; and low amounts of the neutral form, His^0^-Glu^0^. The addition of His-Glu evoked both an inward current and intracellular acidification in MFSD1/GLMP oocytes with, remarkably, an acidification/current ratio of 2.5 ± 0.2 (n=4) (**Figure 4C, E, F**). MFSD1/GLMP thus substantially transports His-Glu in cationic form, despite its lower abundance at pH_out_ 5.0, since this cationic form must release two protons per elementary charge (**Figure 4G**). The deviation above this theoretical ratio of 2.0 implies that His-Glu is also taken up in other protonation states. Partial uptake (1/3 of the total His-Glu flux) of the predominant zwitterionic form, which carries additional protons in an electroneutral manner, would account for the observed ratio of 2.5 (**Figure 4G**). Finally, we tested the dipeptide Glu-Lys, which stood out as an atypical substrate in the proteoliposome assay. Glu-Lys also evoked both an inward current and intracellular acidification in MFSD1/GLMP oocytes, with an acidification/current ratio of 1.2 ± 0.2 protons per elementary charge, in agreement with an entry in protonated, cationic state Glu^0^-Lys^+^ (**Suppl. Figure 4A, B, C**). Additional uptake of Glu-Lys in its predominant zwitterionic form, Glu^-^-Lys^+^, may also occur as it cannot be detected by the dual TEVC/pH_in_ recording technique.

Taken together, these data highlight that MFSD1 is not coupled to protons. It operates instead as a dipeptide uniporter. The apparent discrepancy between the proteoliposome assay and the oocyte assay thus only reflects the inability of the former to detect the transport of substrates that do not carry, and subsequently release, a proton bound to their side chain(s).

### MFSD1 efficiently transports Leu-Ala

The observation that MFSD1 transports ∼30% of His-Glu molecules in zwitterionic form (**Figure 4F, G**) prompted us to test whether it can transport *bona fide* neutral dipeptides into oocytes using a biochemical assay. As preliminary targeted LC-MS/MS analysis of MFSD1/GLMP oocyte extracts did not show accumulation of a robust substrate like Lys-Ala, we used a stable isotope tracer approach to monitor the transport activity. Leu-Ala, a good binder of MFSD1 both in vitro (**Figure 1H**) and *in cellula* (**Suppl. Figure 2C**), was synthesized in deuterated form (Leu(*d_3_*)-Ala) and applied at 10 mM to MFSD1/GLMP or mock oocytes for 20 min at pH 5.0. Oocyte extracts were then analyzed by targeted LC-MS/MS analysis (**Figure 5A**). Leu(*d_3_*)-Ala showed a very poor, yet significant, accumulation in MFSD1/GLMP oocytes incubated with Leu(*d_3_*)-Ala. In contrast, these oocytes, but not mock oocytes, dramatically accumulated deuterated leucine (Leu(*d_3_*)) (**Figure 5B, C**), showing that Leu(*d_3_*)-Ala is actually transported by MFSD1, yet quickly cleaved within the oocyte by intracellular peptidases. In agreement with this interpretation, MFSD1/GLMP oocytes incubated with Leu(*d3*)-Ala also accumulated ‘light’ alanine over its endogenous

**Figure 5.**
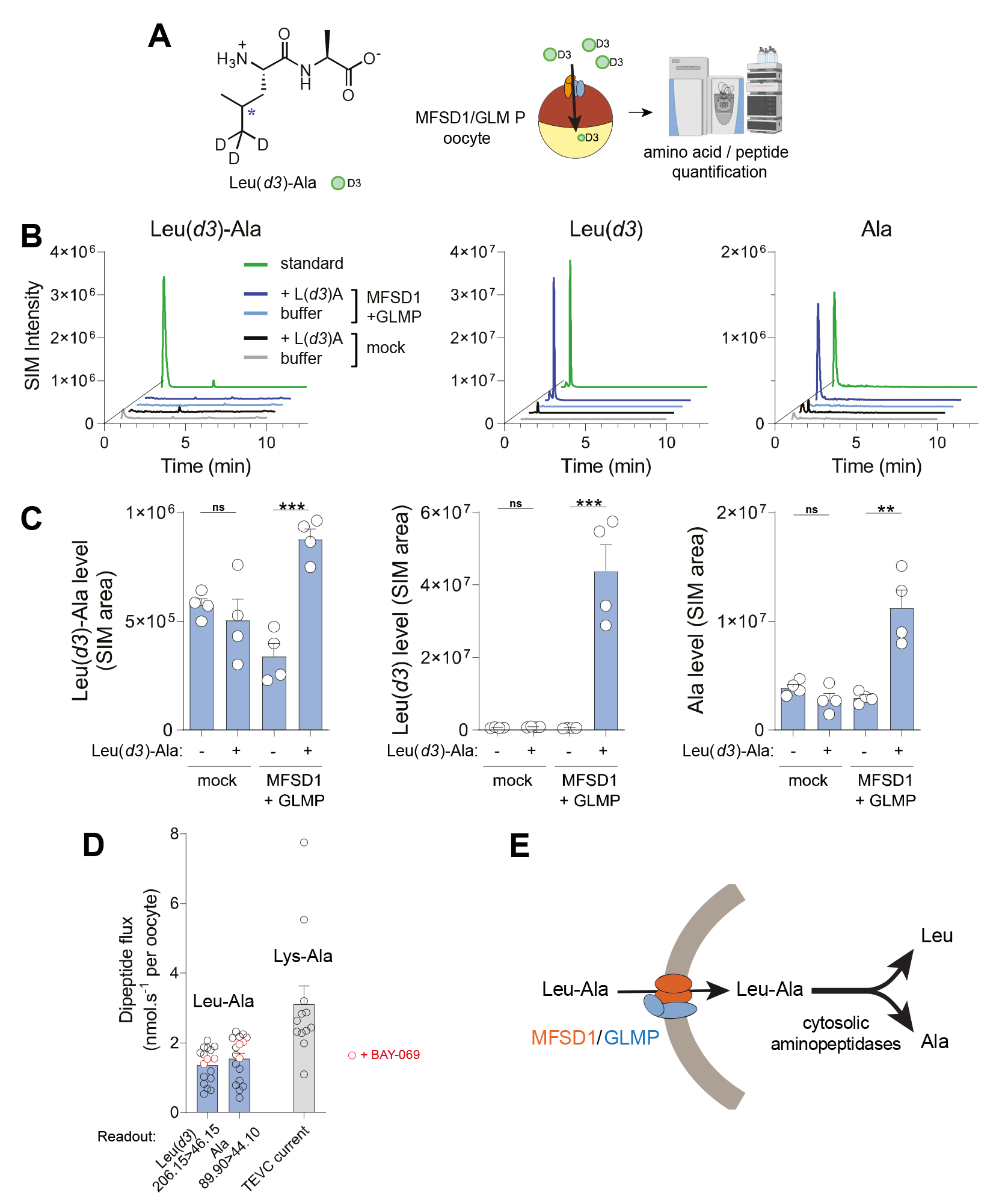
MFSD1 efficiently transports Leu-Ala. **(A)** Heavy isotope tracer approach used to monitor Leu-Ala transport. **(B)** Representative LC-MS chromatograms of ≥ 5 independent experiments. The amount of standard (green lines) was 3.9 pmol for Leu(*d3*)-Ala, and 15.6 pmol for Leu(*d3*) and Ala. **(C)** Relative quantification of the chromatographic peak area of Leu(*d3*)-Ala, Leu(*d3*) and Ala in extracts from mock and MFSD1/GLMP oocytes, in presence or absence of, 10 mM Leu(*d3*)- Ala for 20 min at pH 5.0. Data are means ± SEM of 4 oocytes from a representative example of 3 independent experiments. Two-tailed unpaired t-tests: ns = not significant; * p ≤ 0.05; ** ≤ 0.01; *** p ≤ 0.001. **(D)** Absolute quantification of Leu-Ala uptake. Oocyte extracts were analyzed by LC-MS/MS along with a range of standard concentrations. The 46.15 and 44.10 MS2 fragment ions of Leu(*d3*) and Ala, respectively, were used as quantifiers to determine the absolute amount Leu(*d3*) and Ala accumulated over time upon incubation of MFSD1/GLMP oocytes with Leu(*d3*)-Ala. Data are means ± SEM of 17 oocytes from 2 oocyte batches. In one experiment, some oocytes were treated with the branched-chained amino acid transaminase inhibitor BAY-069. Lys-Ala currents from **Figure 2C** were divided by the Faraday constant and plotted with the same scale (grey bar) to allow comparison between Leu-Ala and electrogenic substrates. **(E)** A model accounting for the LC-MS/MS data.

level (**Figure 5B, C**). The dependence of Leu(*d_3_*) accumulation on Leu(*d3*)-Ala concentration follows Michaelis-Menten kinetics with a K_M_ of 4.4 mM (**Suppl. Figure 5A**), a value similar to that of Lys-Ala (**Figure 2D**). To compare how the *rate* of Leu-Ala transport stands among MFSD1 substrates, we performed absolute quantification of the Leu(*d_3_*) and Ala signals during the linear phase of Leu(*d_3_*)-Ala uptake with time (**Suppl. Figure 5B**) and compared these measurements with the current evoked by electrogenic MFSD1 substrates. This quantification yielded a Leu-Ala transport rate of 1.32 ± 0.14 pmol.s^-1^ per MFSD1/GLMP oocyte (n = 10) for the Leu(*d_3_*) signal, and 1.52 ± 0.20 pmol.s^-1^ per oocyte (n = 10) for the Ala signal (**Figure 5D**). This rate is about half that of Lys-Ala (3.11 ± 0.52 pmol.s^-1^ per oocyte, n = 12; **Figure 2C, 5D**), which, in contrast with Leu-Ala, is actively driven by the negative potential of the oocyte membrane (-40 mV) in the TEVC assay. Of note, the absolute quantification also showed that Leu(*d_3_*) and Ala are released at equimolar levels (Ala/Leu(*d_3_*) ratio = 1.09 ± 0.09, n = 3) following Leu(*d_3_*)-Ala import (**Suppl. Figure 5C**).

### CryoEM structure determination of GLMP-MFSD1

In order to elucidate the molecular mechanism of substrate recognition, we determined the structure of MFSD1 in the apo and dipeptide-bound states. Since MFSD1 is a small membrane protein (∼51 kDa) and lacks any distinguishable structural features that would guide cryo-EM particle alignment, we made use of the known interaction of GLMP with MFSD1 (Massa Lopez et al., 2019).

First, we tested if the interaction of MFSD1 and GLMP is stable *in vitro*. For this, MFSD1 and GLMP were individually or co-expressed in Expi293F cells (**Suppl. Figure 6A**), and a subsequent pull-down assay confirmed that MFSD1 forms an intact complex with GLMP even after detergent-extraction from the cellular membrane (**Suppl. Figure 6A**). In addition, we designed a fusion construct connecting GLMP with MFSD1 via a Glycine/Serine linker (termed GLMP-MFSD1).

The purified GLMP+MFSD1 co-complex (**Suppl. Figure 6B**) and the GLMP-MFSD1 fusion construct (**Suppl. Figure 6E**) exhibited similar stabilization effects by dipeptides as MFSD1_WT_ (**Suppl. Figure 6C and 6F**). They were more thermostable than MFSD1_WT_ (T_m_=40 °C), though the transport activity in proteoliposomes was reduced for the fusion, whereas the purified complex was as active as MFSD1_WT_ (**Suppl. Figure 6D, 6G**). Since the GLMP- MFSD1 fusion protein could be purified at high yields and still exhibits peptide interaction despite reduced uptake activity, we used this construct for subsequent structure determination. For the apo structure of GLMP-MSFD1 (GLMP-MFSD1_apo_), we obtained a 3D reconstruction at a nominal resolution of 4.2 Å (**Suppl. Figure 6A-C**, **Suppl. Table 4**). A second sample was prepared to determine the structure in a substrate-bound state. For this, the confirmed substrate His-Ala (**Figure 3D**) was chosen. Due to the low binding affinity of His-Ala to MFSD1 (K_D_=6.7 ± 0.55 mM) (**Figure 3E**), we made use of the buffering abilities of histidine-containing peptides and dialyzed purified GLMP-MFSD1 against His-Ala-containing buffer, replacing the current sample buffer by 20 mM His-Ala. For the GLMP-MSD1_His-Ala_ sample, we obtained an overall reconstruction at a resolution of 4.1 Å (**Suppl. Figure 8A-C**, **Suppl. Table 4**), though the luminal domain of GLMP and core parts of MFSD1 reach a local resolution up to 3.5 Å (**Suppl. Figure 8C**). Given the slightly higher resolution of this data set, we used this reconstruction to build the model of GLMP-MFSD1, guided by an AlphaFold2 (Tunyasuvunakool et al., 2021) model of the complex. The EM map resolved most of both proteins, including N-glycans, on the luminal domain of GLMP (**Figure 6A, B, Suppl. Figure 9A-C**). Missing regions include residues 1-35 and 392-404 for GLMP, as well as residues 99-100, 135-141, and 178-181. For MFSD1, residues 1-35, 446-464, and the inter-domain loop region (residues 241-260) could not be modeled.

**Figure 6.**
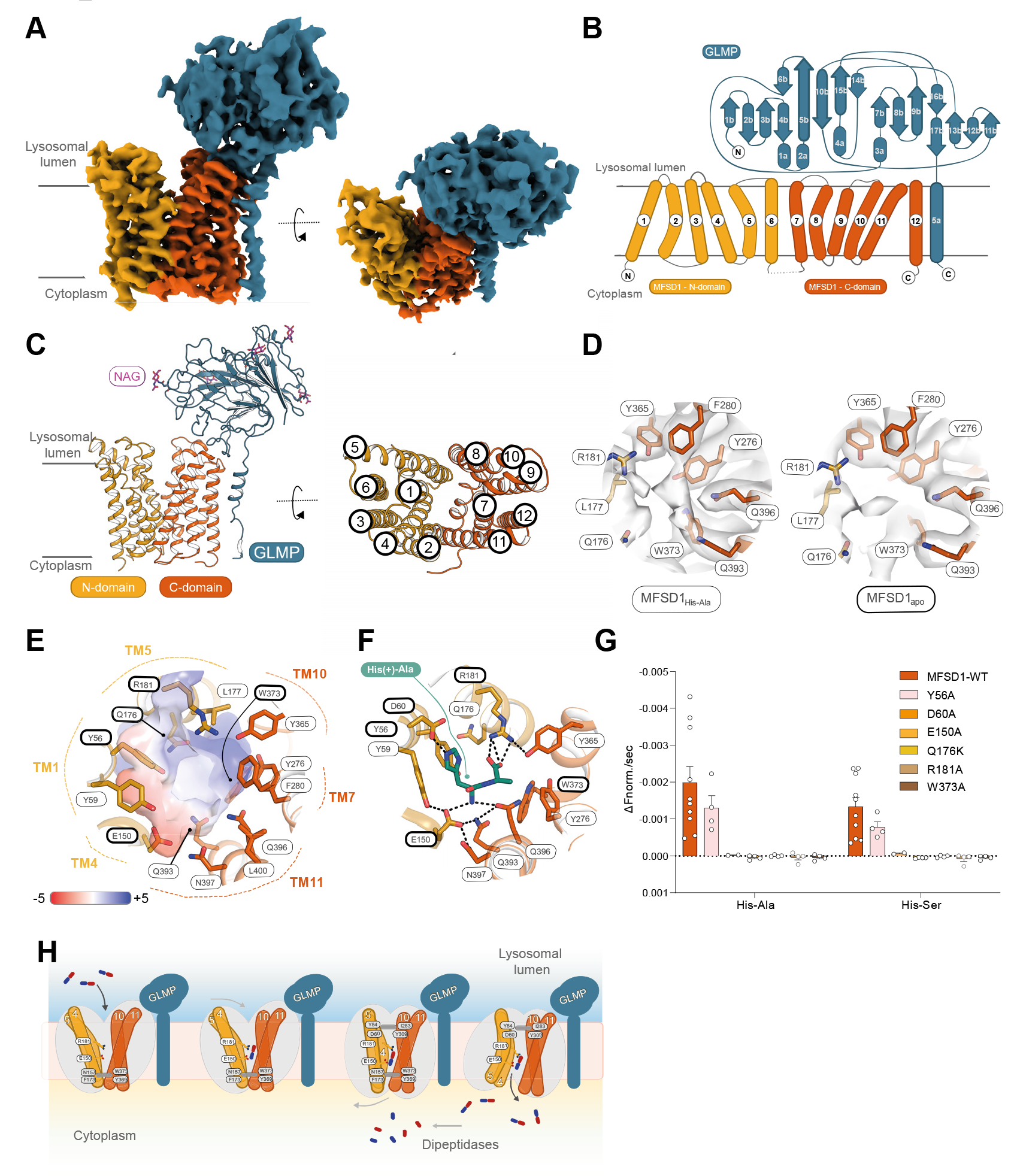
The outward open structure of GLMP-MFSD1. **(A)** Cryo-EM map of GLMP- MFSD1_His-Ala_. The N- and C-domain of MFSD1 are colored yellow and orange, respectively. GLMP is colored blue. **(B)** Topology diagram of MFSD1 and GLMP. N- and C-termini are labeled, and secondary structure elements are numbered. **(C)** Cartoon representation of GLMP-MFSD1 with top view of MFSD1. The numbering of TMs is indicated. Sugar modifications (Acetylglucosamine, NAG) identified on GLMP are colored pink. **(D)** Additional binding site density was found for the GLMP- MFSD1 data set in the presence of the dipeptide with His-Ala (MFSD1_His-Ala_) compared to the apo data set (MFSD1_apo_). The maps (light grey) are shown at σ=6. Surrounding residues are labeled. **(E)** The electrostatic surface, calculated with the APBS plug-in in PyMol, highlights the bipolar character of the binding site. Residues that were mutated in this study are framed in bold black. **(F)** Binding of the protonated dipeptide His(+)-Ala (green) as observed after 300 ns of MD simulations. Hydrogen bonds are indicated as dashed black lines, and residues used for mutational studies are framed in bold black. (G) Effect of mutations of binding site residues on uptake of His-Ala or His-Ser compared to MFSD1WT. Uptake rates are given as mean ± SD for n=8 (MFSD1WT) or n=4 (mutants) of independent experiments. (H) Schematic of transport of dipeptides by the GLMP/MFSD1 complex. GLMP in complex with MFSD1 resides in the outward open apo form (1st column) with the binding site open to the lysosomal lumen. From the unknown pool of proteolytic products, MFSD1 recognizes and transports only dipeptides, illustrated as blue (N-terminus) and red (C-terminus) sticks. The cytoplasmic side of MFSD1 is blocked by residues N157, F173, W373, and Y369, forming the cytoplasmic gate (shown as a grey bar). Peptide binding is coordinated by residues E150 and R118 highlighted in each model) leading to the outward-open substrate-bound form of MFSD1 (2nd column), upon which the N-terminal domain (yellow) with helices 4 and 5 moves closer to the C-terminal domain (orange). The lysosomal gate (shown as an additional grey bar) constituted by residues Y8, D60, I283, and Y309 closes after substrate binding, resulting in the occluded substrate-bound conformation of MFSD1 (3rd column). Subsequently, the cytoplasmic half of the N-terminal domain of MFSD1 moves away from the C-terminal domain and opens up the binding site to the cytosol (inward-open substrate- bound conformation; 4th column), followed by substrate release. Dipeptides are possibly then further processed by dipeptidases in the cytoplasm to individual amino acids.

MFSD1 is captured in an outward open conformation where the binding site is accessible from the lysosomal lumen (**Figure 6A, B, C**). The transmembrane domains of MFSD1 adopt the canonical MFS fold formed by twelve transmembrane helices (TM) organized in two six- helix bundles (N-domain by TM1-6 and C-domain by TM7-12) with both termini facing the cytoplasm (**Figure 6B, C**) (Drew et al., 2021; Pao et al., 1998; Reddy et al., 2012). For GLMP, the luminal domain and its single-span transmembrane helix could be resolved (**Figure 6A, B, C, Suppl. Figure 9B**). The transmembrane helix of GLMP is located directly adjacent to the C-domain of MFSD1. For the luminal domain, we could identify five of the six N-linked glycosylation sites present in a previously determined X-ray structure of the luminal domain (PDB: 6NYQ) and confirmed *in vivo* (Lopez et al., 2020) (**Suppl. Figure 9C**). N- Glycosylation sites are located at Asn85, Asn94, Asn157, Asn228 and Asn331 (**Figure 6C**). The luminal domain of GLMP adopts a β-sandwich fold (**Figure 6B**) that is structurally similar to a dimerization domain found in a cellodextrin phosphorylase from *Clostridium thermocellum* (PDB: 5NZ7) (O’Neill et al., 2017).

### The substrate binding site of MFSD1

A comparison of the 3D reconstructions of both data sets, apo vs. His-Ala bound, revealed an additional density for the latter (**Figure 6D**) in the cavity between the two helical bundles of MFSD1 (**Figure 6C, D**). This potential binding site is located approximately halfway into the membrane-spanning region and is formed by TM1 (Tyr56, Tyr59), TM4 (Glu150), and TM5 (Gln176, Arg181) of the N-domain and TM7 (Tyr276, Phe280), TM10 (Tyr365, Trp373), TM11 (Gln393, Gln396, Asn397, Leu400) of the C-domain (**Figure 6D**). The cavity exhibits a bi-polar surface character mainly caused by residues Glu150 and Arg181, which possibly help in orienting the dipeptide in the binding cavity (**Figure 6E**).

Owing to the insufficiently resolved density of the potential His-Ala peptide (**Figure 6D**), unambiguous placement of the peptide was not possible. To further investigate the peptide binding of MFSD1, we performed molecular dynamics (MD) simulations in a lipid bilayer in the presence of different dipeptides. The dipeptides Leu-Ala, Lys-Ala, and His-Ala (with either a neutral or positively charged histidine, His^0^-Ala or His^+^-Ala, respectively) were placed in two different orientations based on the peptide density observed in the Cryo-EM reconstruction of GLMP-MFSD1_His-Ala_ using only the model of MFSD1 for the simulations. Ligands placed in peptide orientation 1 (PO1) had their C-terminus positioned towards a patch of polar residues, namely Gln393, Gln396, and Asn397, and the side chain of the first dipeptide residue is pointing in the direction of Arg181 (**Suppl. Figure 10**). For the ligands placed in peptide orientation 2 (PO2), the dipeptide’s N- and C-termini are near residues Glu150 and Arg181, respectively (**Suppl. Figure 1**). Starting from peptide orientation 2 (PO2), the N- and C-termini of the dipeptide are in the vicinity of residues Glu150 and Arg181, respectively (**Suppl. Figure 11**). After 300 ns of simulation time, peptides starting from PO2 deviate less from their starting pose while peptides in PO1 flipped (**Suppl. Figure 11**) so that their N- and C-termini interact with Glu150 and Arg181 (**Suppl. Figure 11**). In two simulations with a peptide starting orientation PO1, Leu-Ala_pose1-run1,_ and His^0^-Ala_pose1-run2_, the corresponding peptides diffused from the binding cavity (**Suppl. Figure 10**). For the substrate His-Ala, it is apparent that in its protonated state, the histidine side chain is in close proximity to residue Asp60, though in two simulations, the C-termini of the peptides lost their interaction with Arg181 (**Suppl. Figure 10**). On the other hand, the neutral His-Ala peptide displays more flexibility of the histidine side chain in the binding site, while the peptide remains sandwiched between Arg181 and Glu150 (**Suppl. Figure 10**). In two simulations, the dipeptide Lys-Ala forms additional interactions with residues Tyr84 and Tyr87, either through its lysine side chain or the C-terminus of the peptide (**Suppl. Figure 10**).

Based on these simulation data in conjunction with the Cryo-EM data, we hypothesize that the orientation of the peptide at the end of the MD simulation from PO2 (**Figure 6F, Suppl. Figure 11**) represents the most likely binding mode of these dipeptides. In comparison to other peptide-bound structures of the well-studied members of the POT family, namely PepT1 or DtpB (Kotov et al., 2023), it is striking that MFSD1 displays a similar recognition pattern, even though MFSD1 does not share any of the POT signature motifs or their coupling mechanism (**Suppl. Figure 11 and 12**).

To validate our peptide recognition and transport findings, we performed mutagenesis experiments on selected highly-conserved peptide-binding site residues (**Figure 6G**), with Gln176 showing greater variability among different organisms (**Suppl. Figure 12**). Most mutants, except for MFSD1_E150R_, MFSD1_W373F,_ and MFSD1_Y56F_, could be expressed, solubilized, and purified (**Suppl. Figure 13A, B**). Peak fractions of the remaining mutants were used for subsequent nanoDSF experiments and liposome-based transport assays (**Figure 6G, Suppl. Figure 13C**). MFSD1_D60A_, MFSD1_E150A_, MFSD1_R181A,_ and MFSD1_R181E_ did not exhibit a characteristic thermal unfolding trace and thus could not be used to analyze peptide binding (**Suppl. Figure 13D, E**). MFSD1_W373A_ had an overall higher melting temperature (T_m_=46.6 °C) than MFSD1_WT_ (T_m_=40 °C), which did not increase upon peptide addition (**Suppl. Figure 13D, E**). The remaining mutants could still interact with dipeptides (**Suppl. Figure 13D, E**). Alterations in the stabilization pattern of selected peptides compared to MFSD1_WT_ occurred for MFSD1_Q176K_, where no stabilization by Pro-Arg, Arg-Pro, or Lys-Val was measured (**Suppl. Figure 13D, E**). Based on these results, MFSD1_Y56A_, MFSD1_D60A_, MFSD1_E150A_, MFSD1_Q176K_, MFSD1_R181A,_ and MFSD1_W373A_ were selected for liposome-based uptake assays of His-Ala and His-Ser. Most MFSD1 mutants lost their transport activity (**Figure 6G**). For MFSD1_Y56A,_ transport of His-Ala and His-Ser was still detectable, although the signal was reduced by ∼50% compared to MFSD1_WT_ (**Figure 6G**). While MFSD1_Q176K_ was still able to bind peptides, it did not transport them. Residue Gln176 is close to the potential ligand density identified in the Cryo-EM map of GLMP-MFSD1_His-Ala_ (**Figure 6D**) but is oriented away from the peptides screened in MD simulation experiments (**Figure 6F, Suppl. Figure 10**). Nevertheless, this residue is likely crucial for the transport mechanisms and, to a lesser extent, for peptide binding. As expected, mutating D60, E150, and R181 had greater implications on the stability of the protein and its ability to transport peptides, implying that these residues are critical for the interaction of the dipeptide with MFSD1 (**Figure 6E, G, Suppl. Figure 10, Suppl. Figure 13D, E**) as evident by the MD data (**Figure 6F, Suppl. Figure 10, 11**). The putative transport cycle model is shown in **Figure 6H**.

### Gating mechanism of MFSD1 using conformational predictions

The transition from the outward-open to the inward-open state is crucial for substrate translocation across the lysosomal membrane. For MFS transporters, the alternating access to the binding site is mediated by the movement of the N- and C-domains against each other, also known as the rocker-switch model (Drew et al., 2021; Quistgaard et al., 2016). Though the experimental structure of MFSD1 represents the outward-open state only, we made use of two additional conformations (representing the inward-open (in) and outward-occluded (occ) state) derived from AlphaFold2 (Tunyasuvunakool et al., 2021) predictions (**Suppl. Figure 14**). This allowed us to analyze the conformational transitions on a molecular level occurring during a transport cycle. Therefore, we aligned the N- and C-domains of the outward-occluded and inward-open models to the outward-open cryo-EM structure, termed MFSD1_out_. Overall, the two domains do not differ greatly when superimposed individually onto the N- or C-domain of MFSD1 (RMSD_Cα_ range of 0.85-1.27 Å, **Suppl. Figure 14A, B, Suppl. Video S1**). However, the superposition of the full-length proteins (RMSD_Cα_(out- occ)=3.5 Å, RMSD_Cα_(out-in)=4.87 Å) revealed that the N-domain undergoes a larger helical rearrangement in both predicted states, compared to the C-domain (**Suppl. Figure 14C, D, E, F**). During the transition from the outward-open to the outward-occluded state, the N-domain folds onto the substrate cavity, thereby closing it off from the lysosomal lumen, while the cytoplasmic bottom of the transporter stays relatively static (**Suppl. Figure 14G, H**). The cytoplasmic gate of MFSD1_out_ is formed by residues Asn57 (TM4), Phe173 (TM5), Trp373 (TM10), and Tyr389 (TM11) (**Suppl. Figure 14G, H**). Interestingly, the mutation of Trp393 residue to Alanine stabilized MFSD1 but interfered with peptide binding (**Suppl. Figure 13D, E**) and transport (**Figure 6F**). Further interactions between the N- and C-domain retained by Glu150 with Asn397, Arg181 with Tyr365, and a pi-cation interaction of Lys287 with Phe378 on the cytoplasmic side stabilize the outward-open conformation (**Suppl. Figure 14I**).

Interactions on the cytoplasmic side are similar between the outward-occluded AF2 model and MFSD1_out_. However, access to the binding cavity from the lysosomal lumen is blocked by residues Tyr59 (TM1), Asp60 (TM1), Met81 (TM2), Tyr84 (TM2), Ile283 (TM7), and Tyr309 (TM8), forming the lysosomal gate. During the transition from the outward-occluded state to the inward-open state, the cytoplasmic gate opens by a swinging motion of the bottom half of the N-domain away from the C-domain (**Suppl. Figure 14J**). This disrupts the cytoplasmic gate to open the cavity and thus facilitates the release of the substrate. The lysosomal gate remains closed and consists of the same residues as observed in the outward-occluded state (**Suppl. Figure 14H, J**). The conformation is further stabilized through interactions between the side chain of Lys287 with the backbone carbonyl of Ala64 and between the Gln66 side chain and the backbone amide of Val288 (**Suppl. Figure 14I, J**). Based on the analysis between the experimentally determined outward-open structure and the two AF2 models in the occluded and inward-open states, it becomes apparent that the positions of the crucial peptide binding residues Glu150 and Arg181 move towards the cytoplasmic side (**Suppl. Figure 14K**) and thus might push the dipeptide coordinated between both residues towards the cytoplasmic opening of MFSD1 to facilitate substrate release.

### The interaction of GLMP with MFSD1

Previous *in vivo* studies highlighted that GLMP is crucial for the protection of the transporter from degradation by lysosomal proteases and possibly crucial for the correct trafficking of MFSD1 to the lysosomal membrane (Lopez et al., 2020; Massa Lopez et al., 2019). Our data show that GLMP and MFSD1 form a stable complex even after detergent extraction from the cellular membrane and also during MD simulations (**Suppl. Figure 6A, 16A-D**). Based on the analysis of the Cryo-EM structure of the GLMP-MFSD1 complex, we identified a loop region of GLMP (residues 250-263) in close proximity to the luminal region of the C-domain of MFSD1. This region seems to be pivotal for the interaction between both proteins (**Figure 7A, B**) and, interestingly, was not resolved in the X-ray structure of GLMP (**Suppl. Figure 9C**), though it is conserved in other GLMP homologs (**Suppl. Figure 15**). The electrostatic surface of MFSD1 in this region is mainly positively charged, while it is negative for GLMP, indicating that both areas interact via polar interactions (**Figure 7B**). Arg292(GLMP) is in hydrogen bonding distance with Tyr416(MFSD1), and the loop is further stabilized by an intra-loop interaction of Asp256(GLMP) with the backbone amide of Ala261(GLMP) (**Figure 7C**). To evaluate the functional role of this interaction in a cellular context, we used *Glmp* knockout mouse embryonic fibroblasts, in which endogenous MFSD1 is strongly reduced, and the remaining MFSD1 is quantitatively localized to the Golgi apparatus. Ectopic re-expression of GLMP rescues this phenotype, and MFSD1 localizes to lysosomes again after GLMP re-transfection (Massa Lopez et al., 2019). We exchanged the entire interaction- surface loop region in influenza hemagglutinin (HA)-tagged GLMP (253 – 263) with four alanine residues and additionally generated constructs with individual amino acid exchanges (E250A, D256A, R292A) to test if these constructs can still rescue the lysosomal localization of MFSD1 and thus still interact (**Figure 7D**). HA-tagged LAMP1 (with the same type I topology and a similar high degree of N-glycosylation) was used as a negative control. Re-expression of WT GLMP efficiently restored the levels and localization of endogenous MFSD1 in *Glmp* knockout MEFs. In contrast, the re-expression of a construct coding for the variant with the deleted interaction-surface loop did not restore lysosomal MFSD1 localization. From the three tested point mutants, two (Glu250Ala and Arg292Ala) fully restored lysosomal MFSD1, while Asp256Ala did not, indicating that this amino acid is most critical in the interaction between MFSD1 and GLMP (**Figure 7D**). These data confirm the interaction surface between MFSD1 and GLMP located in the loop between 250 – 263 in an *in vivo* context.

**Figure 7.**
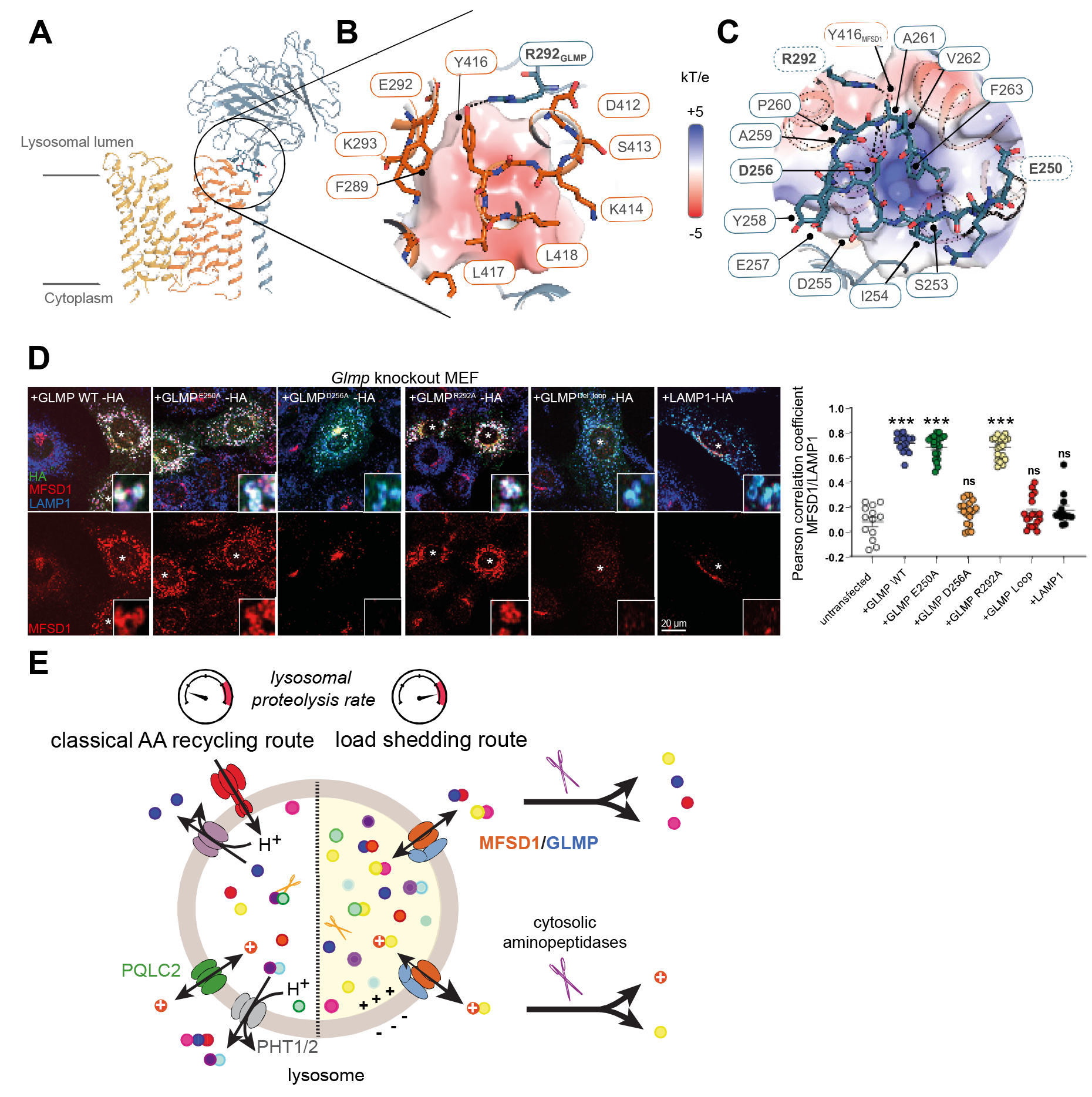
Interaction of GLMP with MFSD1. **(A)** Cartoon representation of GLMP in complex with MFSD1. The interaction site of GLMP with MFSD1 is highlighted in stick representation. **(B)** Zoom in on the interaction of MFSD1 to GLMP as viewed from MFSD1. The electrostatic surface of GLMP is shown. Y416 (MFSD1) is at a salt-bridge distance from R292 (GLMP) and is highlighted as a black dotted line. **(C)** Zoom in on the interaction of GLMP to MFSD1 as viewed from GLMP. The electrostatic potential surface of MFSD1 is highlighted, indicating complementarity to the GLMP surface. Besides the salt bridge between residue Y416 (MFSD1) and R292 (GLMP), residue D256 (GLMP) is at an H-bond distance from the backbone amide of A261 (GLMP) shown as black dotted lines. The loop region spanning residues 253 to 260 was mutated (blue border). Single-point mutants are highlighted in bold. **(D)** Immunofluorescence-staining of endogenous MFSD1 (red) after transfection with HA-tagged GLMP, GLMP mutants, and LAMP1 (green) in *Glmp* knockout MEFs. Endogenous LAMP1 is shown in blue. Transfected cells are marked with an (*). The Pearson correlation coefficient for MFSD1/ endogenous LAMP1 is shown in the right panel. Two-tailed unpaired t-tests: ns = not significant; * p ≤ 0.05; ** ≤ 0.01; *** p ≤ 0.001. **(E)** Cellular model for the role of MFSD1 in the recycling of amino acids derived from lysosomal proteolysis. Owing to its broad selectivity and low affinity for dipeptides, MFSD1 provides an alternative recycling route when lysosomal breakdown of proteins exceeds the capacity of lysosomal amino acid exporters. Fast cleavage of the released dipeptides by cytosolic aminopeptidases drives MFSD1 activity in the export direction and provides amino acids for biosynthetic pathways. The narrow selectivity of MFSD1 for dipeptides (in contrast with PHT1 and PHT2 transporters) prevents competition by single amino acids and protects this load shedding route from the amino acid overload.

## DISCUSSION

In recent years, the export of amino acids resulting from lysosomal proteolysis has received increasing attention (Wolfson and Sabatini, 2017). Several of the underlying transporters have been identified (Adelmann et al., 2020; Jezegou et al., 2012; Liu et al., 2012; Verdon et al., 2017; Wyant et al., 2017), although many of them are still missing. Various regulatory mechanisms of lysosomal amino acid transport have been discovered (Abu-Remaileh et al., 2017; Bandyopadhyay et al., 2022; Leray et al., 2021), and transporter structures have been characterized (Guo et al., 2022; Lei et al., 2018; Lobel et al., 2022). In contrast, lysosomal peptide transporters have received much less attention, although it has been known for decades that in lysosomes, specific peptides are not completely proteolytically degraded to single amino acids and that lysosomal peptide transporters must exist (Bird and Lloyd, 1990; Coffey and De Duve, 1968; Isenman and Dice, 1993; Thamotharan et al., 1997). Two members of the POT family, PHT1 (SLC15A4) and PHT2 (SLC15A3) localize to lysosomes and endosomes (Bockman et al., 1997; Kobayashi et al., 2014; Nakamura et al., 2014) and have been experimentally shown to transport carnosine, MDP, tri-DAP, Gly-Sar by PHT1 and His-Leu by PHT2 (Dong et al., 2023; Oppermann et al., 2019; Sakata et al., 2001; Wang et al., 2018). Additionally, both are thought to transport histidine, though evidence of uptake using different cell lines varied greatly (Bhardwaj et al., 2006; Lindley et al., 2011; Song et al., 2018; Yamashita et al., 1997). However, they are expected to transport a broader spectrum of di- and tripeptides due to their close relationship to the extensively studied bacterial and mammalian POT members.

In this study, we identify a novel lysosomal peptide transporter, MFSD1, and provide compelling evidence that it functions as a general, low-affinity uniporter for dipeptides. Some cationic dipeptides strongly accumulated in MFSD1-deficient lysosomes, providing a clue to elucidate the activity of this orphan lysosomal transporter. Studies of purified MFSD1 and of the MFSD1/GLMP complex expressed in a cell model showed that MFSD1 binds and efficiently transports diverse cationic or neutral dipeptides but not single amino acids nor longer peptides. Moreover, our combined cryo-EM and MD simulation data provided a structural basis for this substrate selectivity since a highly conserved glutamate (Glu150) and

arginine (Arg181) residue clamps the N- and C-termini, respectively, of the dipeptide in an extended conformation. The substrate binding site of MFSD1 thus acts as a “molecular ruler’’ that dictates the selectivity for dipeptides over longer peptides or single amino acids while accommodating diverse side chains, thus accounting for MFSD1’s promiscuity among dipeptides. This binding mode is reminiscent of the POT family (Kotov et al., 2023; Lyons et al., 2014; Martinez Molledo et al., 2018a), although MFSD1 lacks any of the typical POT signature motifs. A similar molecular ruler principle applied to cystine, the oxidized form of cysteine, instead of peptides, underlies the narrow substrate selectivity of cystinosin, the lysosomal transporter defective in cystinosis (Guo et al., 2022; Lobel et al., 2022).

When considered from a lysosomal physiology perspective, MFSD1 differs from PHT1 and PHT2 in several respects. First, MFSD1 is ubiquitously expressed (Massa Lopez et al., 2019), whereas the expression of PHT1 and PHT2 strongly varies across mammalian organs and tissues (Dong et al., 2023). Second, it has a strict selectivity towards dipeptides, as discussed. Third, MFSD1 has a low affinity for its substrates in the mM range, whereas PHT1 and PHT2 operate in the 10 to 100 µM range (Custodio et al., 2023; Dong et al., 2023). Therefore, lysosomal export of dipeptides by MFSD1 may intervene when there is a build-up of intralysosomal dipeptides, for instance, when cathepsin C, which has dipeptidyl peptidase activity (Turk et al., 2001), is more active or, more generally when the overall endopeptidase activity of the lysosomal lumen exceeds its exopeptidase activity.

Finally, MFSD1 differs from POT family members and from many lysosomal transporters by its bioenergetical properties. Although its transport rate is accelerated by an acidic lumen, it is not driven by the proton electrochemical gradient, as shown by the absence of intrinsic proton coupling in our electrophysiological recordings. Luminal protons (extracellular protons in our topologically inverted oocyte assay) were co-transported exclusively with a subset of substrates (His-containing dipeptides, Glu-Lys) harboring a side chain pK_a_ relatively close to the pH of our extracellular medium, but not with a substrate like Lys-Ala (side chain pK_a_ = 10.5). These protons are thus carried by the protonatable side chain (His, Glu) of the former substrates rather than through an MFSD1 proton pathway.

MFSD1 is thus a uniporter. This lack of ion coupling implies that the transport activity of MFSD1 can easily reverse in the lysosomal membrane, in contrast with the proton-coupled lysosomal transporters, which are powerfully driven in the export direction by the steep pH gradient of the lysosome. Of note, this propensity of MFSD1 to reverse direction may help understand the old paradoxical observation that high concentrations of dipeptide enter and burst purified lysosomes more efficiently than single amino acids (Bird and Lloyd, 1990).

Our data show that two other forces drive MFSD1 in the export direction in a cellular context (**Figure 7E**). The first major one is the powerful hydrolysis of dipeptides by cytosolic

aminopeptidases (Botbol and Scornik, 1989; Bouma et al., 1976), as highlighted in our tracer experiments by the full cleavage of Leu(*d3*)-Ala to Leu(*d3*) and Ala after its discharge into the cytosol. Another driving force, which is restricted to cationic dipeptides, is the positive-inside polarization of the lysosomal membrane (Koivusalo et al., 2011; Saminathan et al., 2021). This polarization selectively accelerates the lysosomal export of cationic dipeptides, presumably explaining why this dipeptide subclass specifically stood out in our initial metabolomics profiling of candidate substrates.

Taken together, these features (ubiquitous expression; low affinity; uniporter activity driven by a fast cytosolic cleavage) strongly suggest that MFSD1 provides an alternative route to supply amino acids for biosynthetic pathways when the ‘classical’ route mediated by lysosomal amino acid transporters and, in some cell types, PHT1 and PHT2 is overloaded (**Figure 7E**). Moreover, the narrow selectivity of MFSD1 for dipeptides protects this load-shedding route from competition by single amino acids or longer peptides.

Our structural study also unveils the molecular architecture of the MFSD1/GLMP hetero-oligomer, and it suggests a putative mechanism for the transport of dipeptides (**Figure 6H**). By combining experimental structural work with AF2 models and MD simulations, we propose that dipeptide binding is mainly facilitated by the coordination of its N- and C-termini via the highly conserved Glu150 and Arg181 residues. We assume that peptide binding induces the helical rearrangement of TM1, TM2, TM7, and TM8, as observed in other MFS-fold-type systems. The closure of the lysosomal gate results in the displacement of Glu150 and Arg181 towards the cytoplasmic part of the binding site, which in turn pushes the dipeptide downwards. Additionally, if the substrate had remained in its initial position, it would lead to a steric clash with Tyr59, which undergoes an inward rotation when closing the binding site to the lysosomal lumen. To release the substrate to the cytoplasm, helices TM4, TM5, TM10, and TM11 need to be rearranged, resulting in a rotation of the cytoplasmic gating residues away from each other and thus opening the binding site to the cytoplasm.

MFSD1 forms a tight complex with GLMP, and under physiological conditions, the stability of both proteins fully depends on the other interaction partner, as shown by an almost complete co-deficiency of each protein in tissues and cells of *Mfsd1* or *Glmp* knockout mice, respectively (Massa Lopez et al., 2019). Additionally, there is evidence of a partial interdependence of MFSD1 and GLMP in their intracellular trafficking to lysosomes in mammalian cells (Lopez et al., 2020). This interdependence is also highlighted in our oocyte experiments, in which only the co-expression of MFSD1 together with GLMP (both mutated in their lysosomal sorting motifs) led to detectable MFSD1 at the plasma membrane and, accordingly, a transport current. This, again, suggests a function as reciprocal chaperones or

stabilizers. Under these conditions, the system did not allow for the analysis of the question of how GLMP affects the substrate translocation activity of MFSD1. Our *in vitro* liposome reconstitution experiments, however, allowed a direct comparison of the MFSD1 activity alone or with GLMP either as a fusion construct or in complex. The reconstituted complex of GLMP and MFSD1 exhibited similar uptake rates compared to MFSD1_WT_ only, whereas the transport activity for the fusion protein was reduced. This is probably due to the linker approach used to connect both proteins, which has been beneficial for Cryo-EM studies but reduced the conformational flexibility crucial for transport activity. Our Cryo-EM data revealed a crucial loop within GLMP interacting with the lysosomal surface of the C-terminal domain of MFSD1, mainly stabilized by polar interaction between GLMP and MFSD1. Mutagenesis confirmed the importance of this loop region of GLMP since mutations in this interaction surface could not rescue MFSD1. The structure of MFSD1 and GLMP in the complex illustrates that the highly N-glycosylated GLMP shields the luminal loops and the surface of the unglycosylated MFSD1 from proteases, supporting the presumed function as a

„protector“ similar to OSTM1 for the lysosomal chloride channel CLCN7 (Kornak et al., 2001; Schrecker et al., 2020). Interestingly, our MD simulations of the GLMP-MFSD1 complex, compared to MFSD1 alone, revealed the presence of a POPE molecule close to the peptide binding site, but the role of this lipid remains unclear (**Suppl. Figure 15**). However, we did observe an additional density between the N- and C-terminal bundle of MFSD1 in the Cryo-EM data, which could possibly correspond to a lipid or detergent molecule.

## DATA AVAILABILITY

The EM data and fitted models for GLMP-MFSD1 have been deposited in the Electron Microscopy Data Bank under accession code EMD-19005 and the PDB under accession code 8R8Q. The crystal structure of GLMP used for comparative analysis in this study can be found in the PDB under accession code 6NYQ AlphaFold2 predictions of MFSD1 as well as the models of MFSD1 and GLMP-MFSD1 after 300 ns of molecular dynamics simulations were deposited to Zenodo (Alphafold models: 10.5281/zenodo.10276738; MFSD1 apo/with ligands in initial poses and after 300 ns MD: 10.5281/zenodo.10276760). All protein sequences used in this study are publicly available at Uniprot (https://www.uniprot.org/)

## ACKNOWLEDGEMENT

Sebastian Held is acknowledged for excellent technical assistance. We thank the BioMedTech animal facility at Université de Paris (CNRS UMS2009, INSERM US36) for housing the frogs and the "Service de spectrométrie de masse de l’UMR8601" for mass spectrometry analysis. BioRender was used to create parts of the figures. M.A.-R. is a Stanford Terman Fellow and a Pew-Stewart Scholar for Cancer Research, supported by the Pew Charitable Trusts and the Alexander and Margaret Stewart Trust. K.E.J.J. and V.K. were supported by a research fellowship from the EMBL interdisciplinary postdoctoral (EIPOD) program under Marie Curie Cofund Actions MSCA-COFUND-FP (grant agreement 847543).We thank the Sample Preparation and Characterisation facility of EMBL Hamburg and the Cryo-EM facility of CSSB for their support, technical assistance, and advice. The groups of Henning Tidow, Charlotte Utrecht, and Holger Sondermann are acknowledged to grant access to their equipment necessary for this project. We thank Maxime Killer, Siavash Mostafavi, and Grzegorz Chojnowski for their scientific input on Cryo-EM data processing and Josi Steinke for their additional assistance in liposome reconstitution experiments.

## AUTHOR CONTRIBUTIONS

M.D., B.G., C.L., O.G., K.E.J.J., and M.AR. designed the study. M.D., O.G., K.E.J.J., M.I.,C.D., K.S., D.M.L., J.D., S.M., V.K., F.G., A.E. I.MT, N.N.L and S.H.C performed the experiments. M.D., B.G., C.L., O.G., K.E.J.J., W.D., V.K., F.G., and M.AR. analyzed the data and/or its significance. M.D., B.G., C.L., O.G., and K.E.J.J. wrote the manuscript with contributions from W.D. and P.S.; M.D., B.G., and C.L. acquired funding.

## SUPPLEMENTAL INFORMATION

Supplemental information can be found online at XXX.

### ABBREVIATIONS

AF: AlphaFold
DSF: Differential scanning fluorimetry
EM: Electron microscopy
PNS: Postnuclear supernatant
POPE: 1-palmitoyl-2-oleoyl-sn-glycero-3-phosphoethanolamine
POPG: P1-palmitoyl-2-oleoyl-sn-glycero-3-phospho-1’-rac-glycerol
MD: Molecular dynamics
MFS: Major Facilitator Superfamily
NMR: Nuclear magnetic resonance
TM: Transmembrane

