## Supplemental Table 1 for "MFSD1 in complex with its accessory subunit GLMP functions as a general dipeptide uniporter in lysosomes"

**Supplemental Table 1. List of compounds for nanoDSF screening.**

|  | **ligand** | **category** | **company** | **order number** |
| --- | --- | --- | --- | --- |
| 1 | A | amino acid | Fluka | 5129 |
| 2 | C | amino acid | Calbiochem | 2470 |
| 3 | D | amino acid | Sigma | A6683 |
| 4 | E | amino acid | Sigma | G8415 |
| 5 | F | amino acid | Sigma | P2126 |
| 6 | G | amino acid | Aldrich | 241261 |
| 7 | H | amino acid | Sigma | H9386 |
| 8 | I | amino acid | Calbiochem | 4160 |
| 9 | K | amino acid | Sigma | L1262 |
| 10 | L | amino acid | Sigma | L8125 |
| 11 | M | amino acid | Sigma | M5308 |
| 12 | N | amino acid | Sigma | A0884 |
| 13 | P | amino acid | Sigma | P0380 |
| 14 | Q | amino acid | Sigma | G-3126 |
| 15 | R | amino acid | Roth | 3144.2 |
| 16 | S | amino acid | Sigma | S4311 |
| 17 | T | amino acid | Sigma | T8625 |
| 18 | V | amino acid | Sigma | V6504 |
| 19 | glucose | sugar | Roth | X997.2 |
| 20 | D-galactonate | sugar | Sigma | 44511 |
| 21 | maltose | sugar | Sigma | M-5885 |
| 22 | lactose | sugar | Roth | 8921.1 |
| 23 | L-galactonate | sugar |  | synthesised from  L-Galactono-1,4-lactone |
| 24 | AA | di-peptide | Sigma | A502 |
| 25 | AD | di-peptide | Bachem | G-1195 |
| 26 | AE | di-peptide | Bachem | G-1200 |
| 27 | AF | di-peptide | Bachem | A-3128 |
| 28 | AH | di-peptide | Bachem | G1245 |
| 29 | AI | di-peptide | Bachem | G1260 |
| 30 | AK | di-peptide | Bachem | G-1290 |
| 31 | AL | di-peptide | Bachem | A-1878 |
| 32 | AP | di-peptide | Bachem | G-1350 |
| 33 | AQ | di-peptide | Sigma | G8541 |
| 34 | AV | di-peptide | Bachem | G-1320 |
| 35 | beta-Ala-beta-Ala | di-peptide | Bachem | G-1150 |
| 36 | AR | di-peptide | Bachem | G-1175 |
| 37 | DA | di-peptide | Bachem | G-1550 |
| 38 | DE | di-peptide | Sigma | A1916 |
| 39 | EA | di-peptide | Bachem | G-1900 |
| 40 | EE | di-peptide | Sigma | G-3640 |
| 41 | EK | di-peptide | Bachem | G-1955 |
| 42 | ES | di-peptide | Bachem | G-1980 |
| 43 | GG | di-peptide | Sigma | 50199 |
| 44 | GS | di-peptide | Sigma | G-3127 |
| 45 | HA | di-peptide | Bachem | G-2285 |
| 46 | HD | di-peptide | Bachem | G-2295 |
| 47 | HE | di-peptide | Bachem | G-2300 |
| 48 | HH | di-peptide | Bachem | G4595 |
| 49 | HK | di-peptide | Bachem | G2320 |
| 50 | HS | di-peptide | Sigma | H3129 |
| 51 | KA | di-peptide | Bachem | G-2630 |
| 52 | KK | di-peptide | Bachem | G-2675 |
| 53 | KP | di-peptide | Bachem | G-4190 |
| 54 | KV | di-peptide | Bachem | G-2705 |
| 55 | LA | di-peptide | Bachem | G-2460 |
| 56 | LL | di-peptide | Bachem | M-1535 |
| 57 | MS | di-peptide | Sigma | M9380 |
| 58 | MT | di-peptide | Bachem | G-2810 |
| 59 | NV | di-peptide | Bachem | G-3720 |
| 60 | PA | di-peptide | Bachem | G-2985 |
| 61 | PR | di-peptide | Bachem | G-2990 |
| 62 | RA | di-peptide | Bachem | G-4170 |
| 63 | RF | di-peptide | Bachem | G-1515 |
| 64 | RP | di-peptide | Bachem | G-3675 |
| 65 | SH | di-peptide | Bachem | G-3195 |
| 66 | SL | di-peptide | Bachem | G-3200 |
| 67 | TF | di-peptide | Bachem | G-3295 |
| 68 | AAA | tri-peptide | Bachem | H-1445 |
| 69 | AFA | tri-peptide | Bachem | H-5420 |
| 70 | AHA | tri-peptide | GL Biochem | Custom synthesis |
| 71 | AMA | tri-peptide | GL Biochem | Custom synthesis |
| 72 | APA | tri-peptide | Bachem | H-1595 |
| 73 | APF | tri-peptide | Bachem | H-1615 |
| 74 | AQA | tri-peptide | GL Biochem | Custom synthesis |
| 75 | ARA | tri-peptide | GL Biochem | Custom synthesis |
| 76 | ASA | tri-peptide | GL Biochem | Custom synthesis |
| 77 | FAD | tri-peptide | GL Biochem | Custom synthesis |
| 78 | FAT | tri-peptide | GL Biochem | Custom synthesis |
| 79 | GGG | tri-peptide | Sigma | G1377 |
| 80 | GGH | tri-peptide | Sigma | G4541 |
| 81 | LGG | tri-peptide | Sigma | L9750 |
| 82 | LLA | tri-peptide | Bachem | H-3905 |
| 83 | MAS | tri-peptide | Sigma | M1004 |
| 84 | PVG | tri-peptide | Bachem | H-4820 |
| 85 | Carnosine | other | Bachem | G-1250 |
| 86 | Gly-Sar | other | Sigma | G3127 |
| 87 | Anserine | other | Bachem | G-4555 |
| 88 | 5-aminolevulinic acid | other | Sigma | A3785 |
| 89 | valacyclovir | other | Sigma | PHR1601 |
| 90 | Penicillin | other | Sigma | 13752 |
| 91 | gemcitabine | other | N/A | N/A |
| 92 | GGGG | tetra-peptide | Bachem | H-3380 |
| 93 | AAAA | tetra-peptide | Bachem | H-1260 |
| 94 | 5 % DMSO |  | Sigma | 472301 |
| 95 | apo |  |  |  |
