## Supplemental Table 2 for "MFSD1 in complex with its accessory subunit GLMP functions as a general dipeptide uniporter in lysosomes"

**Supplemental Table 2. Selected ions monitored (SIM) and multiple reaction monitoring (MRM) and their retention times for** **LC-MS/MS analysis.**

| **Analyte** | **m/z** | **RT (min)** |
| --- | --- | --- |
| D3- Leucine (SIM) | 135.11 | 1.6 |
| D3-Leucine (MRM) | 135.10>89.10 | 1.6 |
|  | 135.10>46.15 |  |
|  | 135.10>44.15 |  |
| D3- Leucine-Alanine (SIM) | 206.15 | 4.3 |
| D3-Leucine-Alanine (MRM) | 206.15>89.15 | 4.3 |
|  | 206.15>46.15 |  |
|  | 206.15>44.15 |  |
| Alanine (SIM) | 89.90 | 1.2 |
| Alanine (MRM) | 89.90>44.10 | 1.2 |

RT = retention time
