## Supplemental Table 3 for "MFSD1 in complex with its accessory subunit GLMP functions as a general dipeptide uniporter in lysosomes"

**Supplemental Table 3. Description of the molecular dynamics simulations.**

|  | **Substrate** | **Protonation status** | **Length (ns)** |
| --- | --- | --- | --- |
| **Apo** | --- | --- | 3 ✕ 300 |
| **Apo+GLMP** | --- | --- | 3 ✕ 300 |
| **Lys-Ala** | Lys-Ala | N- and C- terminal charged | 3 ✕ 300 |
| **Leu-Ala** | Leu-Ala | N- and C- terminal charged | 3 ✕ 300 |
| **His-Ala** | His-Ala | N- and C- terminal charged  Histidine side chain neutral | 3 ✕ 300 |
| **His-Ala** | His-Ala | N- and C- terminal charged  Histidine side chain charged | 3 ✕ 300 |
