## Supplemental Table 4 for "MFSD1 in complex with its accessory subunit GLMP functions as a general dipeptide uniporter in lysosomes"

**Supplemental Table 4. Cryo-EM data collection, refinement, and validation statistics.**

|  | **GLMP-MSD1_apo_**  (EMDB: EMD-19005) | **GLMP-MSD1_His-Ala_**  (PDB: 8R8Q; EMDB: EMD-19006) |
| --- | --- | --- |
| **Data collection and processing** | | |
| Voltage [kV] | 300 | 300 |
| Magnification | 81,000 x | 81,000 x |
| Pixelsize [Å] | 1.1 | 1.1 |
| Exposure Time [s] | 4.2 | 4.2 |
| Total Dose (e^-^/A^2^) | 55 | 55 |
| Defocus Values [µm] | -1.0 to -3.0 (in 0.5 increments) | -1.0 to -3.0 (in 0.5 increments) |
| Micrographs | 3179 + 2551 | 3,193 |
| Initial particle images | 2,507,968 | 1,375,022 |
| Final particle images | 308,511 | 400,191 |
| Map resolution [Å] | 4.2 | 4.1 |
| FSC threshold | 0.143 | 0.143 |
| **Refinement** | | |
| Initial model used | - | AlphaFold2 GLMP + MFSD1 |
| Model composition | |  |
| Chains | - | 2 |
| Nonhydrogen atoms | - | 5,702 |
| Protein residues | - | 732 |
| Ligands | - | NAG: 5 |
| Average B factors [Å^2^] | | |
| Protein | - | 63.01 |
| Ligand | - | 103.01 |
| r.m.s.d. | |  |
| Bond lengths [Å] | - | 0.003 |
| Bond angles [°] | - | 0.646 |
| **Validation** | | |
| MolProbity score | - | 1.91 |
| Clashscore | - | 7.7 |
| Poor rotamers [%] | - | 0 |
| Ramachandran plot | | |
| Favored [%] | - | 92.08 |
| Allowed [%] | - | 7.92 |
| Outlier [%] | - | 0 |
