## Supplementary Figures for "MFSD1 in complex with its accessory subunit GLMP functions as a general dipeptide uniporter in lysosomes"

### Supplemental Figure 1

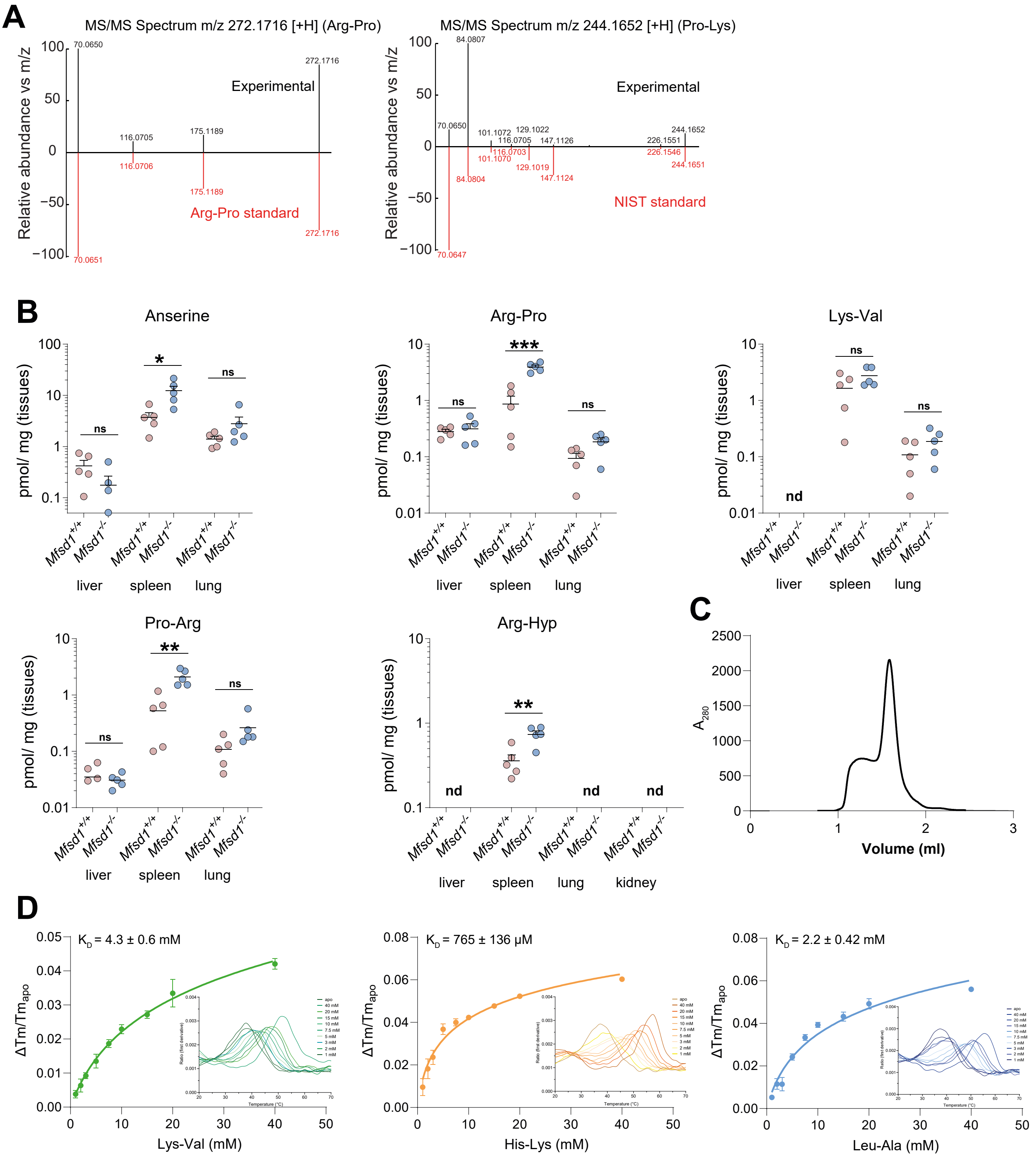


**Supplementary Figure 1. Validation of Arg-Pro and Pro-Lys with standards and dipeptide levels in tissues of *Mfsd1*$^{tm1d/tm1d}$ mice (A).** Mirror plots for the experimental MS/MS spectra of Arg-Pro (left) and Pro-Lys (right) and authentic chemical standards. The individual spectra for the experimentally determined metabolites are shown in black, and the spectra of the chemical standards are shown in red. **(B)** Quantification of the levels of the dipeptides anserine, Arg-Pro, Lys-Val, Pro-Arg, and Arg-Hyp in total tissue lysates of 6-month-old wildtype ad *Mfsd1* knockout mice. P-values were calculated using two-tailed paired t-tests. Error bars show the mean ± SEM. **(C)** SEC chromatogram of purified MFSD1 with a Streptavidin-tag. **(D)** $K_D$ measurements for Lys-Val, His-Lys, Leu-Ala. $K_D$ measurements are based on changes in the thermal stability of MFSD1 in the presence of varying concentrations of the dipeptides Lys-Val (green), His-Lys (orange), and Leu-Ala (blue). $K_D$ values were determined using Moltenprot (Kotov et al., 2021).

### Supplemental Figure 2

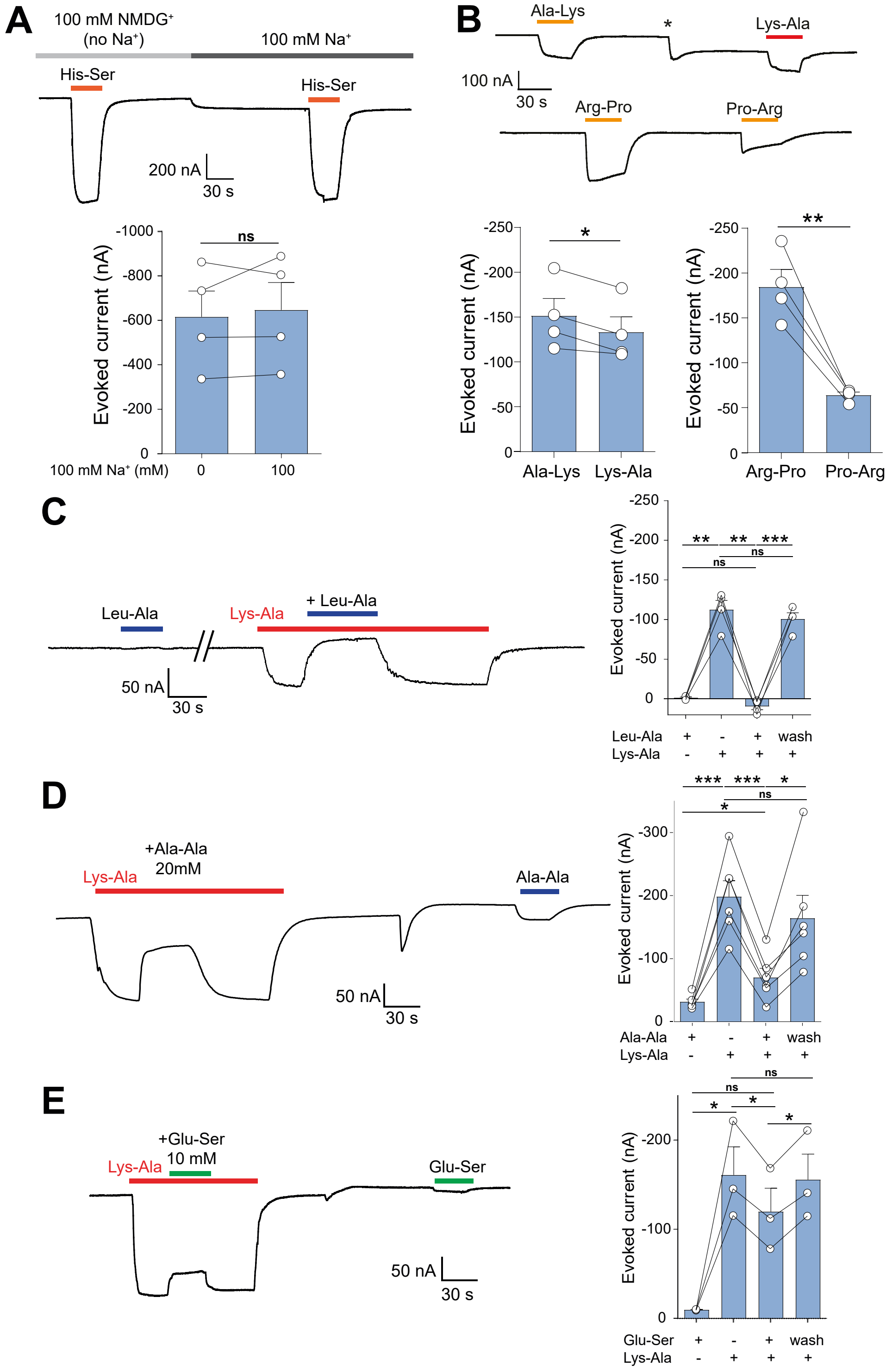


**Supplementary Figure 2. Dipeptide selectivity of MFSD1/GLMP in the TEVC oocyte assay. (A)** The MFSD1/GLMP transport current does not depend on sodium ions. His-Ser (10 mM) was applied to MFSD1/GLMP oocytes at pH 5.0 in the presence of $Na^+$ or $NMDG^+$ as the major cation. P values were calculated using two-tailed paired t-tests. **(B)** Residue order effect for two cationic dipeptides. Representative traces and mean TEVC currents ± SEM of four MFSD1/GLMP oocytes. P-values were calculated using two-tailed paired t-tests. **(C, D)** Competition of the Lys-Ala current by neutral dipeptides. Lys-Ala (3 mM) and Leu-Ala or Ala-Ala (20 mM) were applied separately or simultaneously to MFSD1/GLMP oocytes at pH 5.0. Representative traces and mean currents ± SEM of four (Leu-Ala) and six (Ala-Ala) oocytes. **(E)** The competition experiment was repeated with the anionic dipeptide Glu-Ser (10 mM). P-values were calculated using two-tailed paired t-tests.

### Supplemental Figure 3

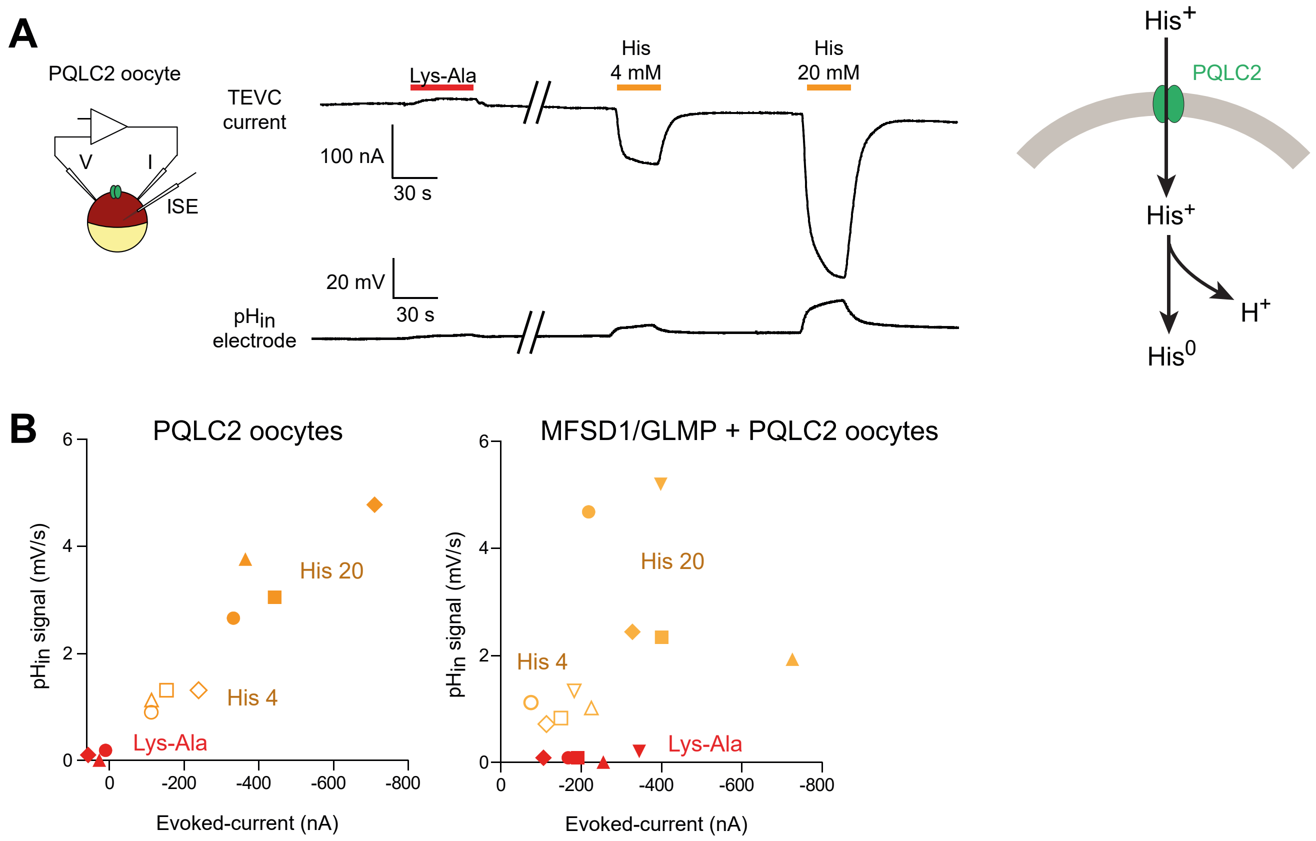


**Supplementary Figure 3. Additional evidence for the uniporter model. (A)** Combined TEVC and intracellular pH ($pH_{in}$) recording of oocytes expressing only the sorting mutant of PQLC2. Lys-Ala (20 mM) is not transported by PQLC2. The traces are representative of four PQLC2 oocytes. **(B)** The current/acidification relationship of the experiments is shown in **Figure 4A** and **Suppl. Figure 3A**. The graphs show individual TEVC and $pH_{in}$ responses to Lys-Ala (10 mM) and His (4 or 20 mM). Each symbol shape represents a distinct oocyte.

### Supplemental Figure 4

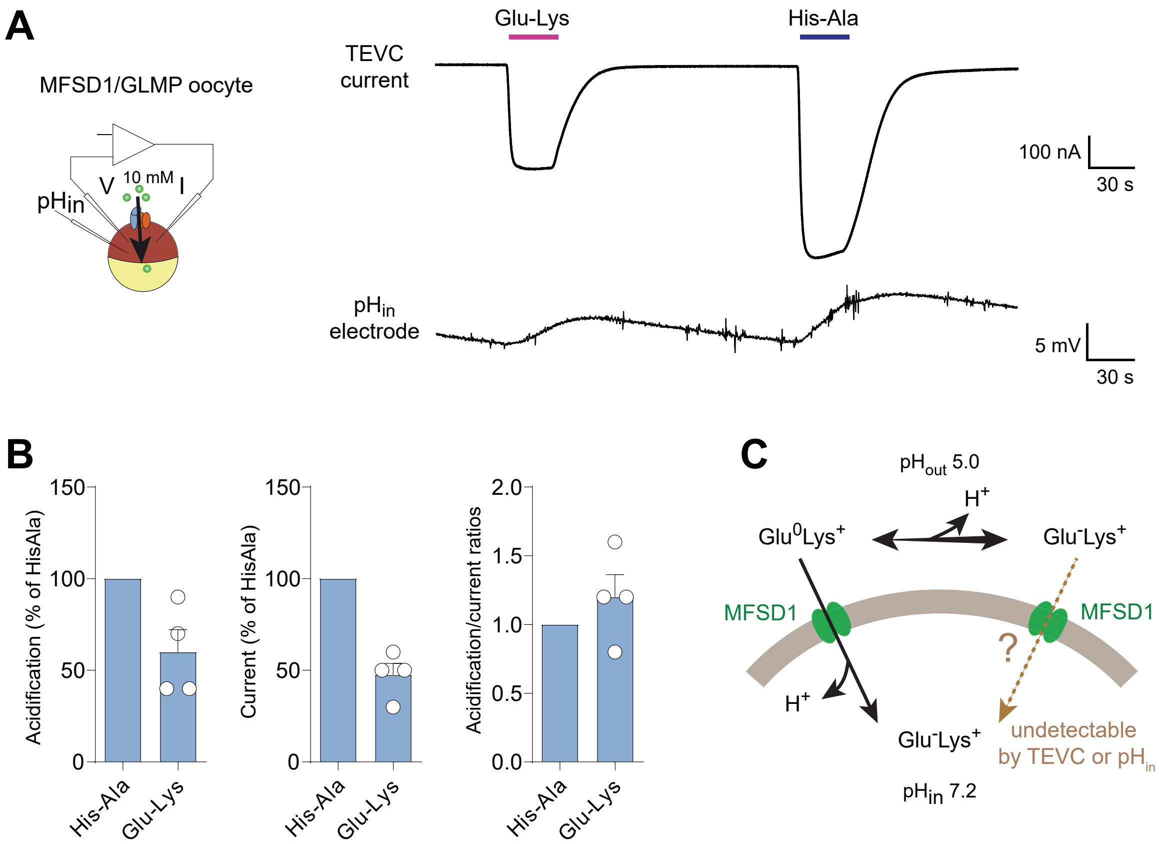


**Supplementary Figure 4. Uptake of Glu-Lys into MFSD1/GLMP oocytes. (A)** Representative TEVC and $pH_{in}$ traces of the response of MFSD1/GLMP oocytes to Lys-Glu **(B)** Acidification and current responses normalized to His-Ala and normalized acidification/current ratios. Data are means ± SEM of 4 oocytes. **(C)** A model accounting for the uptake of Glu-Lys by MFSD1/GLMP. Only uptake of the minor cationic form, $Glu^{0}$-$Lys^{+}$, can be detected in this electrophysiological technique.

#### Supplemental Figure 5

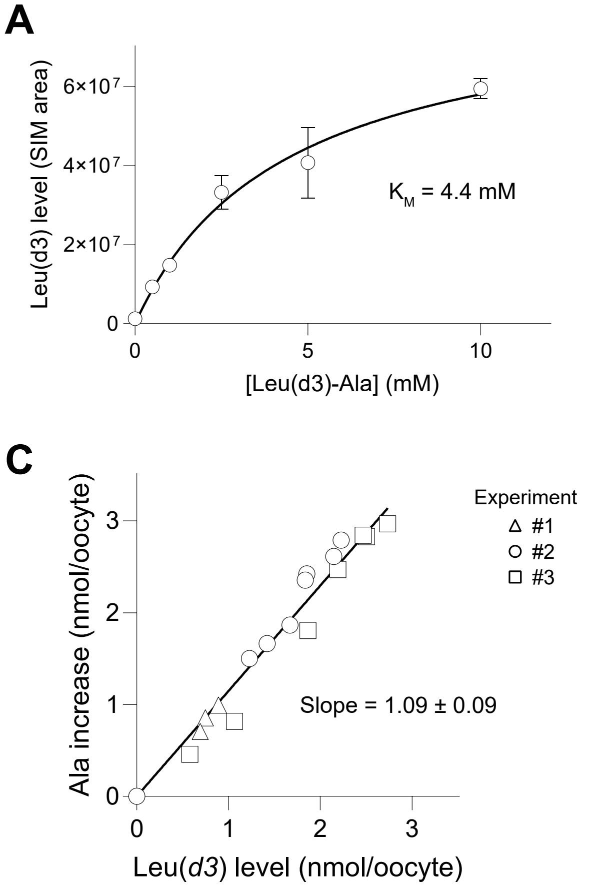


**Supplementary Figure 5. Additional evidence for Leu-Ala uptake by MFSD1. (A)** Dose-response relationship of the accumulation of Leu(*d3*) in MFSD1/GLMP oocytes exposed to Leu(*d3*)-Ala (means ± SEM of 3 oocytes). The line shows a hyperbolic curve fit with a $K_M$ value of 4.4 mM ($R^2$ = 0.989). **(B)** Time course of Leu(*d3*) accumulation in the presence of 10 mM Leu(*d3*)-Ala (means ± SEM of 3 oocytes). Linear regression $R^2$ = 0.980. **(C)** Relationship between the accumulation of Leu(*d3*) and the increase of 'light' Ala over its endogenous level. Data shown in **Figure 5D** were replotted to show the equimolar ratio between these two proxies of Leu(*d3*)-Ala uptake. Linear regression of the pooled data yielded a ratio of 1.15 Ala molecule co-released with each Leu(*d3*) molecule ($R^2$ = 0.980), or a mean ratio of 1.09 ± 0.09 when the 3 experiments were analyzed separately.

### Supplemental Figure 6

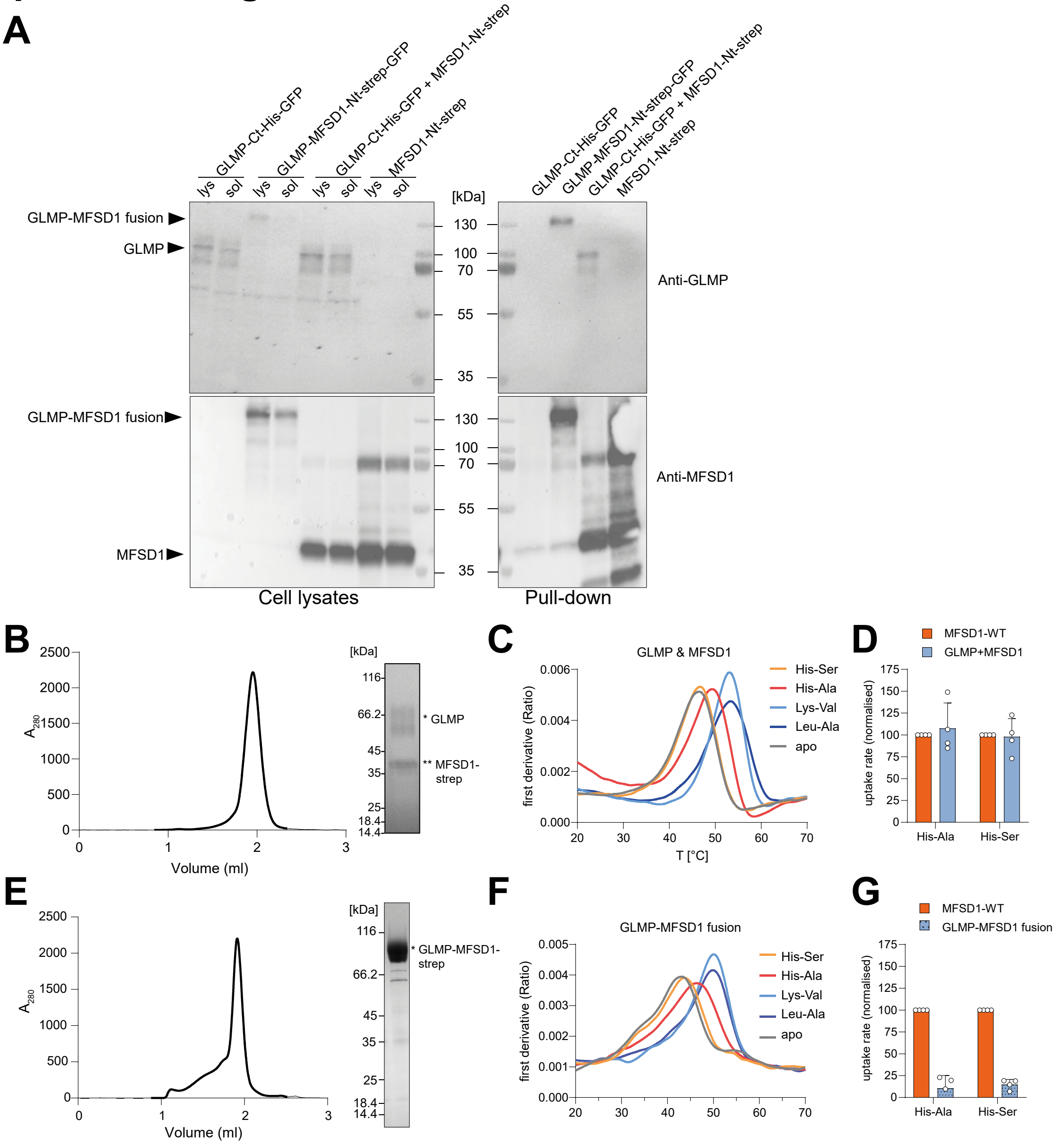


**Supplementary Figure 6. The recombinantly expressed proteins interact *in vitro*. (A)** Pull-down assays of Twin-Streptavidin (strep) tagged MFSD1 (MFSD1-Nt-strep), GFP-strep-tagged GLMP-MFSD1 (GLMP-MFSD1-Nt-strep-GFP) and GFP-8×His-tagged GLMP (GLMP-Ct-His-GFP) and GLMP-MFSD1-Nt-strep-GFP. Each protein was individually over-expressed in Expi293F, and additionally, MFSD1-Nt-strep was co-expressed with GLMP-Ct-His-GFP. MFSD1 and GLMP were detected in Western blot using specific primary antibodies against either MFSD1 or GLMP. Samples either contained crude lysate (lys) or the soluble fraction (sol) of each construct over-expressed in Expi293F cells (left panel) or the elution fraction after pull-down over Strep-Tactin beads (right panel). Bands corresponding to GLMP, GLMP-MFSD1, or MFSD1 are indicated. **(B)** SEC chromatogram and SDS-PAGE gel of purified GLMP in complex with MFSD1 carrying a twin-streptavidin-tag (MFSD1-strep). **(C)** Thermal stability of GLMP in complex with MFSD1-strep in the absence (apo) or presence of 5 mM His-Ser (HS), His-Ala (HA), Lys-Val (KV), or Leu-Ala (LA). **(D)** Normalized initial uptake rates of the dipeptides His-Ala or His-Ser during liposome-based assays by $MFSD1_{WT}$ and GLMP/MFSD1 complex. n=4, of two reconstitution batches; Error bars are shown as SD. **(E)** SEC chromatogram and SDS-PAGE gel of purified GLMP-MFSD1-fusion protein carrying a twin-streptavidin-tag. **(F)** Thermal stability of GLMP-MFSD1-fusion protein in the absence (apo) or presence of 5 mM His-Ser (HS), His-Ala (HA), Lys-Val (KV), or Leu-Ala (LA). **(G)** Normalized initial uptake rates of the dipeptides His-Ala or His-Ser during liposome-based assays by $MFSD1_{WT}$ and GLMP-MFSD1 fusion protein. n=4, Error bars are shown as SD.

#### Supplemental Figure 7

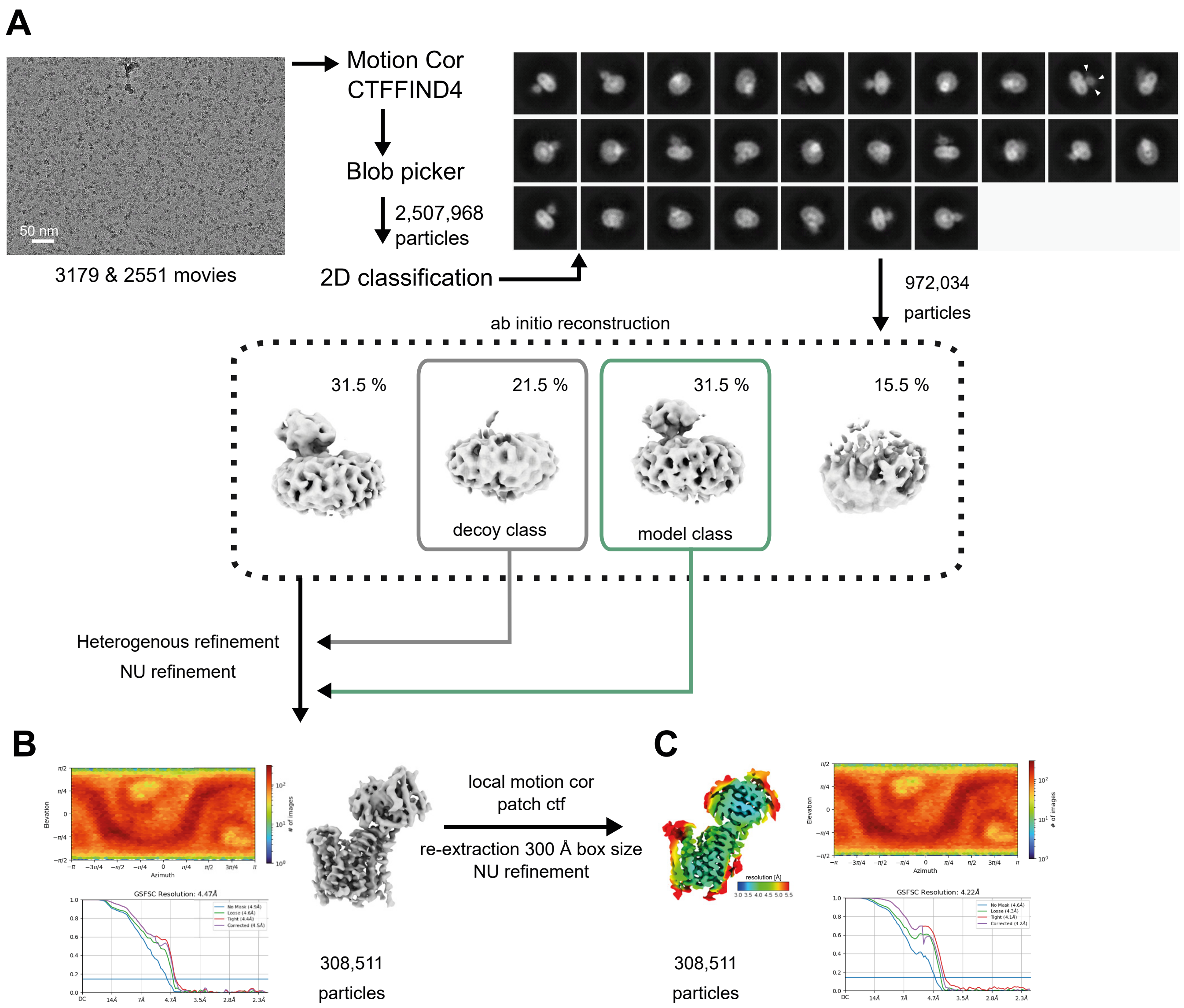


**Supplementary Figure 7. CryoEM data collection and processing of the GLMP-MFSD1$_{apo}$ data set. (A)** Image data processing workflow with a representative micrograph and 2D classes of the GLMP-MFSD1$_{apo}$ data set. All data were processed in cryoSPARC. **(B)** Angular distribution plot, GSFSC plot, and cryoEM map of initial reconstruction before further refinement. White arrowheads denote densities corresponding to N-glycans. **(C)** Angular distribution plot, GSFSC plot, and cryoEM map of final reconstruction colored by local resolution.

#### Supplemental Figure 8

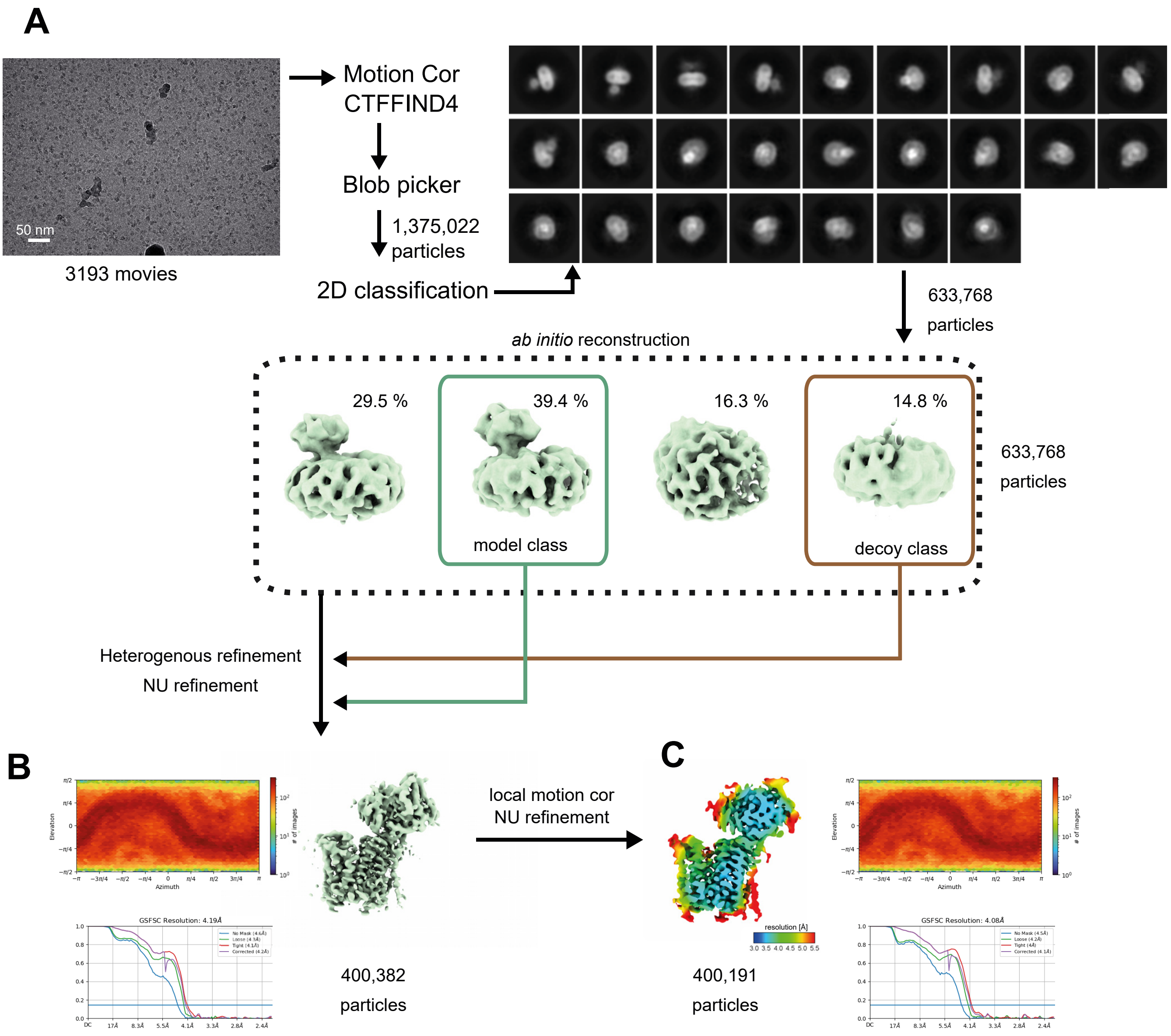


**Supplementary Figure 8. Cryo-EM data collection and processing of the GLMP-MFSD1$_{\text{His-Ala}}$ data set. (A)** Image data processing workflow with a representative micrograph and 2D classes of the GLMP-MFSD1$_{\text{His-Ala}}$ data set. All data were processed in cryoSPARC. **(B)** Angular distribution plot, GSFSC plot, and Cryo-EM map of initial reconstruction before further refinement. White arrowheads in 2D class references denote densities corresponding to N-glycans. **(C)** Angular distribution plot, GSFSC plot, and Cryo-EM map of final reconstruction colored by the local resolution.

### Supplemental Figure 9

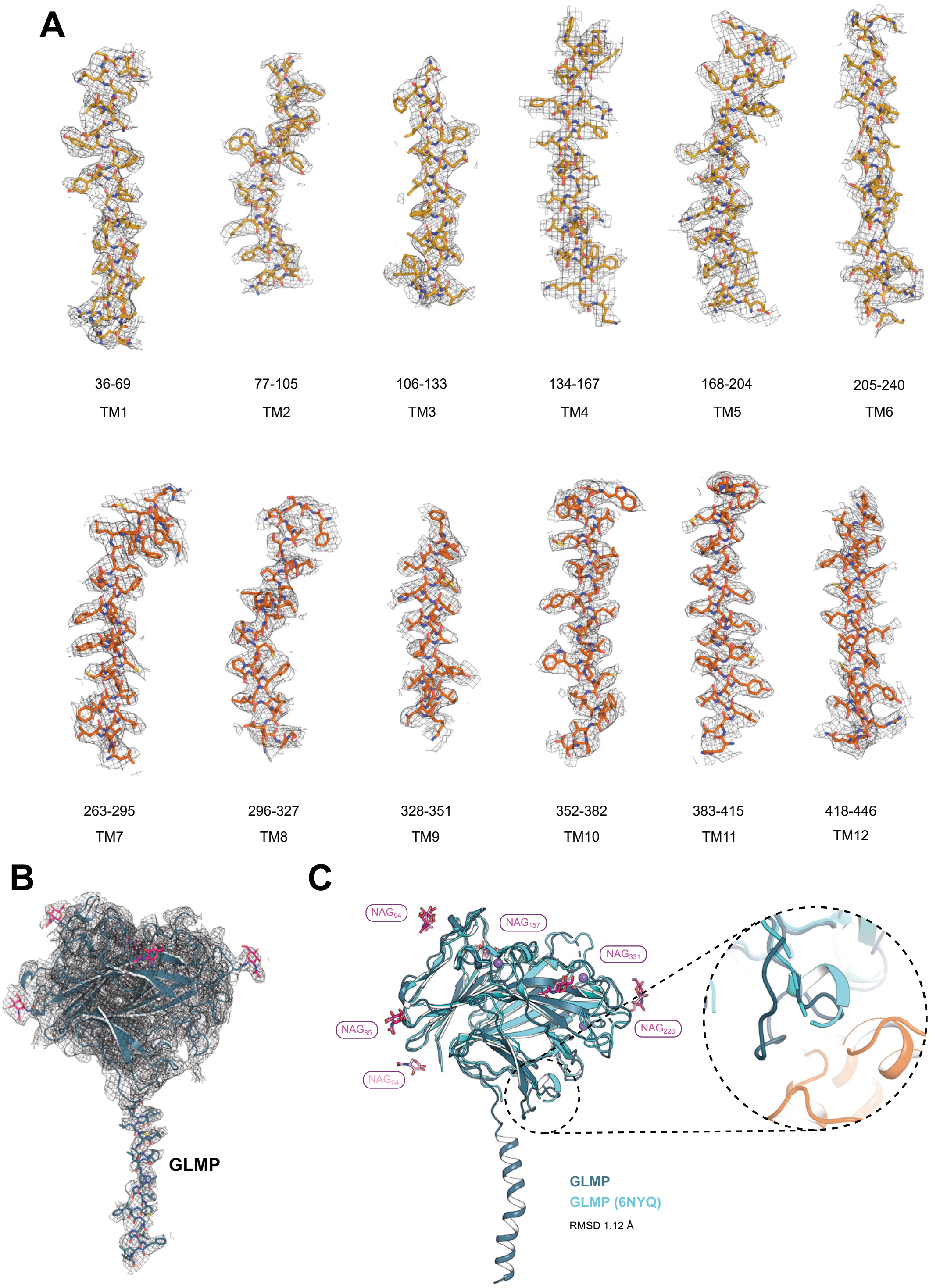


**Supplementary Figure 9. Density map of the Cryo-EM structure of GLMP-MFSD1$_{\text{His-Ala.}}$ (A)** Cyro-EM map (grey mesh) within a 2.5 Å radius around the MFSD1 model. Maps are shown for individual helices of MFSD1, with individual residues shown as sticks (yellow and orange). **(B)** Cryo-EM map (grey mesh) within a 2.5 Å radius around the model of GLMP (blue). Individual residues of the transmembrane helix and the five NAG molecules (pink) are shown as sticks. **(C)** Overlay of the Cryo-EM structure of GLMP (blue) with the X-ray structure of GLMP (light blue, PDB-ID: 6NYQ). The RMSD$_{C\alpha}$ of the superimposition is 1.12 Å over 271 residues of the luminal GLMP domain. Five of the NAG molecules (pink) identified in the Cryo-EM structure overlap with the six NAG molecules (light pink) found in the X-ray structure of GLMP (PDB-ID: 6NYQ). Additionally, the X-ray structure of GLMP contains three sodium ions (purple spheres). The zoom-in highlights the loop region, responsible for the interaction of GLMP with MFSD1, which has not been modeled in the crystal structure and is structured in the EM-derived model.

### Supplemental Figure 10

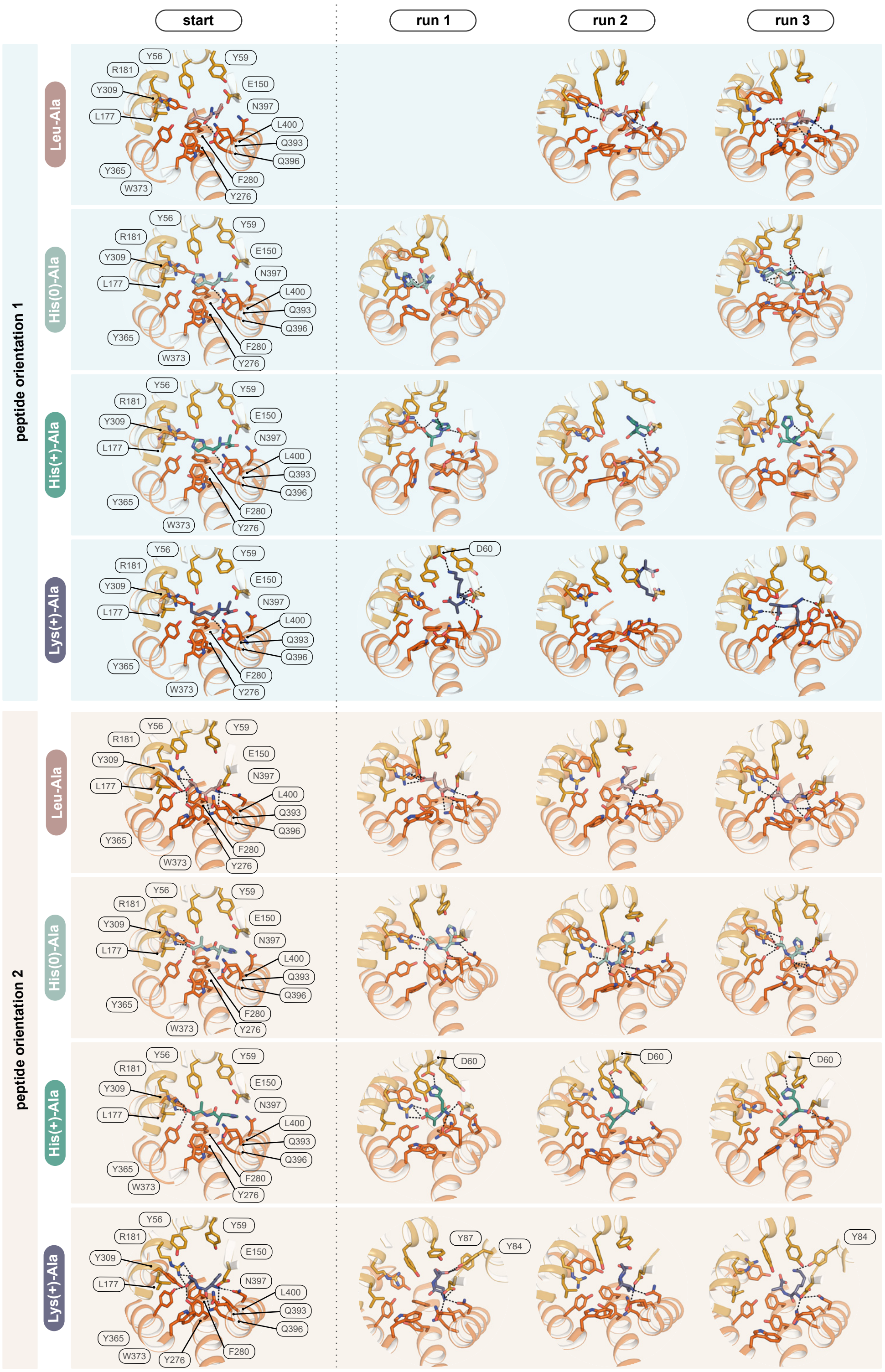


**Supplementary Figure 10. MD simulations of dipeptide-bound MFSD1.** MD simulations were performed on MFSD1 in complex with the dipeptides Leu-Ala (rose), His-Ala in its neutral (His(0)-Ala, pale teal) and charged (His(+)-Ala, teal) state and Lys(+)-Ala, purple). The basis of the binding mode for each peptide was the initial non-protein density found in the GLMP-MFSD1$_{\text{His-Ala}}$ map. The dipeptide His-Ala was placed in two different binding poses, denoted peptide orientation 1 (pale blue background) and peptide orientation 2 (light orange background). Based on this pose, the remaining ligands were oriented. Shown and labeled are critical binding site residues for each starting structure and the same view for the binding site of each simulation run after 300 ns. Additional interacting residues appearing at the endpoint of the simulation are highlighted in the respective panels.

### Supplemental Figure 11

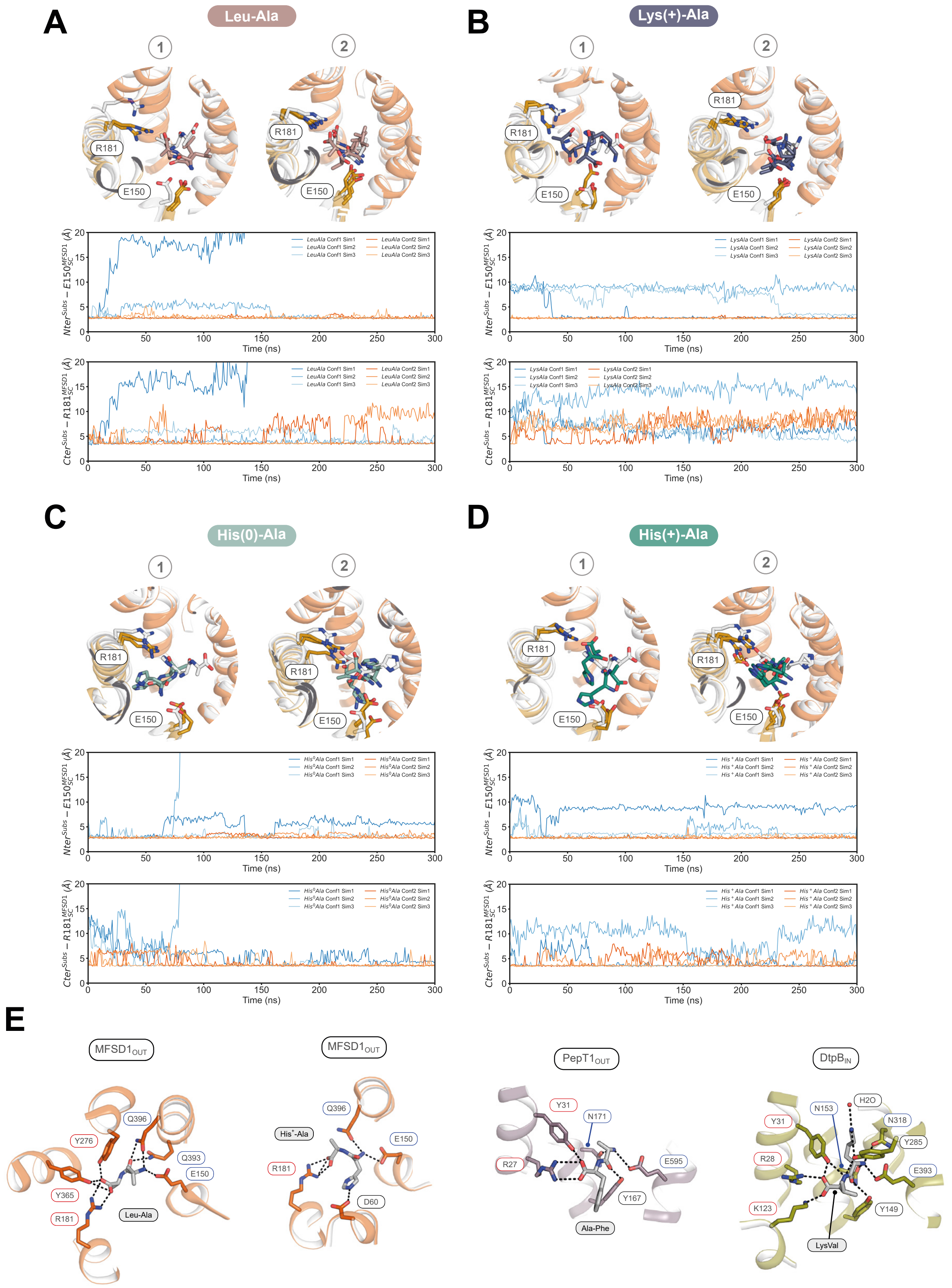


**Supplementary Figure 11. Flexibility of dipeptide binding in MFSD1 during MD simulations. (A)** The binding site of MFSD1 represents the starting pose of Leu-Ala (grey) and the final pose of the peptide after 300 ns of MD simulation (light purple) for each of the two peptide orientations (1 and 2). Below are RMSD plots of distant changes of the N- and C-terminus of the peptide with respect to residues E150 and R181. Plots show the results for each peptide orientation (orientation 1-blue, orientation 2-orange). MD simulations were run in triplicates. **(B)** Illustration of the MFSD1 binding site of MFSD1 with the starting pose of Lys-Ala (grey) and the final pose of the peptide after 300 ns of MD simulation (dark purple) for each of the two peptide orientations (1&2). Below are shown RMSD plots of distant changes of the N- and C-terminus of the substrate with respect to residues E150 and R181. Plots show the results for each peptide orientation (orientation 1-blue, orientation 2-orange). MD simulations were performed in triplicates. **(C)** Binding site of MFSD1 showing the starting pose of the dipeptide His(0)-Ala (grey) and the final pose after 300 ns of MD simulation (pale teal) for each of the two peptide orientations (1&2). Below are RMSD plots of distant changes of the N- and C-terminus of the substrate with respect to residues E150 and R181. RMSD plots highlight the results for each peptide orientation (orientation 1-blue, orientation 2-orange) run in triplicates. **(D)** The Starting pose of the dipeptide His(+)-Ala (grey) and the final pose in the MFSD1 binding site after 300 ns of MD simulation (dark teal) are shown for each of the two peptide orientations (1&2). Below are RMSD plots of distant changes of the N- and C-terminus of the substrate with respect to residues E150 and R181. MD simulations were run in triplicates for each peptide orientation (orientation 1-blue, orientation 2-orange). **(E)** Comparison of dipeptide binding sites of MFSD1 in the outward-open conformation (orange) either in complex with Leu-Ala (MD simulation run 3, peptide orientation 2) or His(+)-Ala (MD simulation run 1, peptide orientation 2), the Cryo-EM structure of PepT1 (PDB-ID: 7PMX) in the outward-open conformation (pale purple) in complex with Ala-Phe, and the X-ray structure of DtpB (PDB-ID: 8B1H) in the inward-open conformation (pale green) bound to the dipeptide Lys-Val. Critical residues important for the coordinating of the N-terminus of the substrate are framed in blue and the C-terminus in red. Hydrogen bonds are shown as black dashed lines.

### Supplemental Figure 12

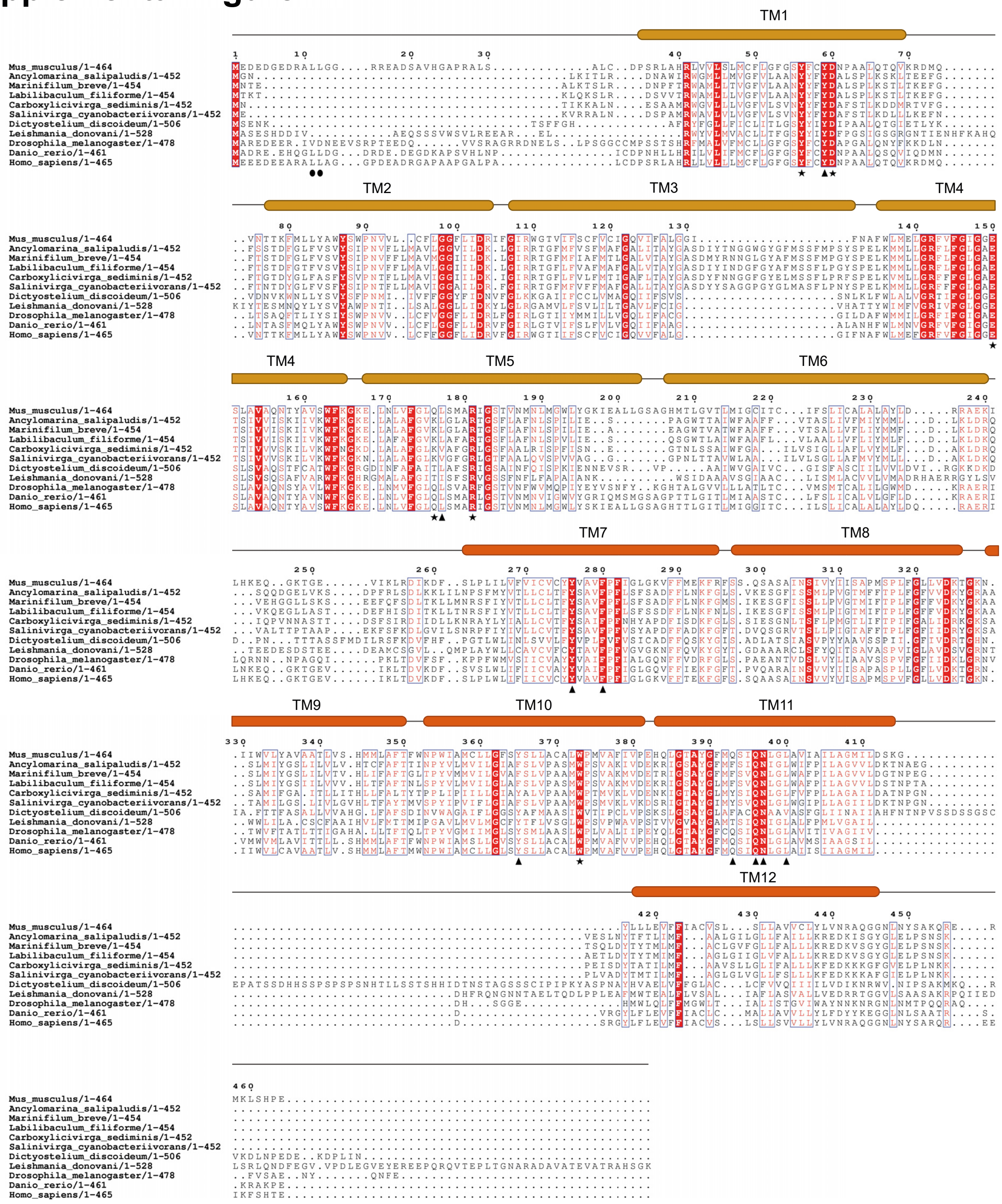


**Supplementary Figure 12. Multiple sequence alignment of MFSD1 with other MFSD1 homologs.** Sequence alignment of MFSD1 from *Mus musculus* (accession code: Q9DC37), *Ancylomarina salipaludis* (accession code: WP_129252052.1), *Marinifilum breve* (accession code: WP_165836040.1), *Labilibaculum filiforme* (accession code: WP_180335631.1), *Carboxylicivirga sediminis* (accession code: WP_212192738.1), *Salinivirga cyanobacteriivorans* (accession code: WP_057952039.1), *Dictyostelium discoideum* (accession code: XP_636334.1), *Leishmania donovani* (accession code: XP_003862116.1), *Drosophila melanogaster* (accession code: NP_001261505.1), *Danio rerio* (accession code: Q32LQ6) and *Homo sapiens* (accession code: Q9H3U5) using ClustalO. Conserved residues are highlighted in red letters, whereas highly conserved residues are shown with a red background. (●) dileucine motif, (▲) binding site residues, (★) binding site residues used for mutational studies. Above the sequence, the transmembrane helices are highlighted based on the determined structure, colored in dark yellow for the N-domain and orange for the C-domain.

### Supplemental Figure 13

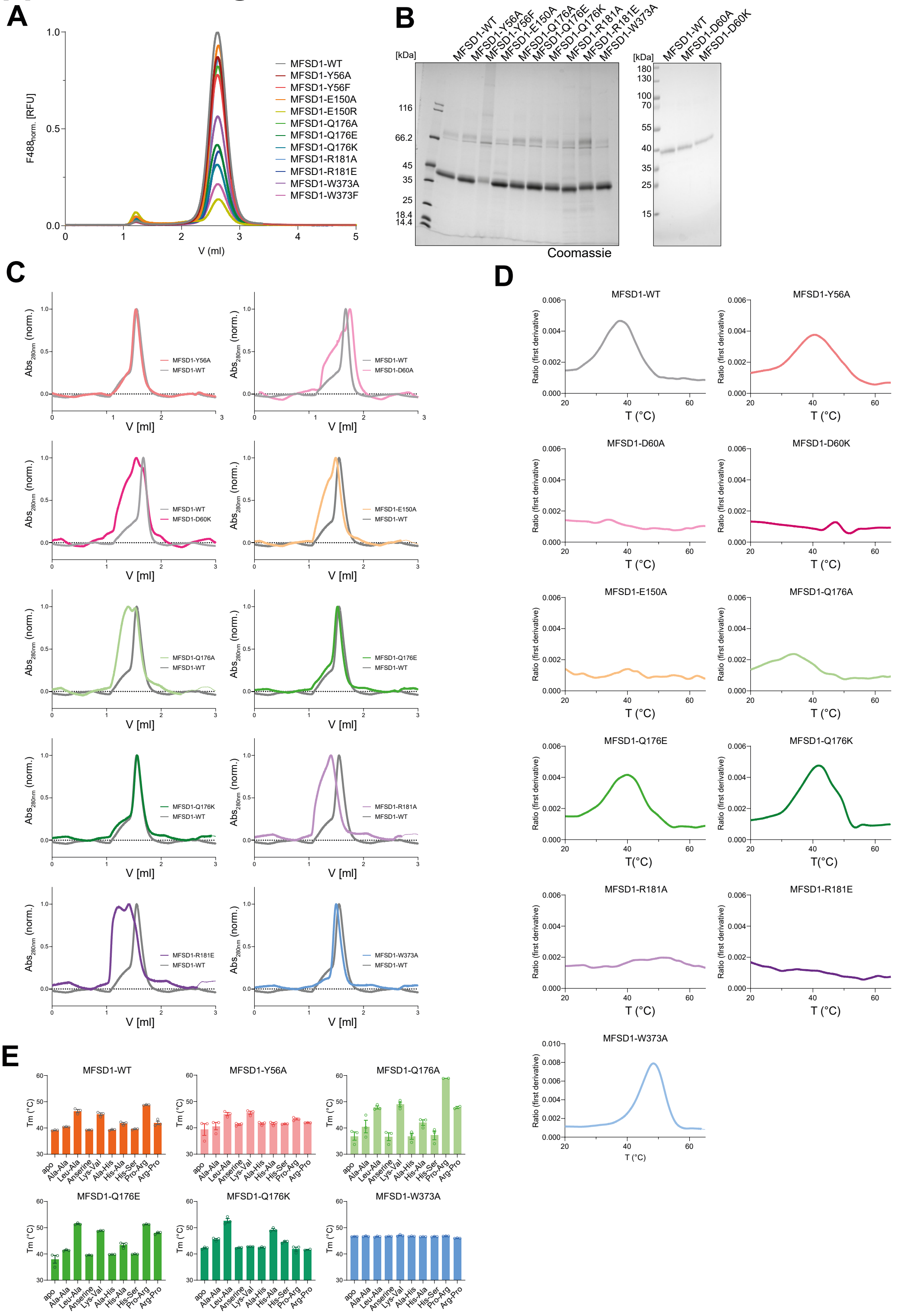


**Supplementary Figure 13. The binding-site mutations in MFSD1 lose binding to peptides in the thermal stability assay. (A)** FSEC analysis of $MFSD1_{WT}$ and mutants normalized to the fluorescent signal at $\lambda_{ex}$=488 nm/$\lambda_{em}$=510 nm of GFP (F488) of $MFSD1_{WT}$. The supernatant of soluble fraction after whole-cell solubilization was loaded onto a Superose 6 5/150 column. **(B)** SDS-PAGE of purified $MFSD1_{WT}$ and binding site mutants. For each lane, 2 µg of protein were loaded. **(C)**. Comparison of SEC traces of binding site mutants (colored) to $MFSD1_{WT}$ (grey) of each mutant. **(D), (E)** Melting temperatures derived from thermal stability experiments of each mutant in the absence (apo, grey) or presence of 5 mM of selected peptides. n=3 independent experiments with data shown as mean ± SD. For mutants for which no bar graph is given, unfolding traces are given for the apo state to show that no $T_M$ value could be determined.

### Supplemental Figure 14

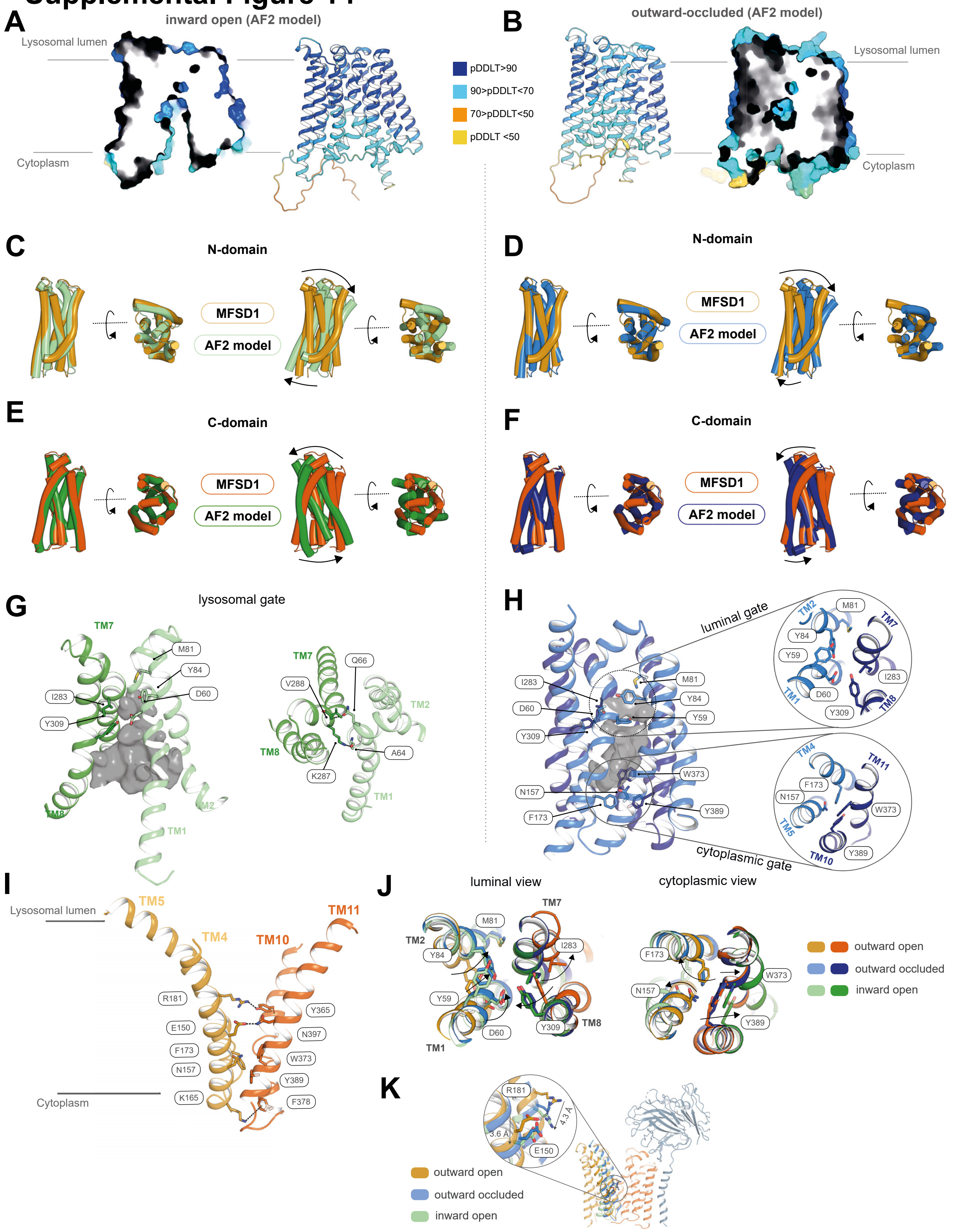


**Supplementary Figure 14. Comparison of Cryo-EM structure of MFSD1 to AlphaFold2 models in different conformations. (A)** AlphaFold2 (AF) prediction of MFSD1 in an inward open conformation colored by its pDDLT score. **(B)** AF model of the outward occluded conformation of MFSD1 colored by its pDDLT score. **(C)** Superposition of N-domains of MFSD1 Cryo-EM model (yellow) and inward-open AF model (pale green). The superpositions are shown either by aligning the N-domains only (left panel) or the entire model of MFSD1 (right panels). Observed changes are indicated by arrows. **(D)** Superposition of N-domains of MFSD1 Cryo-EM model (yellow) and the predicted outward-occluded AF model (pale blue). The superpositions represent the alignments of the N-domains only (left panel) or the complete model of MFSD1 (right panels). **(E)** Superposition of C-domains of MFSD1 cryoEM model (yellow) and inward-open AF model (forest green). The superpositions are shown either by aligning the N-domains (left panel) only or the entire model of MFSD1 (right panels). **(F)** Superposition of C-domains of MFSD1 Cryo-EM model (yellow) and outward-occluded AF model (sky blue). In the left panel, only the C-domains (left panel) were aligned, while the right panel represents the alignment of the entire transporter between the two states. **(G)** Illustration of the luminal gate of the inward-open AF model. Crucial residues involved in gate formation are indicated. **(H)** Luminal and cytoplasmic gates (including the close-up view) of outward-occluded AF model of MFSD1. Residues forming the gates are indicated. **(I)** Cytoplasmic gate and inter-bundle interactions of the outward-open state Cryo-EM model of MFSD1. **(J)** Cartoon representation of the conformational transitions of the gating residues of the outward-open Cryo-EM MFSD1 model and the two AF-predicted conformations viewed from the luminal and cytoplasmic side. The key residues forming the cytoplasmic and luminal gates are shown for each conformation. Color codes for the respective conformations are indicated. Arrows highlight the movement of each gating residue during the transition from inward to outward open. (**K**) Movement of residues E150 and R181, residing in the substrate-binding site within the N-domain (yellow) of MFSD1, during the transport cycle from outward-open (N-domain:yellow and C-domain:orange) to inward-open conformation (N-domain:green) highlighted with arrows to indicate the direction of the movement.

### Supplemental Figure 15

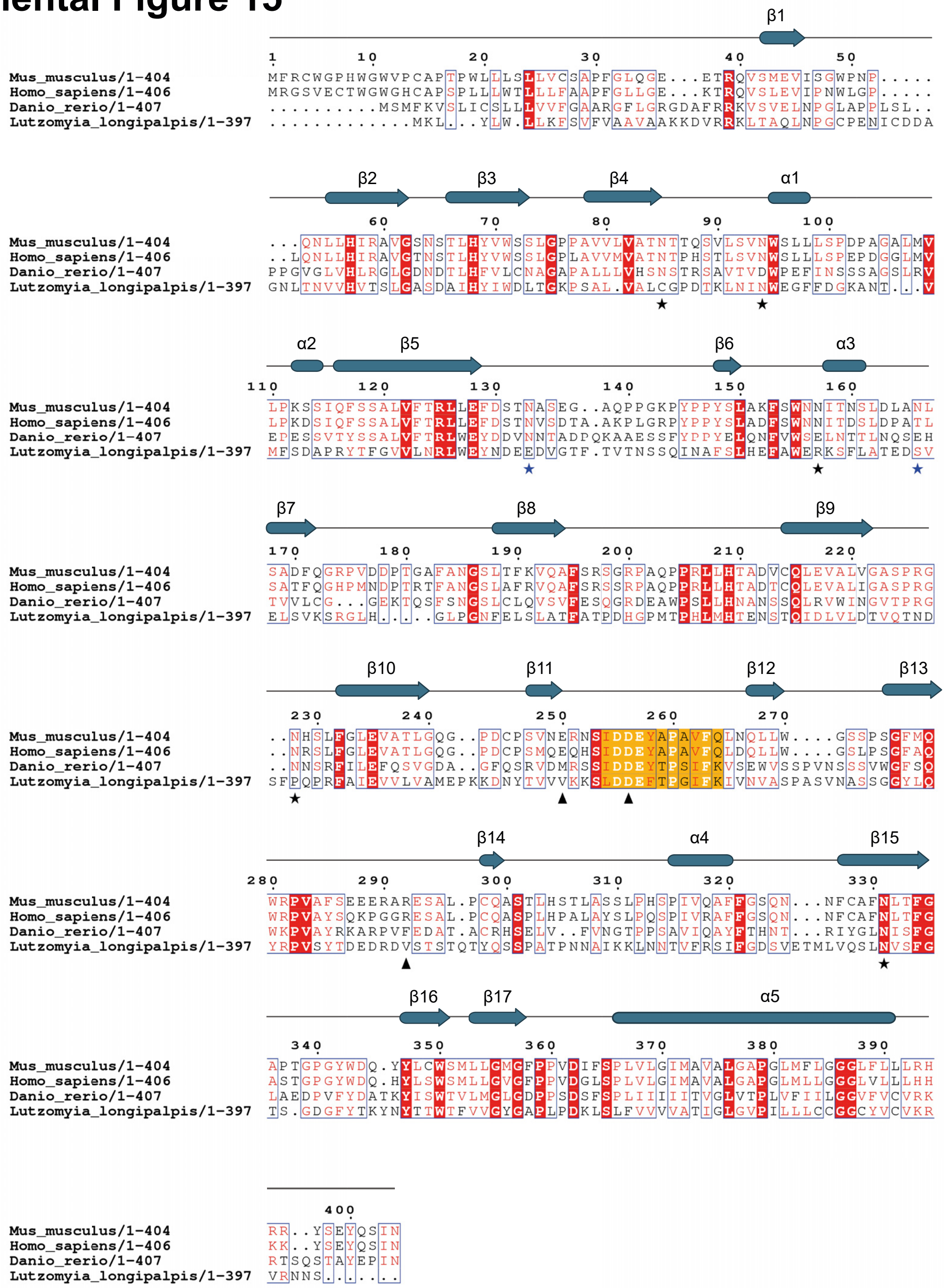


**Supplementary Figure 15. Multiple sequence alignment of GLMP with other GLMP homologs.** Sequence alignment of GLMP from *Mus musculus* (accession code: Q9JHJ3), *Danio rerio* (accession code: Q66HW4), *Lutzomyia longipalpis* (accession code: A0A7G3AQ10), and *Homo sapiens* (accession code: Q8WWB7) using ClustalO. Conserved residues are highlighted in red font, whereas highly conserved residues are shown with red background. Residues with a (★) represent identified glycosylation sites of GLMP in the Cryo-EM structure, whereas (★) label additional glycosylation sites as listed in UniProt. Residues highlighted with (▲) were mutated to test for the rescue of MFSD1 by GLMP. Colored in yellow is the loop region that was exchanged for Ala-Ala-Ala-Ala-Ala to test for MFSD1 rescue by GLMP reexpression in this study. Secondary structure elements of GLMP are shown above the alignment depicted by beta sheets (blue arrows) and alpha helices (blue tubes).

### Supplemental Figure 16

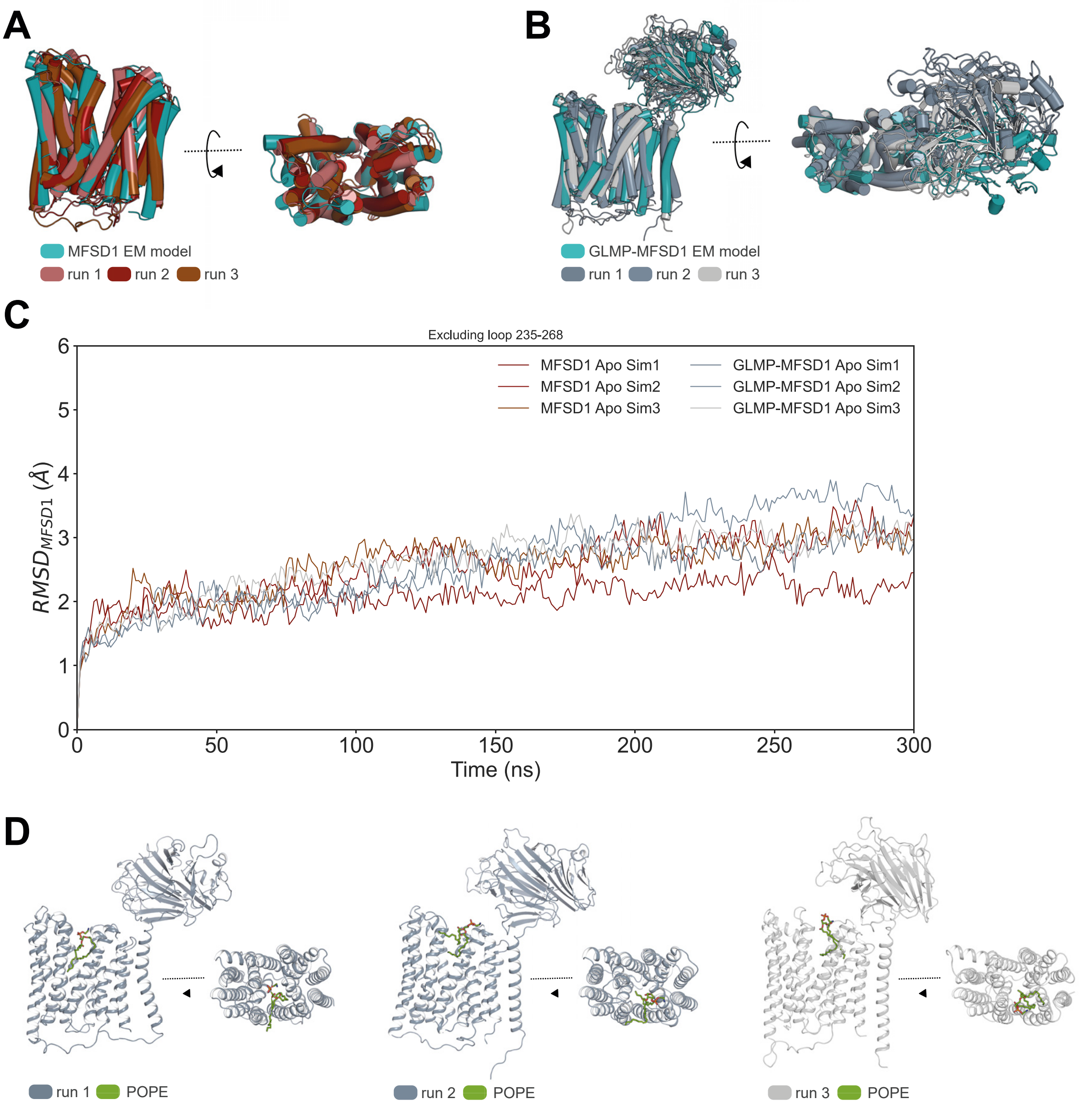


**Supplementary Figure 16. Analysis of MD simulations of GLMP-MFSD1$_{apo}$ and MFSD1$_{apo}$. (A)** Superposition of MFSD1 starting model (in blue, derived from the cryoEM model) and the structure after 300 ns of MD simulations of the three replicates (shades of red). **(B)** Superposition of GLMP-MFSD1 starting models (in blue, representing the cryoEM structure) and the complex after 300 ns of MD simulations of the three replicates (shades of grey). **(C)** RMSD (MFSD1 in relation to the starting CryoEM model) changes over the course of the MD simulation. The change in RMSD of MFSD1 in the apo form is shown in red, and the RMSD change of MFSD1 in the apo form as part of the complex with GLMP is given in grey blue. Each model was run in triplicates (Sim 1-3). **(D)** A POPE lipid molecule (green) is only found between TMs of MFSD1 during simulations (run1-3) of the GLMP-MFSD1 apo complex but not when simulations are run on MFSD1 only.

#### Supplemental Movie 1

**Movie S1. Conformational changes of MFSD1 during the transport cycle.** Video morphing between the apo outward-open (PDB: 8R8Q), apo occluded (AF2 model), and apo inward-open (AF2 model) conformations of MFSD1 (yellow) as viewed parallel to the membrane. Highlighted are residues involved in forming the cytoplasmic gate (dark blue) and the lysosomal gate (dark green), as well as residues forming salt bridges between the N- and C-terminal domain to stabilize the closing of the respective gates (light blue – cytoplasmic gate stabilization, light green – lysosomal gate stabilization).
